# Conserved chromatin and repetitive patterns reveal slow genome evolution in frogs

**DOI:** 10.1101/2021.10.18.464293

**Authors:** Jessen V. Bredeson, Austin B. Mudd, Sofia Medina-Ruiz, Therese Mitros, Owen K. Smith, Kelly E. Miller, Jessica B. Lyons, Sanjit S. Batra, Joseph Park, Kodiak C. Berkoff, Christopher Plott, Jane Grimwood, Jeremy Schmutz, Guadalupe Aguirre-Figueroa, Mustafa K. Khokha, Maura Lane, Isabelle Philipp, Mara Laslo, James Hanken, Gwenneg Kerdivel, Nicolas Buisine, Laurent M. Sachs, Daniel R. Buchholz, Taejoon Kwon, Heidi Smith-Parker, Marcos Gridi-Papp, Michael J. Ryan, Robert D. Denton, John H. Malone, John B. Wallingford, Aaron F. Straight, Rebecca Heald, Dirk Hockemeyer, Richard M. Harland, Daniel S. Rokhsar

## Abstract

Frogs are an ecologically diverse and phylogenetically ancient group of living amphibians that include important vertebrate cell and developmental model systems, notably the genus *Xenopus*. Here we report a high-quality reference genome sequence for the western clawed frog, *Xenopus tropicalis*, along with draft chromosome-scale sequences of three distantly related emerging model frog species, *Eleutherodactylus coqui*, *Engystomops pustulosus* and *Hymenochirus boettgeri*. Frog chromosomes have remained remarkably stable since the Mesozoic Era, with limited Robertsonian (i.e., centric) translocations and end-to-end fusions found among the smaller chromosomes. Conservation of synteny includes conservation of centromere locations, marked by centromeric tandem repeats associated with Cenp-a binding, surrounded by pericentromeric LINE/L1 elements. We explored chromosome structure across frogs, using a dense meiotic linkage map for *X. tropicalis* and chromatin conformation capture (HiC) data for all species. Abundant satellite repeats occupy the unusually long (∼20 megabase) terminal regions of each chromosome that coincide with high rates of recombination. Both embryonic and differentiated cells show reproducible association of centromeric chromatin, and of telomeres, reflecting a Rabl configuration similar to the “bouquet” structure of meiotic cells. Our comparative analyses reveal 13 conserved ancestral anuran chromosomes from which contemporary frog genomes were constructed.

## Introduction

Amphibians are widely used models in developmental and cell biology^1–5^, and their importance extends to the fields of infectious disease, ecology, pharmacology, environmental health, and biological diversity^6–10^. While the principal model systems belong to the genus *Xenopus* (notably the diploid western clawed frog *X. tropicalis* and the paleo-allotetraploid African clawed frog *X. laevis*, other amphibian models have increasingly been introduced due to their diverse developmental, cell biological, physiological, and behavioral adaptations^11–14^.

While genome evolution has been extensively studied in mammals^15^ and birds^16,17^, the relative lack of phylogenetically diverse chromosome-scale frog genomes has limited the study of genome evolution in anuran amphibians. Here, we report a high-quality assembly for *X. tropicalis* and three new chromosome-scale genome assemblies for the direct-developing Puerto Rican coquí (*Eleutherodactylus coqui*), the túngara frog (*Engystomops pustulosus*), which is a model for vocalization, and the Zaire dwarf clawed frog (*Hymenochirus boettgeri*), which has an unusually small embryo and is a model for regulation of cell and body sizes. Genome assemblies are essential resources for further work to exploit the experimental possibilities of these diverse animals. The new high quality *X. tropicalis* genome upgrades previous draft assemblies^18,19^ and our new genomes complement draft chromosome-scale sequences for the African clawed frog^20^ (*Xenopus laevis*), the African bullfrog^21^ (*Pyxicephalus adspersus*), the Leishan moustache toad^22^ (*Leptobrachium leishanense*), the Ailao moustache toad^23^ (*Leptobrachium* [*Vibrissaphora*] *ailaonicum*), and Asiatic toad^24^ (*Bufo gargarizans*), as well as scaffold- and contig-scale assemblies for other species^25^. The rapidly increasing number of chromosome-scale genome assemblies makes anurans ripe for comparative genomic and evolutionary analysis.

Chromosome number variation among frogs is limited^26^. Based on cytological^27,28^ and sequence comparisons^18,20,26,29,30^ most frogs have *n* ∼ 10–12 pairs of chromosomes. The constancy of the frog karyotype is in contrast with the more dramatic variation seen across mammals^15,31^, which as a group is considerably younger than frogs. The constancy of the frog karyotype parallels the static karyotypes of birds^16^, although birds typically have nearly three times more chromosomes than frogs, including numerous microchromosomes (among frogs, only the basal *Ascaphus*^32^ has microchromosomes). Despite the stable frog chromosome number, however, fusions, fissions, and other inter-chromosomal rearrangements do occur, and we can use comparisons among chromosome-scale genome sequences to (1) infer the ancestral chromosomal elements, (2) determine the rearrangements that have occurred along frog phylogeny, and (3) characterize the patterns of chromosomal change among frogs. These findings of conserved synteny among frogs are consistent with prior demonstrations of conservation between *Xenopus tropicalis* with other tetrapods, including human and chicken^18,33^.

Since frog karyotypes are so highly conserved, *X. tropicalis* can be used as a model for studying chromosome structure, chromatin interaction, and recombination for the entire clade. Features that can be illuminated at the sequence level include the structure and organization of centromeres and nature of the unusually long subtelomeres relative to mammals (frog subtelomeres are ∼20 megabases, compared with the mammalian subtelomeres that are typically shorter than a megabase). The extended subtelomeres of frogs form interacting chromatin structures in interphase nuclei that reflect three-dimensional intra-chromosome and inter-chromosome subtelomeric contacts, which are consistent with a Rabl configuration. As in other animals, subtelomeres of frogs have an elevated GC content and recombination rate. Here we show that the unusually high enrichment of recombination in the subtelomeres likely reflects similar structural and functional properties in other vertebrates, though the quality of the assembly reveals that the length of subtelomeres, enrichment of transposon subsequences by unequal crossing over, and high recombination rates are considerably greater than in mammals. We use Cenp-a binding at satellites to confirm centromere identity and extend the predictive power of the repeat structures to centromeres of other frogs. We address the unusually high recombination rate in subtelomeric regions, correlating with the landscape of base composition and transposons. Over the 200 million years of evolution that we address here, centromeres have generally been stable, but the few karyotypic changes reveal the predominant Robertsonian translocations at centromeric regions; we also document the slow degeneration that occurs to inactivated centromeres and fused telomeres, changes that are obscured in animals with rapidly evolving karyotypes.

## Results

### New frog genome sequences

#### High-quality chromosome-scale genome assembly for *X. tropicalis*

To establish a high-quality chromosomal reference genome sequence to study the structure and organization of *Xenopus tropicalis* chromosomes and for comparisons with other frog genomes, we integrated multiple sequencing technologies, including Single-Molecule Real-Time long reads (SMRT sequencing; Pacific Biosciences), linked read sets (10x Genomics), short-read shotgun sequencing, *in vivo* chromatin conformation capture, and meiotic mapping, combined with previously generated dideoxy shotgun sequence (**Supplementary Data 1, Supplementary Figs. 1A–D** and **2**, and **Supplementary Notes 1** and **2**). New sequences were generated from 17th generation individuals from the same inbred Nigerian line that was used in the original Sanger shotgun sequencing^33^. The completeness, protein coding capacity, repeat structure and sequence variation are discussed in supplementary information (**Supplementary Figs. 1–6** and **Supplementary Tables 1–4)** providing the basis for comparisons.

The new v10 reference assembly spans 1,448.4 Mb and is substantially more complete than the previous v9 sequence^18^, assigning 219.2 Mb more sequence to chromosomes (**Supplementary Table 1**). The v10 assembly is also far more contiguous, with half of the sequence contained in 32 contigs longer than 14.6 Mb (vs. 71.0 kb in v9). The assembly captures 99.6% of known coding sequence (**Supplementary Table 2, Supplementary Note 2**). We found that the fragmented quality of earlier assemblies was due in part to the fact that 68.3 Mb (4.71%) of the genome was not sampled by the 8× redundant Sanger dideoxy whole-genome shotgun dataset^33^ (**Supplementary Fig. 3, Supplementary Note 2**). These missing sequences, apparently due to non-uniformities in shotgun cloning and/or sequences (**Supplementary Fig. 1E**), are distributed across an estimated 140.5k blocks of mean size 485.7 bp (longest 50.0 kb) on the new reference assembly and capture an additional 6,774 protein-coding exons (**Supplementary Fig. 1F–G**). The enhanced contiguity of v10 is accounted for by the relatively uniform coverage of PacBio long-read sequences on the genome, as expected from other studies^34–37^. Most remaining gaps are in highly repetitive and satellite-rich centromeres and subtelomeric regions (see below) (**Supplementary Figs. 1H and 3**).

#### Additional chromosome-scale frog genomes

To assess the evolution of chromosome structure across a diverse set of frogs, we generated chromosome-scale genome assemblies for three new emerging model species, including the Zaire dwarf clawed frog *Hymenochirus boettgeri* (a member of the family Pipidae along with *Xenopus* spp.) and two neobratrachians: the Puerto Rican coquí *Eleutherodactylus coqui* (family Eleutherodactylidae) and the túngara frog *Engystomops pustulosus* (family Leptodactylidae). These chromosome-scale draft genomes were primarily assembled from short-read datasets and chromatin conformation capture (HiC) data (**Supplementary Data 1, Supplementary Table 5, Supplementary Note 3**). To further expand the scope of our comparisons, we also updated the assemblies of two recently published frog genomes: the African bullfrog *Pyxicephalus adspersus*^21^ from the neobatrachian family Pyxicephalidae, and the Ailao moustache toad *Leptobrachium* (*Vibrissaphora*) *ailaonicum*^22^ from the family Megophryidae (**Supplementary Fig. 7, Supplementary Note 3**). These species span the pipanuran clade, which comprises all extant frogs except for a small number of phylogenetically basal taxa, such as *Ascaphus*^38^.

The chromosome numbers of the new assemblies agree with previously described karyotypes for *E. coqui*^39^ (2*n* = 26) and *E. pustulosus*^40^ (2*n* = 22). The literature for *H. boettgeri,* however, is more equivocal, with reports^41,42^ of 2*n* = 20–24. The *n* = 9 chromosomes in our *H. boettgeri* assembly are consistent with our chromosome spreads (**Supplementary Fig. 7A**). The karyotype variability in the published literature and discrepancy with karyotypes of our *H. boettgeri* samples may be the result of cryptic sub-populations within this species, or segregating chromosome polymorphisms.

#### Protein-coding gene set for *X. tropicalis*

The improved *X. tropicalis* genome encodes an estimated 25,016 protein-coding genes (**Supplementary Table 3**), which we predicted by taking advantage of 8,580 full-length-insert *X. tropicalis* cDNAs from the “Mammalian” Gene Collection^43^ (MGC), 1.27 million Sanger-sequenced expressed sequence tags^33^ (ESTs), and 334.5 Gbp of RNA-seq data from an aggregate of 16 conditions and tissues^44,45^ (**Supplementary Data 1, Supplementary Note 2**). The predicted gene set is a notable improvement on previous annotations, both in completeness and in full-length gene-level accuracy, due in part to the more complete assembly (**Supplementary Fig. 1, Supplementary Table 2, Supplementary Note 2**). In particular, gaps in the earlier genome assemblies arising from cloning biases in the Sanger sequencing process and encompassing exons embedded in highly repetitive sequences have been filled by single molecule long reads (**Supplementary Figures 1 and 3**).

A measure of this completeness and the utility of the *X. tropicalis* genome is provided by comparing its gene set with those of vertebrate model systems with reference-quality genomes, including chicken^46^, zebrafish^47^, mouse^48^ and human^49,50^ (**Supplementary Fig. 4**). Notably, despite the closer phylogenetic relationship between birds and mammals, *X. tropicalis* shares more orthologous gene families (and mutual best hits) with human than does chicken, possibly because of the loss of genomic segments in the bird lineage^16,51^ and/or residual incompleteness of the chicken reference sequence, due to the absence of several microchromosomes^46^. For example, of 13,009 vertebrate gene families with representation from at least four of the vertebrate reference species, only 341 are missing from *X. tropicalis* versus 1,110 from chicken (**Supplementary Fig. 4**). The current *X. tropicalis* genome assembly also resolves gene order and completeness of gene structures in the long subtelomeres that were missed in previous assemblies due to their highly repetitive nature (**Supplementary Fig. 1F–G**).

#### Protein-coding gene sets for other frogs

We annotated the new genomes of *E. coqui*, *E. pustulosus*, *H. boettgeri,* and *P. adspersus* using transcriptome data from these species (**Supplementary Data 1**) and peptide homology with *X. tropicalis* (**Supplementary Tables 6 and 7**). To include mustache toad in our cross-frog comparisons we adopted the published annotation of Li et al.^22^ (**Supplementary Note 3**). We found 14,412 orthologous groups across the five genera with OrthoVenn2^52^, including genes found in at least four of the five frog genera represented (**Supplementary Fig. 7B**). As expected, due to its reference-quality genome and well-studied transcriptome, only 72 of these clusters were not represented in *X. tropicalis*; the other frog genomes each had between 575 and 712 of these genes missing (or mis-clustered), suggesting better than 95% completeness in the other species. For analyses of synteny, we further restricted our attention to 7,292 one-to-one gene orthologs that were present on chromosomes (as opposed to unlinked scaffolds) in the “core” genomes *X. tropicalis, H. boettgeri, E. coqui*, *E. pustulosus*, and *P. adspersus*. The total branch length in the pipanuran tree shown in **Fig. 1** (including both *X. laevis* subgenomes) is 2.58 substitutions per four-fold synonymous site.

**Fig. 1.**
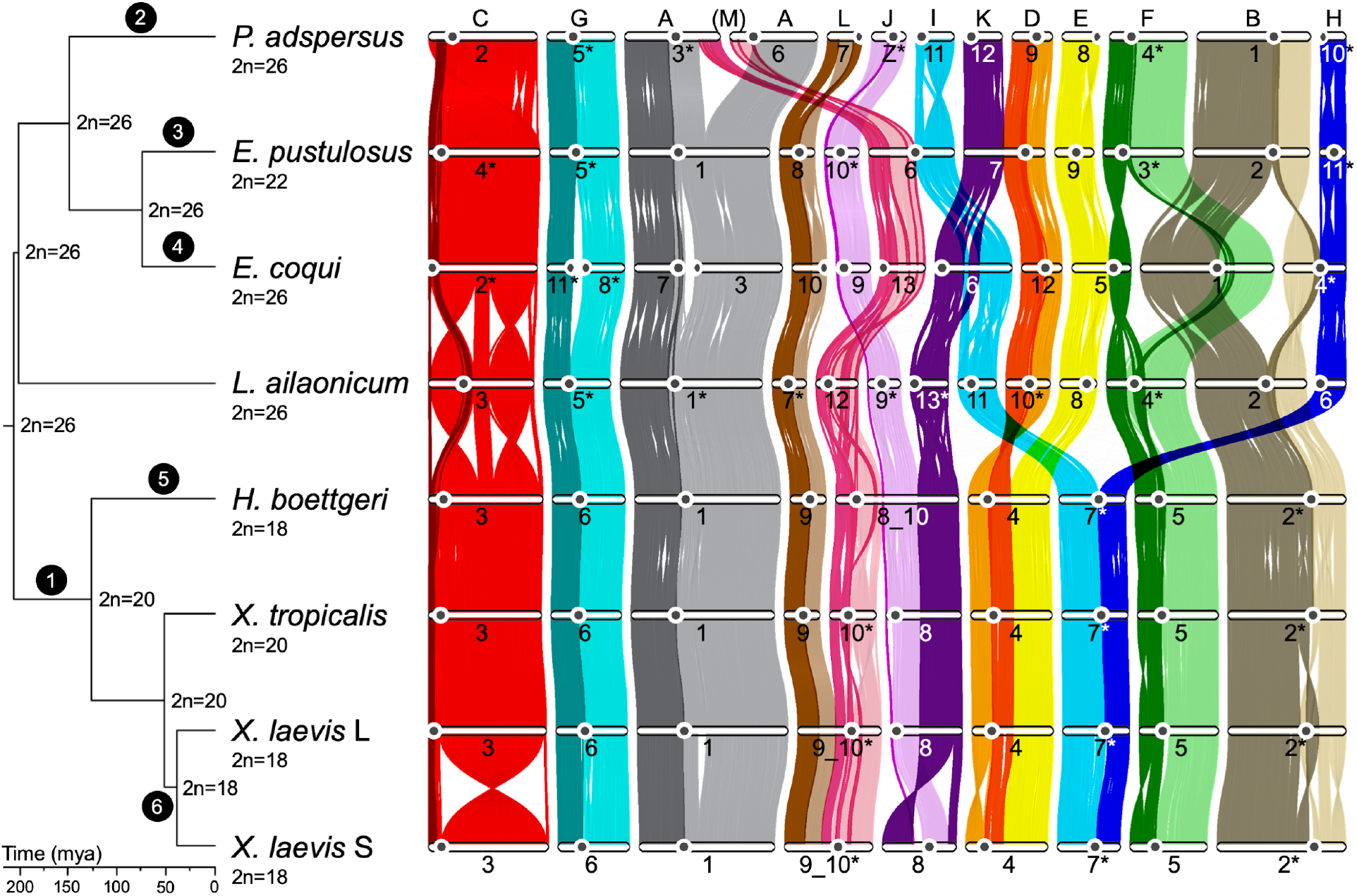
Phylogenetic tree and gene ortholog alignment. The phylogenetic tree of the seven analyzed species, calculated from fourfold degenerate sites and divergence time confidence intervals, was visualized with FigTree (commit 901211e; https://github.com/rambaut/figtree). The ancestral karyotype at each node was labeled on the tree. The alignment plot was generated with jcvi.graphics.karyotype^168^ (v0.8.12; https://github.com/tanghaibao/jcvi) using the 7,292 described chromosome one-to-one gene orthologs from OrthoVenn2 (ref.^52^), followed by visual filtering of single stray orthologs. The pericentromeric region based on HiC inference was represented with a black circle on each chromosome. The ancestral chromosomes (A to M) were labeled at the top of the alignment based on the corresponding region in *P. adspersus*. The alignments for each ancestral chromosome were colored uniquely, with those upstream and downstream of the *X. tropicalis* centromeric satellite repeat from tandem repeat analysis shaded with a light versus dark shade of the ancestral chromosome color. Chromosomes labeled with an asterisk were reverse complemented in this image relative to the orientation in the assembly. Black circles with white text reference chromosome changes outlined in **Table 1**.

### Repetitive landscape

Centromeric and telomeric tandem repeats play a critical role in the stability of chromosome structure^53^. Nonetheless, other kinds of repeats also play a role in the preservation of these important chromosome landmarks^54,55^. The new *X. tropicalis* v10 assembly captures sequences from centromeres and distal sub-telomeres that were fragmented in the previous assemblies^18,33^. The percentage of the genome covered by transposable elements is slightly higher than previously reported^33^ (36.82% vs. 34%) (**Supplementary Table 4**).

Insertional bias in the pericentromeric regions is observed for specific families of long interspersed elements (LINEs), including the relatively young Chicken Repeat 1 (CR1, ref.^56^) (3.14% of the genome) and the ancient L1 (1.06%) (**Fig. 2** and **Supplementary Fig. 5**). The *X. tropicalis* v10 assembly captures significantly more tandem repeats in the distal subtelomeric portions of the genome relative to earlier assemblies. An exhaustive search for tandem repeats using Tandem Repeat Finder^57^ determined that 10.67% of the chromosomes is covered by tandem arrays consisting of 5 or more monomeric units greater than 10 bp. Many tandem repeat footprints are in gaps from previous assemblies^18,33^ (**Supplementary Fig. 3**). Our new hybrid genome assembly closed many gaps containing centromeric and subtelomeric tandem repeats and captured numerous subtelomeric genes (**Supplementary Fig. 1**). The overall repeat landscape derived from the *X. tropicalis* assembly is mirrored in the other frog assemblies, with similar centromeric repeats, and lengthy subtelomeres, as discussed below (**Supplementary Fig. 9**).

**Fig. 2.**
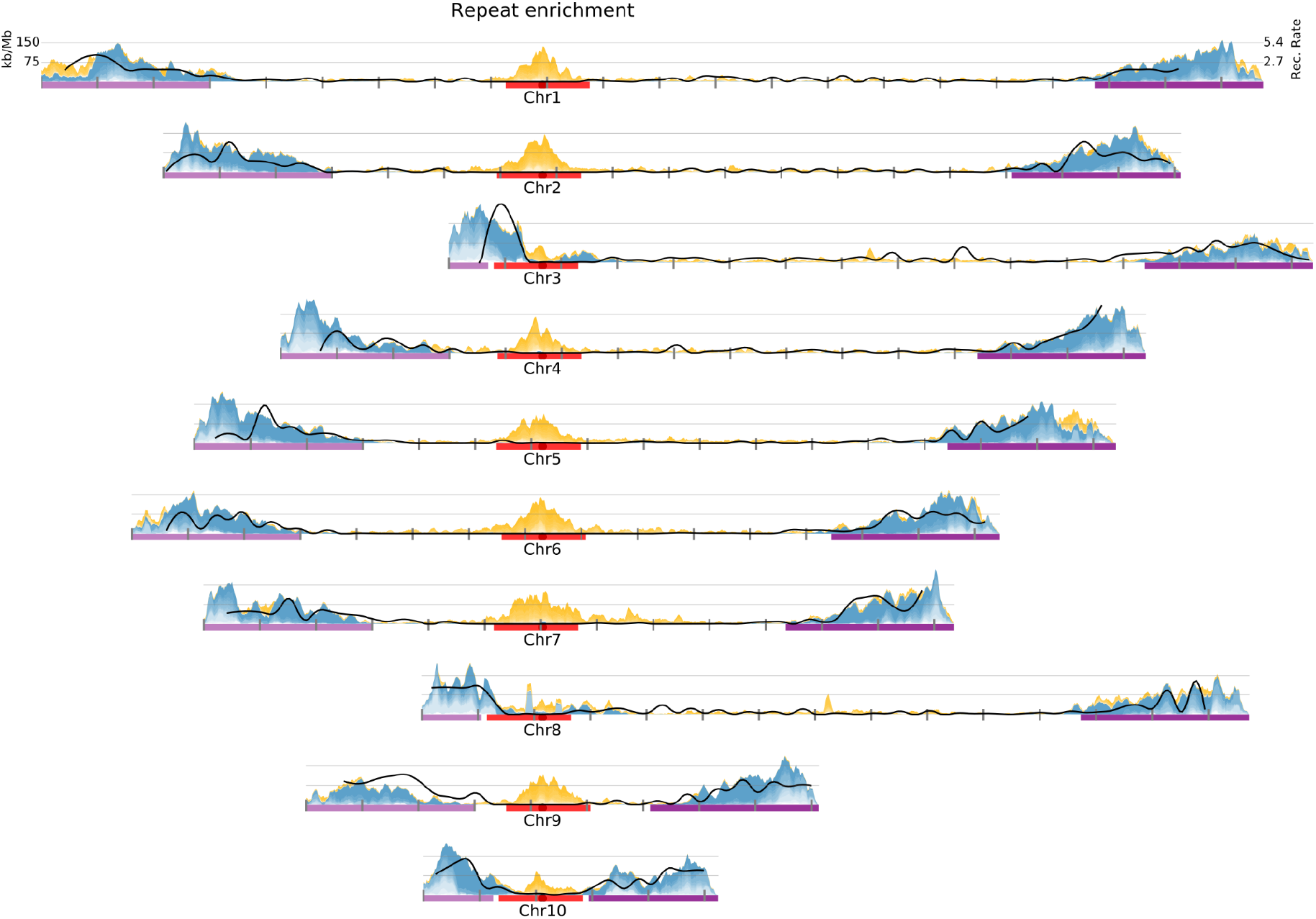
Density of pericentromeric and subtelomeric repeats in *Xenopus tropicalis*. Pericentromeric boundaries (red) and subtelomeric boundaries (purple) were used to obtain enriched repeats excluding chromosomes with short p-arms (chromosomes 3, 8, and 10). Pericentromeric repeats (yellow) correspond to selected subsets of non-LTR retrotransposons (CR1, L1, and Penelope), LTR retrotransposons (Ty3), and DNA transposons (PiggyBac and Harbinger). Subtelomeric enriched repeats (blue) correspond mainly to Satellite repeats and LTR retrotransposons (Ty3, Ngaro). Chromosomes are centered by the position of centromeric tandem repeats (black dot and dotted vertical line). The rate of recombination (cM/Mb) is shown as a solid black line. Tick marks indicate 10 Mb blocks (**Supplementary Fig. 16**).

### Genetic variation

Although the *X. tropicalis* reference genotype is highly inbred, it nevertheless retains 15 long heterozygous blocks ranging in size from 1.34 to 74.6 Mb. This exceeds the expectations based on 17 generations of brother-sister mating, suggesting residual heterozygosity could be maintained by balancing selection. Within the heterozygous blocks we observe 3.0 single nucleotide variants per kilobase. To begin to develop a catalog of segregating variation we sequenced pools of frogs from the Nigerian and Ivory Coast B populations, which have been previously analysed using SSLP markers^58^. From our light pool shotgun analysis we identified a total of 6,546,379 SNPs. There were 2,482,703 variants in the Nigerian pool and 4,661,928 in the Ivory Coast B pool, with 598,252 shared by both pools, pointing to substantial differentiation between populations (**Supplementary Fig. 6, Supplementary Note 2**).

### Evolutionary dynamics of frog chromosomes

#### Conserved synteny and ancestral chromosomes

Comparison of the chromosomal positions of orthologs across seven frog genomes reveals extensive conservation of synteny and collinearity (**Fig. 1**, **Supplementary Fig. 8**). We identified 13 conserved pipanuran syntenic units that we denote A through M (**Methods**, **Supplementary Note 4**). Each unit likely represents an ancestral pipanuran chromosome, an observation consistent with the 2*n* = 26 ancestral karyotype inferred from cytogenetic comparisons across frogs^27,59^. Over 95% (6,952 of 7,292) of chromosomal one-to-one gene orthologs are maintained in the same unit across the five frog species, attesting to the stability of these chromosomal elements (**Fig. 1**). The conservation of gene content per element is comparable to the 95% ortholog maintenance in the Muller elements in *Drosophila* spp.^60^. Despite an over two-fold difference in total genome size across the sampled genomes, each ancestral pipanuran element accounts for a nearly constant proportion of the total genome size, gene count, and repeat count in each species, implying uniform expansions and contractions during the history of the clade (**Supplementary Fig. 7C**).

At least some of these pipanuran elements have a deeper ancestry within amphibians. Comparison with the genome of the axolotl, *Ambystoma mexicanum*—a member of the order Caudata (salamanders and newts), and ∼292 million years divergent from pipanurans^61^—reveals conservation of multiple syntenic units (**Supplementary Fig. 8A**). For example, axolotl chromosomes 4, 6, 7, and 14 are in near 1:1 correspondence with pipanuran elements F, A, B and K, respectively, although small pieces of F and A can be found on axolotl 10, and parts of B can be found on axolotl 9 and 13. Other axolotl chromosomes are fusions of parts of two or more pipanuran elements. For example, axolotl chromosome 5 is a fusion of a portion of J with most of G; the remainder of G is fused with a portion of L on the q arm of axolotl 2. Further comparisons are needed to determine which of these rearrangements occurred on the axolotl vs. the stem pipanuran lineage. Genomes from the superfamilies Leiopelmatoidea and Alytoidea, which diverged prior to the radiation of pipanurans, will also be informative.

#### Chromosome evolution

Block rearrangements of the 13 ancestral elements dominate the evolutionary dynamics of pipanuran karyotypes (**Table 1**, **Fig. 1**). While element C has remained intact as a single chromosome across the group (except for internal inversions), all the other elements have experienced translocations during pipanuran evolution. During these translocations, the elements have remained intact with the exception of the breakage of elements A and M by reciprocal partial arm exchange observed in *P. adspersus* chromosomes 3 and 6.

**Table 1.** Organization and conservation of the 13 ancestral chromosomes of pipanuran genomes.

|  | Phylogenetic position | Structural event |
| --- | --- | --- |
| (1) | Stem pipid lineage | $J + K \Rightarrow JK$<br>$D. + E. \Rightarrow D.E$<br>$I \bullet + \bullet H \Rightarrow I \bullet H$ (Rob. fusion) |
| (2) | *P. adspersus* lineage after divergence from *R. temporaria* | $A + M \Rightarrow A1.m1 + m2.A2$ |
| (3) | *E. pustulosus* lineage after divergence from *E. coqui* | $M + I \Rightarrow M.I$ (Rob)<br>$K + D \Rightarrow K.D$ (Possible end-end) |
| (4) | *E. coqui* lineage after divergence from *E. pustulosus* | $G1 \bullet G2 \Rightarrow G1 \bullet + \bullet G2$ (Rob. fission)<br>$A1 \bullet A2 \Rightarrow A1 \bullet + \bullet A2$ (Rob. fission)<br>$I + K \Rightarrow I \bullet K$ (Rob. fusion + inversion)<br>$E + F1 \bullet F2 + B1 \bullet B2 + H \Rightarrow E \bullet F1 + F2 \bullet B2 + B1 \bullet H$ |
| (5) | *H. boettgeri* lineage after divergence from *Xenopus* | $M + J \bullet K \Rightarrow MJK$ |
| (6) | *X. laevis* progenitor lineage after divergence from *X. tropicalis* | $L + M \Rightarrow LM$ |

Rob, Robertsonian. Middle-dots (i.e., "•") represent centromeres. Periods (i.e., ".") represent translocation breakpoints.

To trace the evolutionary history of centromeres shown in **Fig. 1**, we inferred their positions using HiC contact map patterns as in *X. tropicalis* (where centromeres were also confirmed by analysis of Cenp-a binding as described below). In general, the pericentromeres of other pipanurans were characterized by the same repetitive element families found in *Xenopus*, further corroborating their identification. Overall, we found broad pericentromeric conservation among the species analyzed (**Figs. 1** and **3A**).

**Fig. 3.**
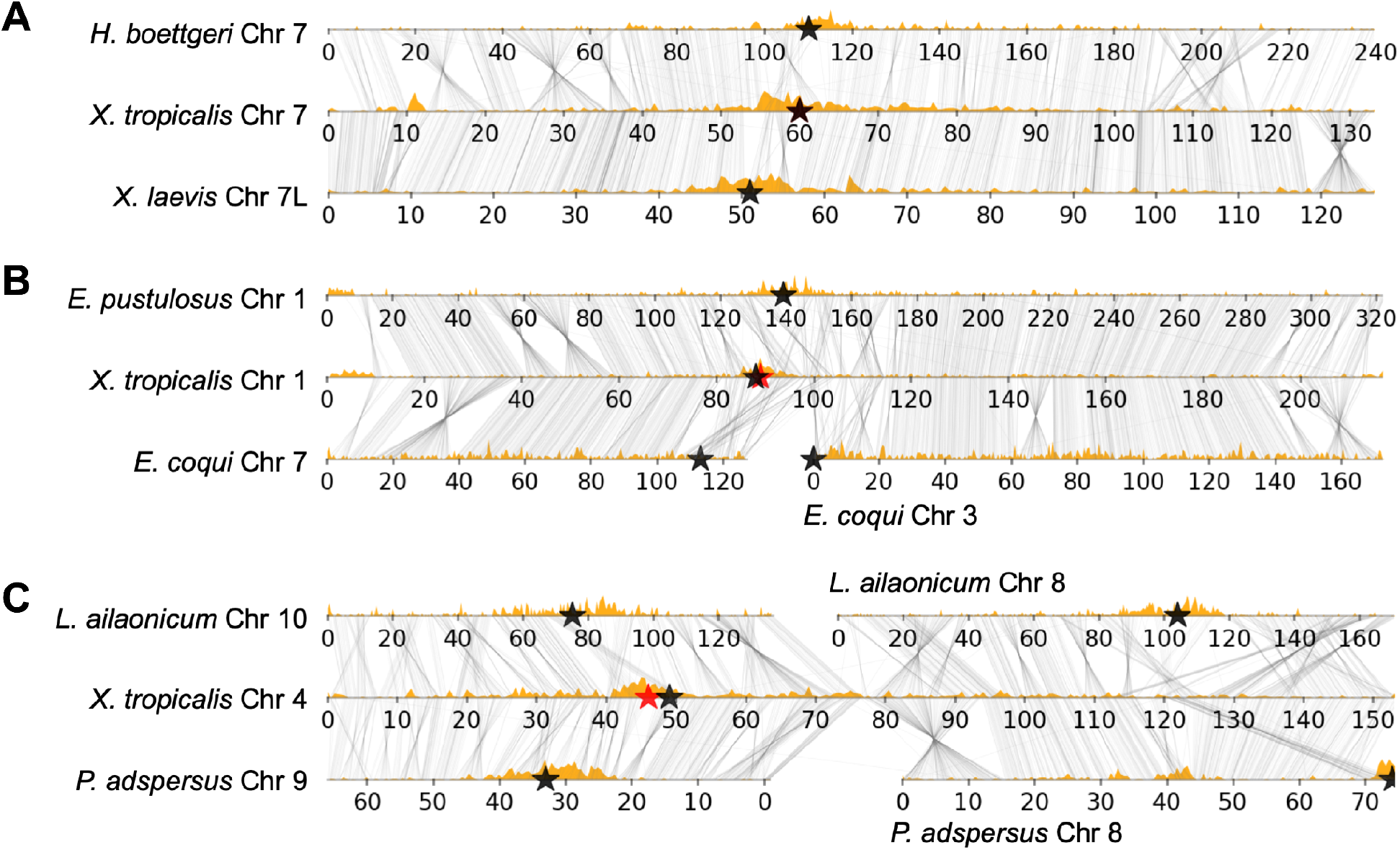
Subtelomeric repeats highlight regions of chromosome fusion. Examples of (**A**) conserved chromosome structure and pericentromere maintenance, (**B**) a Robertsonian translocation in the lineage leading to *E. coqui*, and (**C**) an end-to-end fusion that occurred in the lineage giving rise to *X. tropicalis* and subsequent pericentromere loss. The analyzed species were visualized with a custom script, alignment_plots.py (v1.0; https://github.com/abmudd/Assembly). For each plot, the HiC inference-based pericentromeric regions are depicted with black stars, the *X. tropicalis* centromeric satellite repeat from tandem repeat analysis with a red star, the density of L1 repeats per chromosome with light brown histograms, and the runs of collinearity containing at least one kb of aligned sequence between the species with connecting black lines.

Robertsonian or centric translocations involving breaks and joins near centromeres account for several of the rare rearrangements (**Figs. 1** and **3B**). For example, element G clearly experienced a centric fission in the *E. coqui* lineage. Conversely, I and M underwent centric fusion in the *E. pustulosus* lineage. *E. coqui* has experienced the most intense rearrangement, including Robertsonian fissions of A and G, a Robertsonian fusion of I/K, and a significant series of Robertsonian rearrangements involving B, E, F, and H that resulted in Bprox/H, Bdist/Fdist, and E/Fprox (**Table 1**, **Supplementary Table 8**). (Mechanistically, these “fissions” and “fusions” likely occur by translocations; for a discussion see ref.^62^.) Elements I and H form the two arms of a metacentric chromosome in pipids (**Fig. 3A**), and therefore the pipid ancestor, but are found as either independent acrocentric chromosomes (e.g., in *P. adspersus* and *L. ailaonicum*) or as arms of metacentrics formed by centric fusion with other elements (**Supplementary Table 8**).

We also observed end-to-end “fusions’’ of metacentric chromosomes, for example, the joining of D with K in *E. pustulosus*, and with element E in the common ancestor of pipids (*Hymenochirus* and *Xenopus*) (**Figs. 1** and **3C**). Since bicentric chromosomes are not stably propagated through mitosis, one of the two ancestral centromeres brought together by end-to-end fusion must be lost or inactivated, as shown in **Fig. 3C** for the ancient D-E fusion in pipids. We note that the D centromere persists in both end-to-end fusions involving D, suggesting that centromeres derived from different ancestral elements may be differentially susceptible to silencing.

Using the pericentromere and subtelomere repeat landscape as a proxy, we found several examples of end-to-end chromosome fusions in which residual subtelomeric signals are preserved near the presumptive junction (**Fig. 3**, **Supplementary Fig. 9**). These include the end-to-end fusion of *X. tropicalis*-like chromosomes 9 and 10 (elements L and M) to produce the Chr9_10 progenitor of *X. laevis* that is found in both the L and S subgenomes of this allotetraploid^20^. These *X. laevis* chromosomes display evidence of decaying subtelomeric signatures in the region surrounding the ancestral L-M fusion (**Fig. 1** and **Supplementary Fig. 9A,B**). Similarly, enrichment of subtelomerically associated repeats is observed in *H. boettgeri* chromosome 8_10 (**Supplementary Fig. 9C–E**) near the junction between the portions of the chromosome with M and J/K ancestry (the J/K fusion occurred near the base of pipids). In both cases, the centromere from element M (i.e., the centromere in *X. tropicalis* chromosome 9) is maintained after fusion. The inversion of the p-arm from Chr8S also has evidence of decaying sequence but the median is less than the median JC distance at the Chr9_10 fusion, suggesting that the fusion preceded the inversion.

#### Rate of karyotype change

The long-range and, in most cases, chromosome-scale collinearity (**Supplementary Fig. 8, Supplementary Table 9**) among the frog species we examined, despite a combined branch length of 1.05 billion years (**Supplementary Tables 10** and **11**), parallels the synteny observed in birds^63^ and reptiles^64^ but differs from the substantial chromosome variation found in mammals^15,31^. Maintenance of collinear blocks may reflect an intrinsically slow rate of rearrangement in frogs, perhaps a consequence of large regions devoid of recombination, or selection favoring retention of specific gene order and chromosome structure related to chromosomal functions. We inferred a total of 17 fission, fusion, translocation, and duplication events (excluding smaller intra-chromosome rearrangements) resulting in a karyotype change every 62 million years (**Fig. 1**). This rate is similar to the rate of one chromosome-number change every 70 to 90 million years as previously proposed for frogs and mammals^26,28^ but still slower than karyotype change rates for most mammals^65^ and many reptiles^66^. Of course, our rate calculation is based on only seven species, and the rate may vary depending on the species analyzed. Some frog taxa, such as *Eleutherodactylus* spp. (2*n* = 16–32) and *Pristimantis* spp.^39^ (2*n* = 22–38), have had a higher rate of karyotype change. On the other hand, some species, such as *Leptobrachium ailaonicum*, *L. leishanense*^14^, and *Rana temporaria*^102^, may have had no significant inter-chromosome changes over the past 205 million years (**Fig. 1**). Nonetheless, this analysis of chromosome variation across the frog lineage suggests a remarkably slow rate of karyotype evolution.

### Chromosome structure and conformation

The stasis of *Xenopus* chromosomes relative to other frogs (see below) allows us to examine the repetitive landscape of chromosomes that are not frequently rearranged by translocation, and may be approaching a structural equilibrium.

#### Centromeres, satellites, and pericentromeric repeats

Vertebrate centromeres are typically characterized by tandem families of centromeric satellites (e.g., the alpha satellites of humans) that bind to the centromeric histone H3 protein, Cenp-a, a centromere-specific variant of histone H3^53,67^. Cenp-a binding satellites have been described in *X. laevis*^68^, and here we find distantly related *X. tropicalis* satellite sequences that also co-precipitate with Cenp-a. Thus, chromatin immunoprecipitation and sequencing (ChIP-seq) shows that Cenp-a binding coincides with the predictions of centromere positions derived from chromatin conformation analysis and repetitive content (**Supplementary Fig. 5A–C**, **Supplementary Tables 12** and **13**). Importantly, this concordance supports the prediction of centromere position for other species that we infer below. The Cenp-a bound-sequences are arrays of 205-bp monomers that share a mean sequence identity greater than 95% at the nucleotide level, with a specific segment of the repeating unit showing greatest variability (**Supplementary Fig. 10**). The *X. tropicalis* centromere sequence is different from centromeric-associated repeats found in *X. laevis*^68,69^, suggesting the sequences evolve rapidly after speciation but are maintained within the species.

All metacentric pericentromeric regions of *X. tropicalis* chromosomes are enriched in retrotransposable repetitive elements (15 Mb regions shown in **Fig. 2**). In other vertebrate species and *Drosophila*, retrotransposable elements from the pericentromeric regions are involved in the recruitment of constitutive heterochromatin components^70,71^. Among the pericentromerically enriched repeats we identified specific families belonging to LTR retrotransposons (Ty3), non-LTR retrotransposons (CR1, Penelope, and L1), and DNA transposable elements (PIF-Harbinger and piggyBac families) (**Fig. 2**, **Supplementary Fig. 5**). CR1 (CR1-2_XT) is the most prevalent and it is among the youngest of all pericentromeric retrotransposons (mean Jukes-Cantor (JC) distance to consensus of 0.05). In contrast, L1 and Penelope types have a mean JC greater than 0.4 (**Supplementary Fig. 5**). The age of the repeats, indirectly measured by the JC distance, suggests that pericentromeric retrotransposons have experienced different bursts of activity and tendency to insert near the centromere. Expression of active retrotransposons and random insertion can compromise chromosome stability, and because silencing of these is crucial, genomes develop mechanisms to rapidly silence them. Such insertions may be positively selected, and therefore amplified, to establish pericentromeric heterochromatin, but may be counter selected when they insert in gene rich chromosome arms.

#### Recombination and extended subtelomeres

Although meiotic recombination is distributed across chromosomes, it is enriched near the chromosome ends (**Supplementary Fig. 11A**). While in humans, meiotic recombination is suppressed close to centromeres and elevated near telomeres, recombination is still regularly distributed on chromosome arms^72^. Other groups, including birds and fish, experience most recombination events 5 Mb away from the telomeres and only modest recombination is observed outside those regions^73–76^. Binding events for the protein PRDM9, present in mouse, rat and human, mark recombination hotspots in the chromosome arms in these species^77^. Given that amphibians lack the *prdm9* gene^78^, we analyzed the genomic features that colocalized in areas prone to recombination.

We studied the distribution of recombination along *X. tropicalis* chromosomes using a previously generated Nigerian-Ivory Coast F_2_ cross^18^ (**Supplementary Note 5, Supplementary Data 2**). Half of the observed recombination is concentrated in only 160 Mb (11.0% of the genome) and 90% of the observed recombination occurs in 540 Mb (37.3%). In contrast, the central regions of each chromosome are “cold”, with recombination rates below 0.5 cM/Mb (**Supplementary Fig. 11B, Supplementary Table 14**). Strikingly, we find that (sex-averaged) recombination is concentrated within just 30 Mb of the ends of each chromosome and occurs only rarely elsewhere (**Supplementary Fig. 11A**); the regions of the subtelomeres experiencing high recombination are nearly 6-fold larger than in non-amphibian genomes^73,74^. These rates of recombination were not previously determined, since the repeat-rich subtelomeres were absent from the assemblies, and markers that happened to lie in those regions showed insufficient linkage to be incorporated into the maps.

Due to the elevated recombination, and repeat structure discussed below, we defined the extended sub-telomeres as the terminal 30 Mb of all metacentric chromosomes, and terminal 30 Mb excluding the 15 Mb surrounding the pericentromeric regions of acrocentric chromosomes (Chr3, Chr8, and Chr10) (**Fig. 2**). The median recombination rate in the extended subtelomeres (1.73 cM/Mb) is ten-fold higher than the median rate observed in the rest of the chromosome arms (0.16 cM/Mb). The recombination rate in the 5-Mb region surrounding the centromeric tandem repeats is even lower (0.04 cM/Mb). Since constitutive heterochromatin in pericentromeric regions is known to repress recombination, this observation is expected (reviewed in refs.^79,80^). However, the centromeres of acrocentric chromosomes lie within 30 Mb of the telomere, which precludes the extended sub-telomeric associated repeats (**Fig. 2** and **Supplementary Fig. 12A**).

We examined the relationship between rates of recombination against repetitive elements and sequence motifs associated with recombination hotspots in other vertebrate species (**Supplementary Fig. 13A, Supplementary Table 14**). Similar to chicken and zebra finch, recombination is the highest in subtelomeres and positively correlates with GC content^73,76,81^, which is consistent with GC-biased gene conversion^82–84^ in recombinogenic regions (median GC = 42.5% in the 74 Mb in which half of the recombination occurs) vs. the non-recombinogenic centers of chromosomes (median 38.8%). As in zebra finch (**Supplementary Fig. 14**), recombination in *X. tropicalis* is strongly correlated with satellite repeats (Pearson correlation = +0.68, R^2^ = 0.457). The high density of satellite repeats (**Supplementary Table 15**) in highly recombinogenic subtelomeric regions suggests that unequal crossing over during meiotic recombination mediates tandem repeat expansions^85,86^. Notably in the extended subtelomeric regions, tandem repeats are enriched in specific tetrameric sequences (TGGG, AGGG, and ACAG) compared to non-tandem repeats (**Supplementary Fig. 13B**). In contrast, centromeric tandem repeats are completely devoid of these short sequences.

Some of the tandem arrays enriched in the terminal 30 Mb from all chromosomes derive from portions of transposable elements such as SINE/tRNA-V, LINE/CR1, DNA/Kolobok-2 (**Supplementary Fig. 12, Supplementary Table 16**). For example the minisatellite expansion that arose from the family of SINE/tRNA-V present in the pipid lineage^87^ amplified a 52-bp portion of the 3’UTR-tail from the SINE/tRNA-V element in *Xenopus tropicalis* and other frog species (**Supplementary Table 17**). Although intact SINE/tRNA-V elements are distributed throughout the genome, the minisatellite fragment is only expanded in subtelomeric SINE/tRNA-Vs, suggesting that recombination in subtelomeres has driven minisatellite expansion (**Supplementary Figs. 12** and **15**). Interestingly, although the satellite expansions are similar in *X. laevis* and *X. tropicalis*, they differ in other frogs, suggesting that different satellite expansions can occur repeatedly during the maintenance of the long subtelomeric regions (see below).

#### Chromatin conformation correlates with cytogenetic features

To further refine our understanding of chromosome structure in *X. tropicalis*, we studied chromatin conformation capture (“HiC”) data from nucleated blood cells. These experiments link short reads representing sequences in close three-dimensional proximity^88^. **Fig. 4** shows mapped HiC read pairs for chromosomes 1 and 2, with different minimum mapping quality thresholds above and below the diagonal (**Fig. 4** and **Supplementary Fig. 2, Supplementary Note 5**). We consistently observe a “wing” of intra-chromosome contacts transverse to the main diagonal, which (1) intersects the main diagonal near the cytogenetically defined, Cenp-a-binding centromere, and (2) indicates contacts between p and q arms (**Supplementary Figs. 2** and **16**). These observations imply that interphase chromosomes are “folded” at their centromeres, with contacts between distal arms. We also observe inter-chromosome contacts among centromeres of different chromosomes and between their telomeres (**Supplementary Fig. 10A**).

**Fig. 4.**
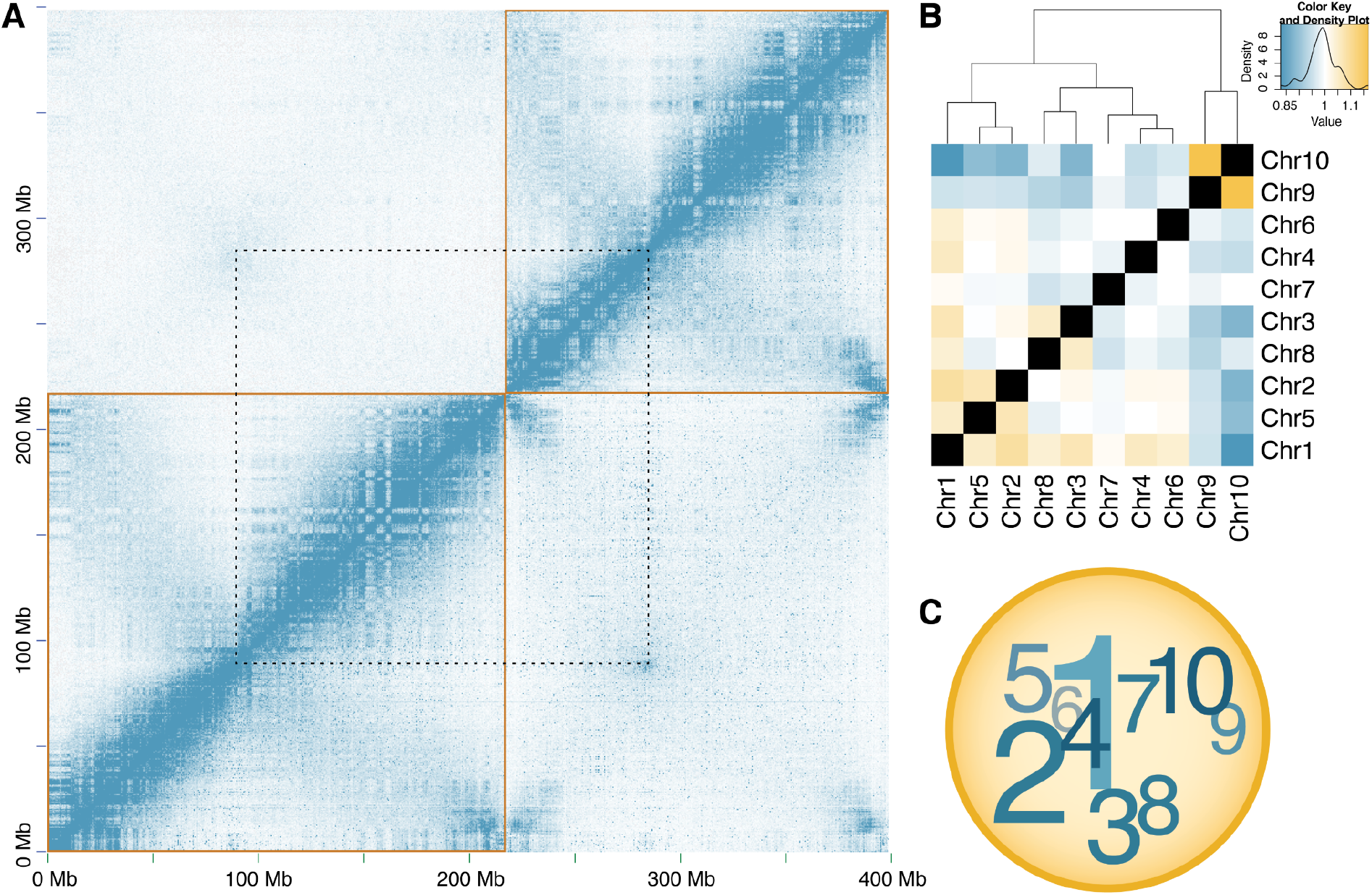
Organization of *X. tropicalis* chromosomes into Rabl configuration and distinct nuclear territories. (**A**) HiC contact matrices at 500 kb resolution for chromosomes 1 and 2 (lower-left and upper-right gold boxes, respectively) showing features of the three-dimensional chromatin architecture within *X. tropicalis* blood cell nuclei. Blue pixels represent chromatin contacts between X-Y pairs of genomic loci, and their intensity is proportional to their contact frequency. HiC read pairs are mapped stringently (MapQ ≥ 30) above the diagonal and permissively (MapQ ≥ 0) below the diagonal. The characteristic A/B compartment (“checkerboard”) and Rabl-like inter-arm (“angel wing”) contact patterns within each chromosome are evident. Above the diagonal, an increased frequency of inter-chromosomal chromatin contacts is observed between pericentromeres (connected by dotted lines) and between chromosome arms, suggesting a centromere-clustered organization of chromosomes in Rabl configuration. Below the diagonal, high-intensity pixels not present above the diagonal are present near the ends of chromosomes, suggesting a telomere-proximal spatial bias in the distributions of similar genomic repeats. See **Supplementary Fig. 1D** for a plot showing all chromosomes. (**B**) Chromosome territories within the nucleus. Yellow, white, and blue colors indicate the normalized relative enrichment, parity, and depletion of chromatin contacts between non-homologous chromosomes in the nucleus. For example, Chromosome 1 exhibits higher relative contact frequencies with all chromosomes except chromosomes Chr7, Chr9, and Chr10, which are generally depleted of contacts except among themselves. (MapQ ≥ 30; *χ*^2^ (81, *n* = 24,987,749) = 3,049,787; *p* < 2.2×10^−308^; Relative range: 0.82774–1.16834). (**C**) Schematic representation of chromosome territories within the nucleus. Chromosome number size is proportional to the number of enriched interactions. Darker and lighter colors indicate chromosomes nearer or distant to the reader, respectively.

Taken together, these intra- and inter-chromosome contacts are consistent with a Rabl configuration of chromosomes^89,90^ in *Xenopus* blood cells. This configuration is understood as a relict structure from the previous mitosis^91,92^, in which the chromosomes have become elongated and telomeres clustered on the nuclear membrane. Associations between centromeres and between telomeres, first observed in salamander embryos^89^, are also observed in other animals^93,94^, fungi^95^, and plants^96–98^. These findings suggest that remnants of this ‘Rabl configuration’^89^ may be a common feature of post-mitotic cells across a wide range of eukaryotes. Here, we quantified the degree to which chromosomes are compacted in the Rabl configuration using HiC data and find that, with the exception of blood cell nuclei (sum of squared distances (SSD) 1.465), chromosomes from early frog development (NF stages 8 to 23) appear more tightly constrained (mean SSD 1.384) in Rabl configuration than the more specialized (liver and brain) adult tissues and sperm (mean SSD 5.583; **Supplementary Fig. 16, Supplementary Table 18, Supplementary Note 5**). Although it is possible some amount of HiC signal may be due to residual incompleteness in the assembly and concomitant mismapping of reads to repeat sequences, these observations are robust to quality filtering, even when using single-copy sequences. Furthermore, such contacts are weakest in sperm^16^, a control that argues strongly against sequence mismapping artefacts (**Supplementary Fig. 10B, Supplementary Note 5**).

We also observed three-dimensional associations between pericentromeric regions of different chromosomes, based on enriched HiC contacts^90,99^ (**Fig. 4**). As with the Rabl signal, these “contacts” are accentuated when HiC reads are allowed to map permissively (**Methods**), which suggests that they may be influenced by common repetitive pericentromeric sequences shared among chromosomes. The signal persists in weaker form with more stringent read mapping, however, and either represents *bona fide* signal or residual incompleteness of the pericentromeric assembly. Notably, the correlation between centromere position and the observed intra-chromosome folding and inter-chromosome contacts at centromeres allows us to use HiC analysis and principal component analysis (PCA) of intra- and inter-chromosome contacts^97^ to infer the likely centromeric positions based purely on HiC data in frogs whose cytogenetics are less well-studied (see below).

#### Chromatin compartments

Chromatin contacts in human^88,100,101^, mouse^101^, chicken^102^ and other phylogenetically diverse species^103–105^ often show a characteristic checkerboard pattern that is superimposed on the predominant near-diagonal signal. This pattern implies an alternating ‘A/B’ compartment structure with enriched intra-compartment contacts within chromosomes (**Fig. 5A**), which has been linked with G-banding in humans^106^. *X. tropicalis* also exhibits an A/B compartment pattern, which emerges as alternating gene rich (‘A’) and gene poor (‘B’) regions (median 19.99 genes/Mb in A and 9.99 genes/Mb in B) (**Fig. 5B**). A/B compartments are also differentiated by repetitive content^101^, with A-compartment domains showing slight enrichment (1.21–1.44 fold) in DNA transposons of the DNA/Kolobok-T2, DNA/hAT-Charlie, and Mariner-Tc1 families. B-compartment domains had significantly higher enrichment for DNA transposons (DNA/hAT-Ac, Mar-Tigger) and retrotransposons (Ty3/metaviridae and CR1), among other repeats (1.12–2.11 fold) (**Fig. 5C**). The association between repeats overrepresented in A and B compartments is also captured in one of the principal components obtained from the repeat densities of all chromosomes (**Supplementary Note 5**); we detect a modest negative correlation (R = −0.44) between the HiC eigenvectors that classified A/B compartments and the third principal component eigenvectors obtained from the repeat density matrix (**Supplementary Fig. 5B**). The association between chromatin condensation and repeat type could be due to a preference for certain transposable elements to insert in specific chromatin contexts, or chromatin condensation to be controlled in part by transposable element content, or a combination of these factors. However, we were unable to find any correlation of AB compartments with the G-banding of condensed chromosomes in *X. tropicalis*^107,108^.

**Fig. 5.**
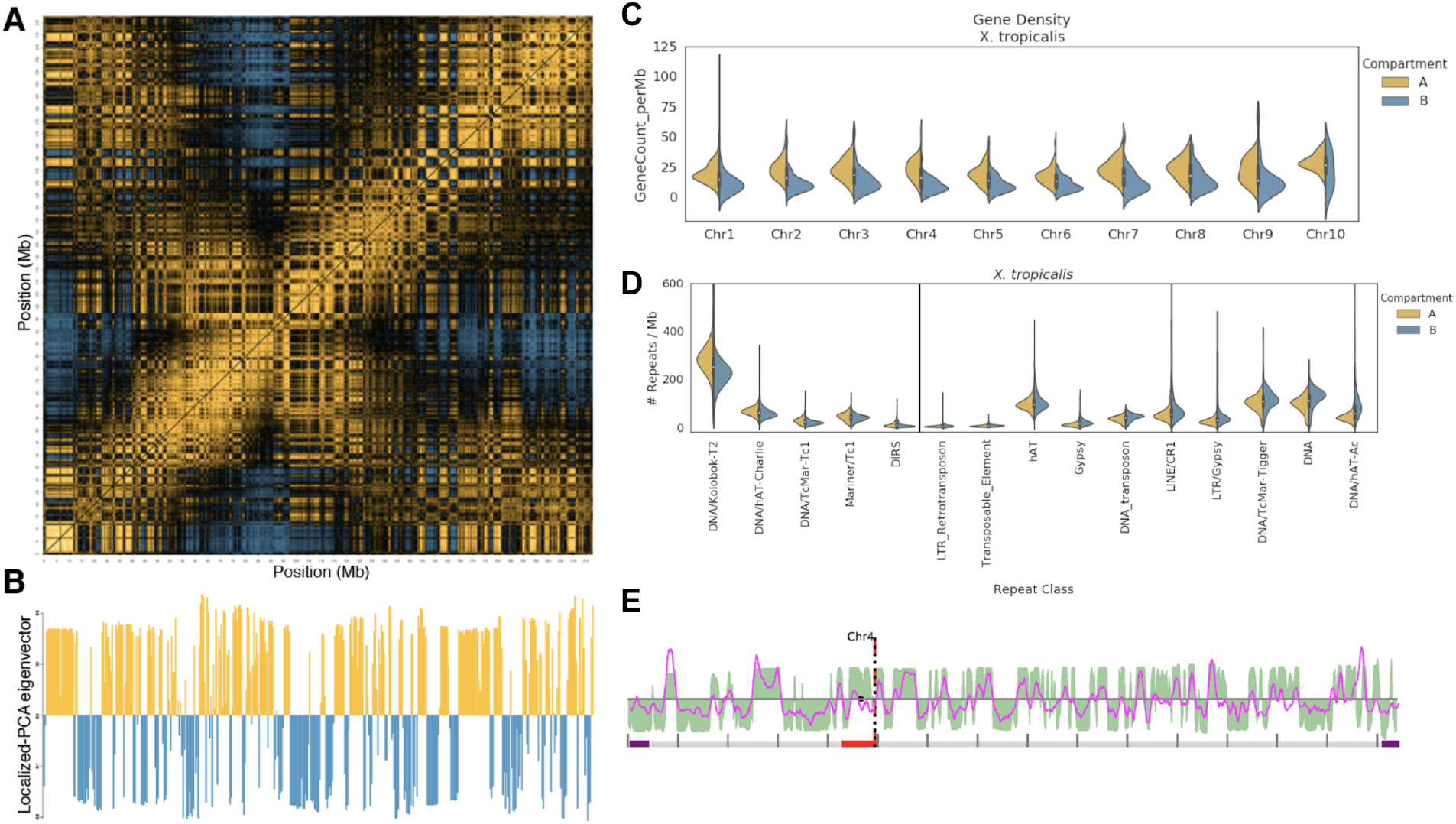
A/B compartment structure and gene/repeat densities. (**A**) Correlation matrix of intra-chromosomal HiC contact densities between all pairs of non-overlapping 250 kb loci on chromosome 1. Yellow and blue pixels indicate correlation and anti-correlation, respectively, and reveal which genomic loci occupy the same or different chromatin compartment. Black pixels indicate weak/no correlation. (**B**) The first principal component eigenvectors revealing compartment structure along chromosome 1, obtained by singular value decomposition of the correlation matrix in **A**. Yellow (positive) and blue (negative) eigenvectors indicate regions of chromosome 1 partitioned into the A and B compartments, respectively. (**C**) Gene density distributions in A vs. B compartments per chromosome. (**D**) Repeat classes significantly enriched in A vs. B compartments. (**E**) The eigenvectors obtained from the HiC correlation matrix (PC1, green) and the eigenvectors from the repeat density matrix (PC3, pink) coincide at the transitions of predicted A/B compartments (**Supplementary Fig. 16B**).

#### Higher-order interactions

Chromatin conformation contacts also provide clues to the organization of chromosomes within the nucleus. We observe non-random (*χ*^2^ (81, *n* = 24,987,749) = 3,049,787; *p* < 2.2×10^−308^) associations between chromosomes in blood cell nuclei (**Fig. 4B**, **Supplementary Tables 19** and **20**): (a) chromosome 1 is enriched for contacts with chromosomes 2–8 (mean 1.05× enrichment), and depleted of contacts with 9 and 10 (mean 0.89×); (b) among themselves, chromosomes 2–8 show differential contact enrichment or depletion; and (c) chromosomes 9 and 10 are enriched (1.17×) for contacts with one another but are depleted with respect to all other chromosomes. These observations suggest the presence of distinct chromosome territories^89,109–111^, where chromosomes 2–8 localized more proximal to—and arrayed around—chromosome 1, with chromosomes 9 and 10 relatively sequestered from chromosome 1 (**Fig. 4C**). The contact enrichment between chromosomes 9 and 10 is particularly notable because these short chromosomes (91.2 and 52.4 Mb, respectively) have become fused in the *X. laevis* lineage^112^, which might have been enabled by their persistent nuclear proximity^113^.

Between chromosomes, p-p and q-q arm interactions exhibit a small but significant enrichment (1.059× enrichment; *χ*^2^ (1, *n* = 24,786,496) = 17,037; *p* < 2.2×10^−308^) over p-q arm contacts. This is a general feature of metacentric and sub-metacentric chromosomes also observed in other frog genomes (see below), except *E. coqui* (0.928× enrichment; *χ*^2^ (1, *n* = 6,850,547) = 3,914; *p* < 2.2×10^−308^), the chromosomes of which are predominantly acrocentric. Finally, the p-arms of chromosomes 3, 4, 8, and 9 are enriched for contacts with chromosome 10, with the sub-metacentric chromosomes 3 and 8 showing the strongest enrichment (and a slight preference between p-arms). The q-arms of chromosomes 3 and 8, however, exhibit a slight enrichment for contacts with chromosomes 1, 2, 4, and 5. Taken together, these observations suggest colocalization of the p and q arms of chromosomes 3 and 8 in blood cell nuclei.

## Conclusions

Anuran amphibians play a central role in biology, not simply as a globally distributed animal taxon, but also as key subjects for research in areas that range from ecology and evolution to cell and developmental biology. The genomic resources generated here will thus provide important tools for further studies. Given the crucial role of *X. tropicalis* for genomic analysis of development and regeneration^114,115^, the improvements to our understanding of its genome reported here will provide a more finely grained view of biomedically important genetic and epigenetic mechanisms. This new genome is also important from the standpoint of evolutionary genomics, as comparisons between the genomes of *X. tropicalis* and *X. laevis* shed light on mechanisms of genome duplication^115^. The new genome described here for *H. boettgeri,* another pipid frog, is also significant in this regard, as it enables an interesting comparison of *Xenopus* genomes to that of a closely related outgroup. Moreover, the genomes of *E. coqui* and *E. pustulosus* provide a foundation for future studies of the evolution of ontogenies and of their underlying developmental mechanisms, as *E. coqui* is a direct-developing frog with no tadpole stage^116^ and *E. pustulosus,* a foam-nesting frog, is a model for studying mating calls and female mate choice^116^. In addition to their interesting life histories, both frogs display interesting patterns of gastrulation^117,118^. Finally, recent work has demonstrated the efficacy of genetic or genomic analysis for understanding the impact of chytrid fungus on various amphibian species^119^. A deeper and broader understanding of amphibian genomes will be useful in the context of the global decline of amphibian populations^120,121^.

## Online Methods

### Genomic extraction and sequencing

High molecular weight DNA was extracted from blood of an F_17_ *Xenopus tropicalis* Nigerian strain female (ref.^18^; **Supplementary Note 1**). Paired-end (PE) Illumina shotgun libraries were constructed by the QB3 Functional Genomics Laboratory (FGL) and sequenced on an Illumina HiSeq 2500 as 2×250 bp reads at the Vincent J. Coates Genomics Sequencing Lab (VCGSL) at the University of California, Berkeley. Single-molecule real-time (SMRT) continuous long-read (CLR) sequencing was performed at the HudsonAlpha Institute for Biotechnology on PacBio RSII machines with P6-C4 chemistry (**Supplementary Note 1, Supplementary Data 1**). 10x Genomics Chromium linked-read sequencing was carried out at HudsonAlpha on HiSeq XTen (**Supplementary Note 1**).

### *Xenopus tropicalis* genome assembly and annotation

Chromium linked reads (10x Genomics) were assembled with Supernova^122^ (v1.1.5). This assembly was used to seed the assembly of continuous long reads (PacBio) using DBG2OLC^123^ (commit 1f7e752). An independent PacBio-only assembly was constructed with Canu^124^ (v1.6-132-gf9284f8). These two assemblies were combined, or metassembled, using MUMmer^125^ (v3.23) and quickmerge^126^ commit e4ea490 (**Supplementary Fig. 1A, Supplementary Note 2**). Residual haplotypic redundancy was identified and removed (**Supplementary Fig. 1B, Supplementary Note 2**). The non-redundant metassembly was scaffolded with Sanger paired-ends and BAC-ends^33^ using SSPACE^127^ v3 and HiC using 3D-DNA^94,128^ commit 2796c3b, then manually curated in JuiceBox^129,130^ v1.9.0 (**Supplementary Note 2**). The assembly was polished with Arrow^131^, Pilon^132^ v1.23, and then FreeBayes^133^ (v1.1.0-54-g49413aa) with ILEC (map4cns; https://bitbucket.org/rokhsar-lab/map4cns). The genome was annotated with the DOE-JGI Integrated Gene Call pipeline^134^ (IGC) using transcript assemblies (TAs) generated with Trinity^135,136^ v2.5.1 from multiple developmental stages and tissues (**Supplementary Data 1, Supplementary Note 2**). RepeatModeler^137^ v1.0.11 was run on all frog species. The frog and ancestral repeat libraries from RepBase^138^ v23.12 were combined with the repeats consensuses identified by Repeat Modeler. The merged repeat library was used to annotate repeats of all frogs with RepeatMasker^139^ v4.0.7 (**Supplementary Notes 2** and **3**).

### *Hymenochirus boettgeri* metaphase chromosome spread

Stage 26 tadpoles (*n* = 10) were incubated at room temperature in 0.01% colchicine and 1× MMR for 4–6 hr. After removing the yolky ventral portion of the tadpoles, the remaining dorsal portions were pooled together in deionized water and allowed to stand for 20 min. The dorsal portions were transferred to 0.2 mL of 60% acetic acid in deionized water and allowed to stand for 5 min. The tissue was then pipetted onto a positively charged microscope slide, and excess acetic acid was blotted away. To flatten the tissue and promote chromosome spreading, the slide was covered with a coverslip and a lead brick was placed on top of it for 5 min. The slide and coverslip were then placed on dry ice for 5 min. The coverslip was removed from the frozen slide, and the slide was stained with 0.1 mg/mL Hoechst Stain solution for 5 min. A fresh coverslip was then mounted on the slide using VectaShield, and the edges were sealed with nail polish. Chromosomes in metaphase spreads (**Supplementary Fig. 7A**) were imaged on an Olympus BX51 Fluorescence Microscope run with Metamorph software using a 60× oil objective. Chromosome number was counted in 75 separate metaphase spreads.

### Genome and transcriptome sequencing of five pipanurans

Illumina PE 10x Genomics Chromium linked-read whole-genome libraries for *E. pustulosus* (from liver), *E. coqui* (from blood) and *H. boettgeri* (from liver) were sequenced on an HiSeq X at the HudsonAlpha Institute for Biotechnology. PacBio SMRT Sequel I CLR data were generated at UC Davis DNA Technologies and Expression Analysis Core for each of *E. pustulosus* and *H. boettgeri* from liver samples. In addition, two TruSeq Illumina PE libraries (from kidney) and two Nextera mate-pair libs (from liver) for *E. coqui* were prepared. HiC chromatin conformation capture libraries were prepared for *H. boettgeri*, *E. pustulosus*, and *E. coqui* using the Dovetail^TM^ HiC Kit for Illumina following the “Animal Tissue Samples” protocol. HiC libraries were sequenced on the Illumina HiSeq 4000 by the VCGSL.

Illumina TruSeq Stranded mRNA Library Prep Kit libraries were prepared from *E. pustulosus* stages 45 and 56 whole tadpoles (gut excluded) and various adult tissues dissected from frogs maintained at the University of the Pacific. Brain (*n* = 3), dorsal skin (*n* = 2), eggs (*n* = 2), eye (*n* = 2), heart (*n* = 2), intestine (*n* = 2), larynx (*n* = 3), liver (*n* = 2), lung (*n* = 2), and ventral skin (*n* = 2) samples were washed twice with PBS, homogenized in TRIzol Reagent, and centrifuged, followed by flash freezing of the supernatant. RNA was isolated following the *TRIzol Reagent User Guide* (Pub. No. MAN0001271 Rev. A.0) protocol. In addition, *H. boettgeri* eggs were homogenized in TRIzol Reagent and processed according to manufacturer’s instructions. RNA was then isolated using the QIAGEN RNeasy Mini Kit (cat 74104). An Illumina mRNA library was prepared using the Takara PrepX RNA-Seq for Illumina Library Kit by the Functional Genomics Laboratory at the University of California Berkeley. All libraries were sequenced at the VCGSL on an HiSeq 4000 as 151 bp PE reads. See **Supplementary Note 3** for additional details about DNA/RNA extractions and library preparations, and **Supplementary Data 1** for a complete list of DNA/RNA sequencing data generated for *E. coqui*, *E. pustulosus*, and *H. boettgeri*.

### Assembly and annotation of five pipanuran genomes

Contigs were assembled with Supernova^122^ v2.0.1 (*E. pustulosus* and *H. boettgeri*) or Meraculous^140,141^ v2.2.4 (*E. coqui*). For *E. coqui*, residual haplotypic redundancy was removed using custom scripts (https://github.com/abmudd/Assembly) prior to scaffolding with SSPACE^127^ (v3.0). *E. pustulosus* and *H. boettgeri* contigs were ordered and oriented using MUMmer^125^ (v3.23) alignments to PBEC-polished (map4cns commit dd89f52; https://bitbucket.org/rokhsar-lab/map4cns) DBG2OLC^123^ (commit 1f7e752) hybrid contigs (**Supplementary Note 3**). All three assemblies were scaffolded further with linked-reads and Scaff10X (v2.1; https://sourceforge.net/projects/phusion2/files/scaff10x).

*E. pustulosus* and *H. boettgeri* chromosome-scale scaffolds were constructed with Dovetail Genomics HiC via the HiRise scaffolder^142^, followed by manual curation in JuiceBox^128–130^ v1.9.0. Due to the fragmented nature of the *E. coqui* assembly, initial chromosome-scale scaffolds were first constructed by synteny with *E. pustulosus*, then refined in JuiceBox^128–130^ v1.9.0. Gaps in the *E. pustulosus* and *H. boettgeri* assemblies bridged by PacBio reads were resized using custom scripts (pbGapLen; https://bitbucket.org/bredeson/artisanal) and filled with PBJelly^143^ (PBSuite v15.8.24). These two assemblies were polished with FreeBayes and ILEC (map4cns commit dd89f52; https://bitbucket.org/rokhsar-lab/map4cns). A final round of gap-filling was then performed on the three assemblies using Platanus^144^ (v1.2.1).

Previously published *L. ailaonicum*^23^ (GCA_018994145.1) *and P. adspersus*^21^ (GCA_004786255.1) assemblies were manually corrected in JuiceBox^128–130^ (v1.11.08) using their respective HiC and Chicago data (**Supplementary Data 1**). Gaps in the corrected *P. adspersus* scaffolds were resized with PacBio reads (as described above) and filled using Platanus^144^ (v1.2.1) with published TruSeq Illumina data obtained from NCBI (PRJNA439445). As described elsewhere^145^, all assemblies were screened for contaminants prior to scaffolding, and only final scaffolds and contigs longer than 1 kb were retained for downstream analyses. More details on assembly procedures can be found in (**Supplementary Note 3**).

Genomic repeats in all five species were annotated with RepeatMasker^137,139^ (v4.0.7 and v4.0.9) using the repeat library generated above. Protein-coding genes were annotated for *E. coqui*, *E. pustulosus*, *H. boettgeri*, and *P. adspersus* using the IGC^134^ pipeline with homology and transcript evidence. For each respective species, newly generated RNA-seq data were combined with public *H. boettgeri*^20^ (BioProject PRJNA306175) and *P. adspersus*^21^ (BioProject PRJNA439445) data, and unpublished *E. coqui* data (stages 7, 10, and 13 hindlimb [Harvard University]; stage 9–10 tail fin skin [French National Center for Scientific Research]). Transcript assemblies used as input to IGC were assembled with Trinity^135,136^ (v2.5.1) and filtered using the heuristics described in **Supplementary Note 3.**

### Synteny and ancestral chromosome inference

One-to-one gene ortholog set between frog proteomes was obtained from the output from OrthoVenn2^52^ using an E-value of 1×10^−5^ and an inflation value of 1.5 (**Supplementary Note 4**). The assemblies of all frog species and axolotl were pairwise aligned against the *X. tropicalis* genome using cactus^146^ (commit e4d0859) (**Supplementary Note 4**). Pairwise collinearity runs were merged into runs of collinearity with ROAST/MULTIZ^147^ (v012109) using the phylogenetic topology from TimeTree^148^ and sorted with last^149^ (v979) (**Supplementary Note 4**).

### Phylogeny and estimation of sequence divergence

Fourfold degenerate bases of one-to-one orthologs were obtained and reformatted from the MAFFT alignment as described in Mudd et al.^145^ (**Supplementary Note 4**). The maximum-likelihood phylogeny was obtained with RAxML^150^ (v8.2.11) using the GTR+Gamma model of substitution with outgroup *Ambystoma mexicanum*. Divergence times were calculated with MEGA7^151^ (v7.0.26) with the GTR+Gamma model of substitution using Reltime method^152^.

### Chromosome evolution

Customized scripts^145^ were used to extract pairwise alignments from the ROAST-merged MAF file and converted into runs of collinearity. The runs of collinearity were visualized with Circos^153^ (v0.69-6) (**Supplementary Note 4**).

### Centromeres, satellites, and pericentromeric repeats

Tandem repeats were called using Tandem Repeat Finder^57^ (trf genome.fa 2 5 7 80 10 50 2000 -l 6 -d -ngs). To identify tandem repeats enriched in pericentromeric and subtelomeric regions, we extracted the monomer sequences of all tandem repeats overlapping the region of interest. A database of non-redundant monomers was created by making a dimer database. Dimers were clustered with BlastClust^154^ v2.2.26 (-S 75 -p F -L 0.45 -b F -W 10). A non-redundant monomer database was created using the most common monomer size from each cluster. The non-redundant sequences were mapped to the genome with BLASTN^155^ (-outfmt 6 -evalue 1e3). The enriched monomeric sequences in centromeres and subtelomeres were identified by selecting the highest normalized rations of tandem sequence footprints in the region of interest over the remaining portions of the genome. For more detail, see **Supplementary Note 5**.

### Genetic variation

Reads were aligned with BWA-MEM^156^ (v0.7.17-r1188) and alignments were processed using SAMtools^157^ (v1.9-93-g0ca96a4), keeping only properly paired reads (samtools view -f3 -F3852) for variant calling. Variants were called with FreeBayes^133^ (v1.1.0-54-g49413aa; --standard-filters --genotype-qualities --strict-vcf --report-monomorphic). Only bi-allelic SNPs with depth within mode ± 1.78SDs were retained. An allele-balance filter [0.3–0.7] for heterozygous genotypes was also applied. Segmental heterozygosity/homozygosity were estimated using windows of 500 kb with 50-kb step using BEDtools^158^ (v2.28.0) for pooled samples or snvrate^159^ (v2.0; https://bitbucket.org/rokhsar-lab/wgs-analysis). For more detail, see **Supplementary Note 2**.

### GC-content, gene, and repeat landscape

GC-content percentages were calculated in 1-Mb bins sliding every 50 kb. Gene densities were obtained using a window size of 250 kb sliding every 12.5 kb. The repeat density matrix for *X. tropicalis* was obtained by counting base pairs per 1 Mb (sliding every 200 kb) covered by repeat families and classes of repeats. The principal component analysis (PCA) was performed on the density matrix composed of 7,253 1-Mb bins and 3,070 repeats (**Supplementary Note 5**). The first (PC1) and second (PC2) components were smoothed using a cubic spline method.

### Chromatin immunoprecipitation

*Xenopus tropicalis* XTN-6 cells (Gorbsky and Horb, unpublished) were grown in 70% calcium-free L-15 (US Biologicals cat# L2101-02-50L), pH 7.2/10% Fetal Bovine Serum/Penicillin-Streptomycin (Invitrogen cat# 15140-163) at RT. Native MNase ChIP-seq protocol performed as described previously in Smith et al.^69^. Approximately 40 million cells were trypsinized and collected; nuclei were isolated by dounce extraction and collected with a sucrose cushion. Chromatin was digested to mononucleosomes by MNase. Nuclei were lysed and soluble nucleosomes were extracted overnight at 4 °C. Extracted mononucleosomes were precleared with Protein A dynabeads (Invitrogen cat# 100-02D) for at least 4 h at 4 °C. A sample was taken for input after preclearing. Protein A dynabeads were bound to 10-μg antibody (either Rb-anti-Xl Cenp-a, cross-reactive with *X. tropicalis*, Rb-anti-H4 Abcam cat# 7311 or Rb-anti-H3 Abcam cat# 1791) and incubated overnight with precleared soluble mononucleosomes at 4 °C. Dynabeads bound to Rabbit IgG antibody (Jackson ImmunoResearch cat#011-000-003) were collected with a magnet and washed three times with TBST (0.1% Triton X-100) before elution with 0.1% SDS in TE and proteinase K incubation at 65 °C with shaking for at least 4 h. Isolated and input mononucleosomes were size-selected using Ampure beads (Beckman cat# A63880) and prepared for sequencing using the NEBNext ultra ii DNA library prep kit for Illumina (NEB cat# E7654). Three replicates were sequenced on an Illumina HiSeq 4000 lane 2×150 bp by the Stanford Functional Genomics Facility. PE reads were trimmed with Trimmomatic^160^ v0.39 filtering for universal Illumina primers and for Nextera-PE indices. Processed PE reads were mapped with minimap2 (ref.^161^) v2.17-r941 against the unmasked genome reference. Samtools^157^ v1.9 was used for sorting and indexing the alignment. Read counts (MapQ0) per 10-kb bin (non-overlapping) for all samples were calculated with multiBamSummary from deeptools^162^ v3.3.0. Read counts were normalized by the total number of counts in the chromosomes per sample (**Supplementary Note 5**).

### Recombination and extended subtelomeres

The reads from the F_2_ mapping population^18^ were aligned to the v10 genome using BWA-MEM^156^ (v0.7.17-r1188). Variants were called using FreeBayes^133^ (v1.1.0-54-g49413aa; -- standard-filters --genotype-qualities --strict-vcf). SNPs were filtered, and valid F_2_ mapping sites were selected when the genotypes of the Nigerian F0 and the ICB F_0_ were fixed and different and there was a depth of at least 10 for each F_0_ SNP. Maps were calculated using JoinMap^163^ v4.1 (**Supplementary Note 5, Supplementary Data 2**). The variation on the linkage map was smoothed using the cubic spline function calculated every 500 kb. The Pearson correlation coefficient was calculated between recombination rates and genomic features that include GC content, repeat densities, and densities of reported CTCF and recombination hotspots^164,165^.

### Chromatin conformations and higher-order interactions

HiC read pairs were mapped with Juicer^128^ (v1.5.6) and observed counts were extracted at 1 Mb resolution with JuicerTools. Centromeres were estimated manually in JuiceBox^129^ and refined with Centurion^99^ v0.1.0-3-g985439c using ICE-balanced MapQ0 matrices (https://bitbucket.org/rokhsar-lab/xentr10/src/master/hic). Rabl chromatin structure was visualized with PCA from Knight-Ruiz^1^^66^-balanced MapQ30 matrices, and significance estimated by permutation testing using custom R scripts. Rabl constraint between p- and q-arms was measured as the sum of square distances (SSD) in PC1-PC2 dimensions, calculated between non-overlapping bins traveling sequentially away from the centromere. Inter-/intra-chromosomal contact enrichment analyses were quantified from MapQ30 matrices using *χ*^2^ tests in R^167^ v3.5.0 (**Supplementary Note 5**; https://bitbucket.org/rokhsar-lab/xentr10/src/master/hic).

### A/B compartments

A/B compartments were called with custom R^167^ (v3.5.0) scripts from Knight-Ruiz-balanced (observed / expected normalized) MapQ30 HiC contact correlation matrices generated with Juicer^128^ (**Supplementary Note 5**). Pearson’s correlation between eigenvectors of PC1 from the HiC correlation matrix and gene density were used to designate A and B compartments per chromosome.

## Data Availability

The assemblies, annotations, and raw data are deposited in NCBI for v10 *X. tropicalis* (BioProjects PRJNA577946 and PRJNA726269), *E. coqui* (BioProject PRJNA578591), *E. pustulosus* (BioProject PRJNA578590), *H. boettgeri* (BioProject PRJNA578589), *L. ailaonicum* (BioProject PRJNA578588), and *P. adspersus* (BioProject PRJNA578592).

## Code Availability

All custom scripts used in this work can be found at https://bitbucket.org/rokhsar-lab/xentr10 and https://github.com/abmudd/Assembly.

## Acknowledgements

We thank Karen Lundy and the Functional Genomics Laboratory at the University of California Berkeley for running quality control on extracted DNA and RNA and for preparing Illumina short-insert libraries; Oanh Nguyen and the DNA Technologies and Expression Analysis Cores at the University of California Davis Genome Center for preparing and sequencing PacBio libraries; Dovetail Genomics for providing the HiC library preparation kit, running quality control on HiC libraries, and preparing and sequencing HiC libraries; Shana McDevitt and the Vincent J. Coates Genomics Sequencing Laboratory at the University of California Berkeley for sequencing HiC and Illumina short-insert libraries; Shengqiang Shu for advice on the use of the IGC annotation pipeline. We thank Rick Elinson for providing *E. coqui* frogs and tissues. We thank Gary Gorbsky from the Oklahoma Medical Research Foundation and Marko Horb and the National *Xenopus* Resource at the MBL for providing the XTN-6 cell lines. Finally, we thank Chunhui Hou and colleagues for permission to access their HiC data prior to publication.

## Author Information

### Contributions

J.V.B., A.B.M., S.M-R., T.M., R.M.H., and D.S.R. wrote the manuscript with feedback from M.L., H.P.S., J.H., J.B.L., J.B.W., M.J.R., O.K.S., D.R.B., M.G-P., J.H., N.B., T.K., L.M.S., R.H., J.S., M.K.K., A.F.S., and D.H. Genomes were assembled by J.V.B., S.S.B. (*Xtr*); A.B.M., and K.C.B. (other frogs). S.M-R., A.B.M., and G.K. assembled transcripts and annotated genomes. S.M-R. and J.V.B. assessed gene completeness; S.M-R. analyzed repeat and recombination landscapes. S.M-R. and J.P. identified centromeric repeats. O.K.S., G.A-F. and A.F.S. conducted ChIP-seq experiments and S.M-R. performed analysis. J.V.B. analyzed HiC features. T.M. constructed the linkage map. T.M. and J.V.B. analyzed heterozygosity. A.B.M. performed genome-wide comparisons. K.E.M. and R.H. examined *Hbo* metaphase spreads. M.K.K. and M.L. inbred *Xtr* frogs. R.M.H. (*Xtr*); M.G-P. (*Epu*); K.E.M. and R.H. (*Hbo*); M.L. and J.H. (*Eco*) collected frogs. R.M.H. (*Xtr*); M.G-P., H.S-P. (*Epu*); and D.R.B. (*Eco*) collected tissue samples. A.B.M., D.R.B. (*Eco*); J.B.L., and I.P. (*Xtr*) extracted DNA. A.B.M., S.M-R. (*Epu*); K.E.M., R.H. (*Hbo*); and L.M.S. (*Eco*) extracted RNA and libraries were prepared by A.B.M. (*Epu*). M.L., J.H. (*Eco*); K.E.M., and R.H. (*Hbo*) provided RNA-seq data. T.K., M.J.R., J.B.W. (*Epu*); and J.B.L. (*Xtr*) coordinated sequencing. C.P., J.G., and J.S. prepared and sequenced 10x Genomics, PacBio, and Illumina mate-pair libraries. D.H. prepared HiC libraries. R.D.D. and J.H.M. provided early access to the *Pad* assembly. N.B. (*Eco*) provided bioinformatic support. L.M.S. led the *Eco* efforts. R.M.H. and D.S.R. led the project.

## Supplementary Information

(Provided in a separate document)

Supplementary Fig. 1 Genome assembly and recovery of missing genes.

Supplementary Fig. 2 *Xenopus tropicalis* genome-wide HiC contact map.

Supplementary Fig. 3 GC landscape and tandem repeats.

Supplementary Fig. 4 Comparison of gene content in assemblies of model vertebrates.

Supplementary Fig. 5 PCA eigenvectors projected on genomic coordinates.

Supplementary Fig. 6 *Xenopus tropicalis* Nigerian strain residual heterozygosity.

Supplementary Fig. 7 Assembly and annotation of other frog species.

Supplementary Fig. 8 Pairwise gene colinearity of frog genomes.

Supplementary Fig. 9 Chromosome fusions in *Xenopus laevis* and *Hymenochirus boettgeri*.

Supplementary Fig. 10 Estimating the positions of *Xenopus tropicalis* centromeres.

Supplementary Fig. 11 *Xenopus tropicalis* recombination landscape.

Supplementary Fig. 12 Distribution of Satellite repeats in *Xenopus tropicalis*.

Supplementary Fig. 13 Correlates of recombination rate.

Supplementary Fig. 14 Zebra finch subtelomeric tandem repeats.

Supplementary Fig. 15 Microsatellite origin SINE/tRNA evolved into a microsatellite sequence.

Supplementary Fig. 16 *Xenopus tropicalis* 3D chromatin structure and nuclear organization.

Supplementary Table 1 Sequence Completeness.

Supplementary Table 2 Transcript coverage of *X. tropicalis* assemblies v9 and v10.

Supplementary Table 3 *Xenopus tropicalis* protein coding loci annotation summary statistics.

Supplementary Table 4 *Xenopus tropicalis* repeat abundances.

Supplementary Table 5 Summary of other frog genome assemblies.

Supplementary Table 6 Summary of annotations for other frog genomes.

Supplementary Table 7 BUSCO genome scores of other frog genome assemblies.

Supplementary Table 8 Ancestral chromosome fusions.

Supplementary Table 9 N50 lengths for collinear runs of orthologous genes between frogs.

Supplementary Table 10 Four-fold degeneracy nucleotide divergence.

Supplementary Table 11 Estimation of divergence times.

Supplementary Table 12 Centromeric Associated Tandem Repeat monomer lengths and counts.

Supplementary Table 13 Mapping statistics for ChIP-seq samples.

Supplementary Table 14 Correlates of recombination rate.

Supplementary Table 15 Subtelomeric enrichment for tandem repeats.

Supplementary Table 16 Correspondence of monomer sequence with annotated repeat elements.

Supplementary Table 17 Copy counts of 52-mer minisatellite.

Supplementary Table 18 Quantification of Rabl structure strength and significance.

Supplementary Table 19 Contact enrichment between chromosomes and chromosome arms.

Supplementary Table 20 Relative enrichment of HiC contacts between chromosomes.

Supplementary Note 1 High-throughput sequencing, *Xenopus tropicalis*.

Supplementary Note 2 Xenopus *tropicalis* genome assembly and annotation.

Supplementary Note 3 Additional chromosome-scale frog assemblies.

Supplementary Note 4 Comparative analysis.

Supplementary Note 5 Genome analysis.

## Supplementary Data Files

**Supplementary Data 1: Table of sequencing data.**

An MS Excel file summarizing the sequencing data used to construct the six frog genome assemblies new or updated in this study, as well as the RNA-seq data used for annotating their protein-coding genes. The *X. tropicalis* ChIP-seq data are also included.

Supplementary Data 2: Genetic markers.

An MS Excel file containing the marker number, locus identifier, chromosome name, centiMorgan position, and chromosome coordinate for each genetic marker in the F_2_ *X. tropicalis* genetic linkage map.

