## Supplementary Information for "Conserved chromatin and repetitive patterns reveal slow genome evolution in frogs"

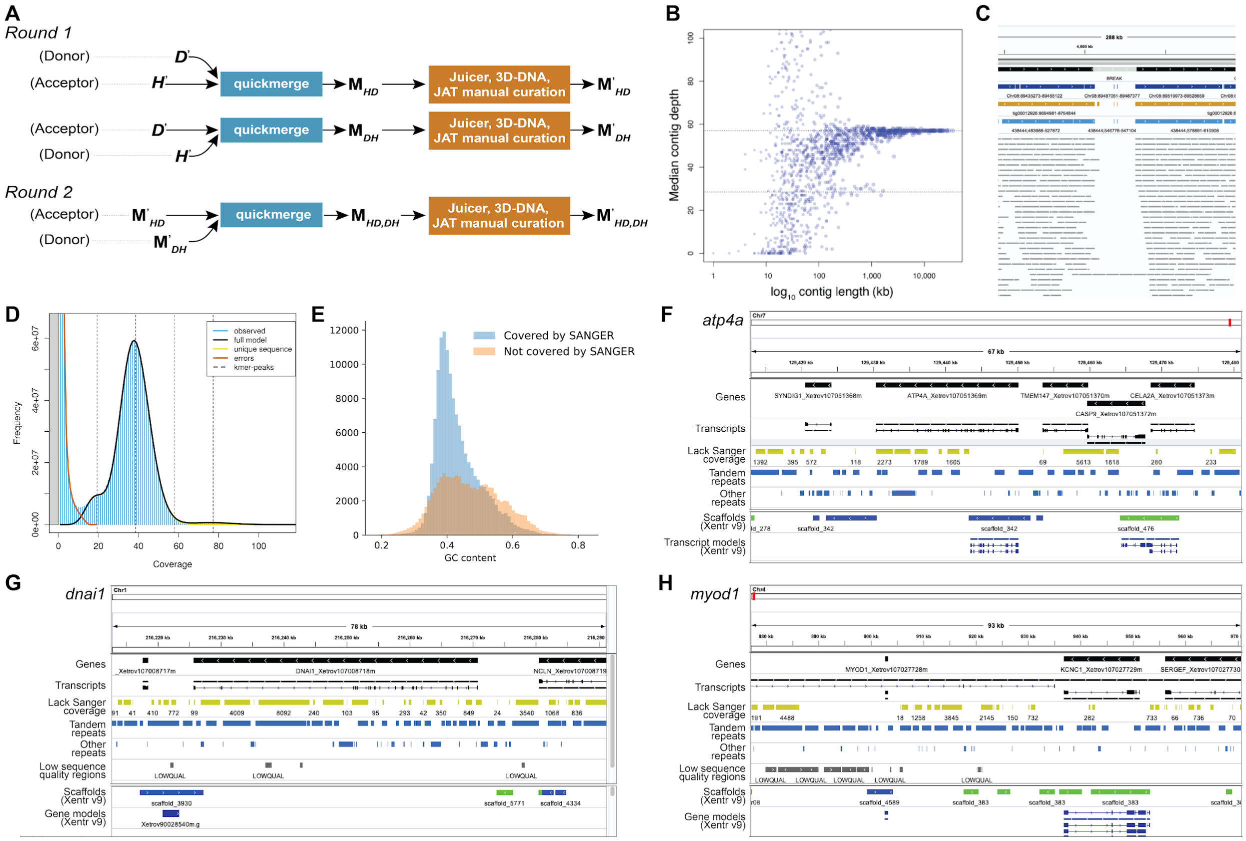

**Supplementary Fig. 1 Genome assembly and recovery of missing genes.**
(**A**) During the first round of metassembly with quickmerge, *de novo* (*D*) and hybrid (*H*) contigs are used as both "donor" and "acceptor" sequences in a reciprocal merging strategy to produce two metassemblies, $M_{HD}$ and $M_{DH}$, which are then corrected for mis-assemblies using the Juicebox Assembly Tools (JBAT) pipeline and manual curation. This results in two sets of corrected contigs, $M'_{HD}$ and $M'_{DH}$. In the second round, the metassembled contigs from the first round are used as inputs to quickmerge, merged together, and corrected as in the first round, producing a final second-order metassembly, $M'_{HD,DH}$. (**B**) The median depth of coverage (Y-axis) was calculated for each contig (blue dots) and plotted against its $log_{10}$-transformed length (X-axis) to identify redundant sequences. The plot shows that contig sequences can be stratified into longer full-depth (y = 58×) and shorter half-depth (y = 29×) categories. (**C**) Black horizontal bars are regions of the genome well-supported by spanning PacBio read alignments (thin grey horizontal bars). The light grey region labeled "BREAK" is poorly supported by read data and represents a DBG2OLC assembly error introduced by incorporating a single PacBio polymerase read that was not successfully broken into individual subreads. Dark blue horizontal bars represent the aligned Sanger-based v9 assembly, while gold and light blue horizontal bars represent the raw Canu and Supernova contigs underlying the hybrid assembly. Note that both these sequences flank the BREAK region, meaning these contigs span the artefact and can be used to patch it. (**D**) GenomeScope2 model fit and genome-size estimate for *X. tropicalis* Nigerian strain female. Observed *k*-mer frequency per depth-of-coverage bin as blue vertical lines; error model curve fit to *k*-mer frequencies resulting from sequencing errors in brown; model fit curve to *k*-mer frequencies generated from unique genomic sequence in yellow; combined unique and error model fit curve in solid black; vertical dashed lines represent one-, two-, three-, and four-copy sequence depth peaks, respectively. (**E**) GC% distribution of genomic regions greater than 100 bp that were covered (*n* = 140,598) or not-covered (*n* = 95,475) by Sanger reads. (**F–G**) IGV views from two fully assembled loci on the current *X. tropicalis* (v10) genome fragmented on the previous genome assembly (v9) due to lack of Sanger read coverage. (**F**) *atp4a* (*Xetrov107051369m*; Chr7:129430384–129450182) that was partially assembled on v9. Note that most regions not covered by Sanger (yellow) overlap repeats (blue). (**G**) *dnai1* (*Xetrov107008718m*; Chr1:216,212,747–216,291,747) was completely missing from v9 and is now captured on the v10 assembly. (**H**) Due to the highly repetitive nature of the sequence surrounding the *myod1* locus (*Xetrov107027728m*; Chr4:719,782–1,128,311) the gene remains fragmented on this assembly. The number below the No-Sanger track corresponds to the size of the fragment not covered by reads.

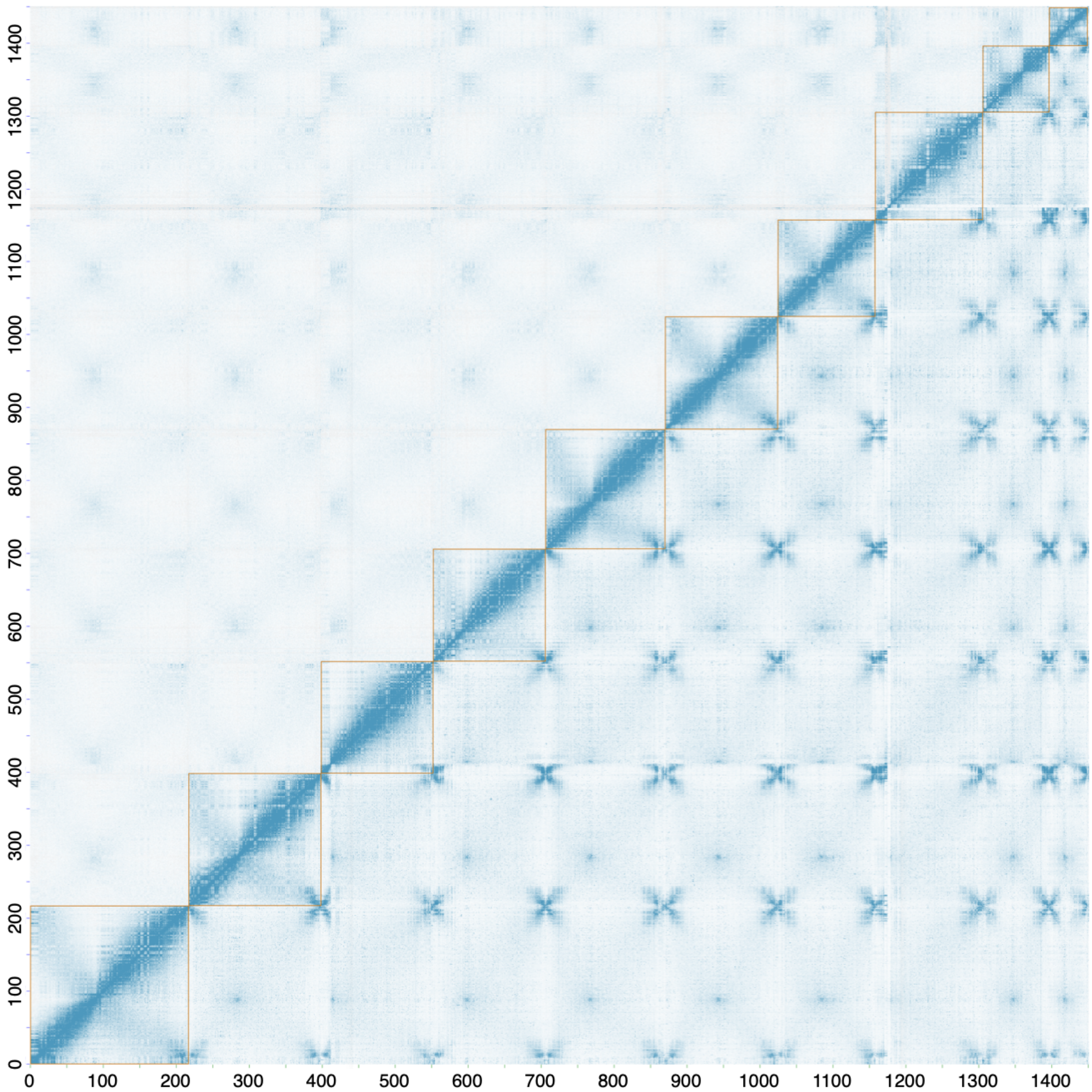

**Supplementary Fig. 2 *Xenopus tropicalis* genome-wide HiC contact map.**
HiC contact matrix from red blood nuclei at 500 kb resolution, balanced using the Knight-Ruiz algorithm, showing reads with a minimum mapping quality of zero (MapQ ≥ 0) below the diagonal and reads with MapQ ≥ 30 above the diagonal. Chromosomes (gold boxes) are shown in ascending numeric order, with p-arms oriented toward the bottom-left of the figure and q-arms toward the top-right. The intensity of blue pixels is proportional to chromatin contact frequencies between X-Y pairs of 500 kb genomic loci. Intra-chromosomal contacts exhibit the highest frequency of contacts between adjacent loci along the linear chromosome (along the diagonal). Above the diagonal, inter-chromosomal contacts are the strongest between centromeres (puncta), while below the diagonal the strongest contacts are observed between subtelomeric sequences.

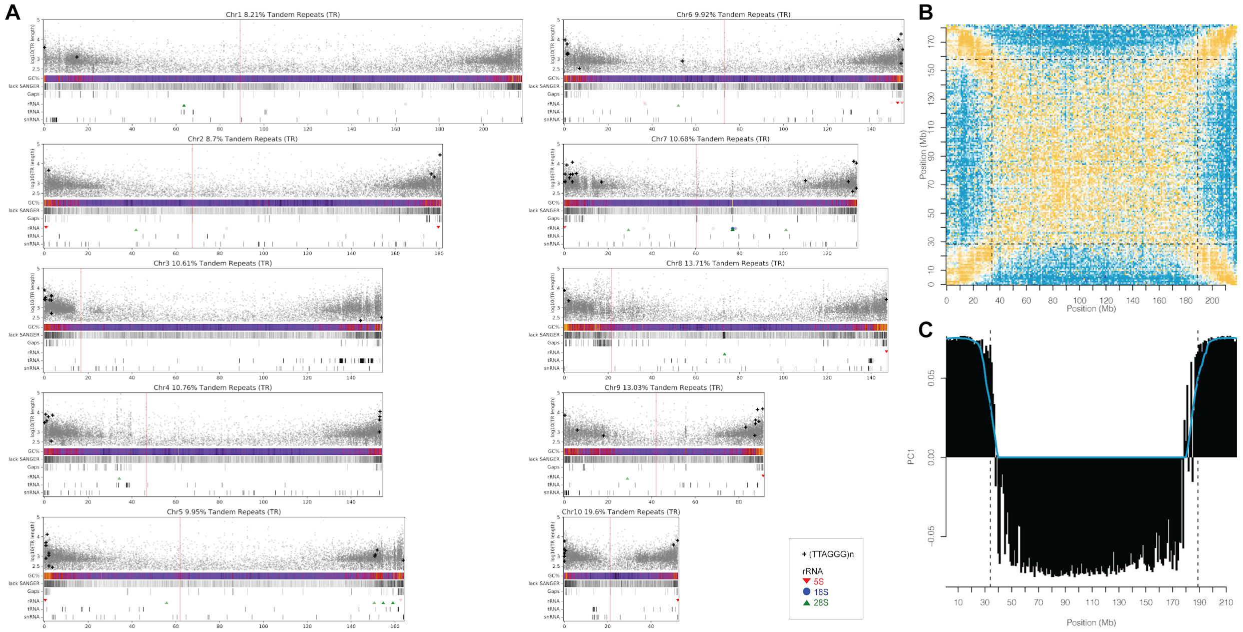

**Supplementary Fig. 3 GC landscape and tandem repeats.**
(**A**) Long arrays of tandem repeats localize to subtelomeric portions of the genome. A total of 68.27 Mb (4.71%) of the *X. tropicalis* genome assembly was not covered by Sanger shotgun sequences. Of the regions lacking Sanger read alignments 48.6% overlapped with tandem repeats, and 34.77% by other types of repetitive elements. Of the 4,718 CDS not supported by Sanger reads, 3,511 (74.41%) of them are localized in close proximity (< 2 kb) to tandem repeats. Elevated GC% content in distal subtelomeric regions is elevated, regions lacking Sanger read coverage are more prevalent in the subtelomeres (48.6% of these regions overlap with tandem repeats). Telomeric-associated tandem repeats (TTAGGG)n are indicated with plus ("+") symbols. Gaps on the current Xentr10 genome assembly tend to co-localize with regions adjacent to long tandem arrays. The locations of clusters of 5S (green), 18S (blue), and 18S (green) rRNA, tRNA, and snRNA are indicated. Red vertical lines indicate the position of the centromere. (**B**) An example of subtelomere boundary inference using feature selection with k-means clustering on a matrix selecting for repetitive HiC read placements in the subtelomeres (yellow, high-density corners), depletion of subtelomere-specific signal with the rest of the chromosome (blue edges), and background/aspecific repetitive signal (center). Dashed horizontal and vertical lines demarcate the inferred subtelomere boundaries. (**C**) Principal component analysis of the matrix in **B**, decomposing variances into subtelomere-specific (positive Y values) and subtelomere-depleted repeat signals (negative Y values). Dashed vertical lines demarcate the inferred subtelomere boundaries.

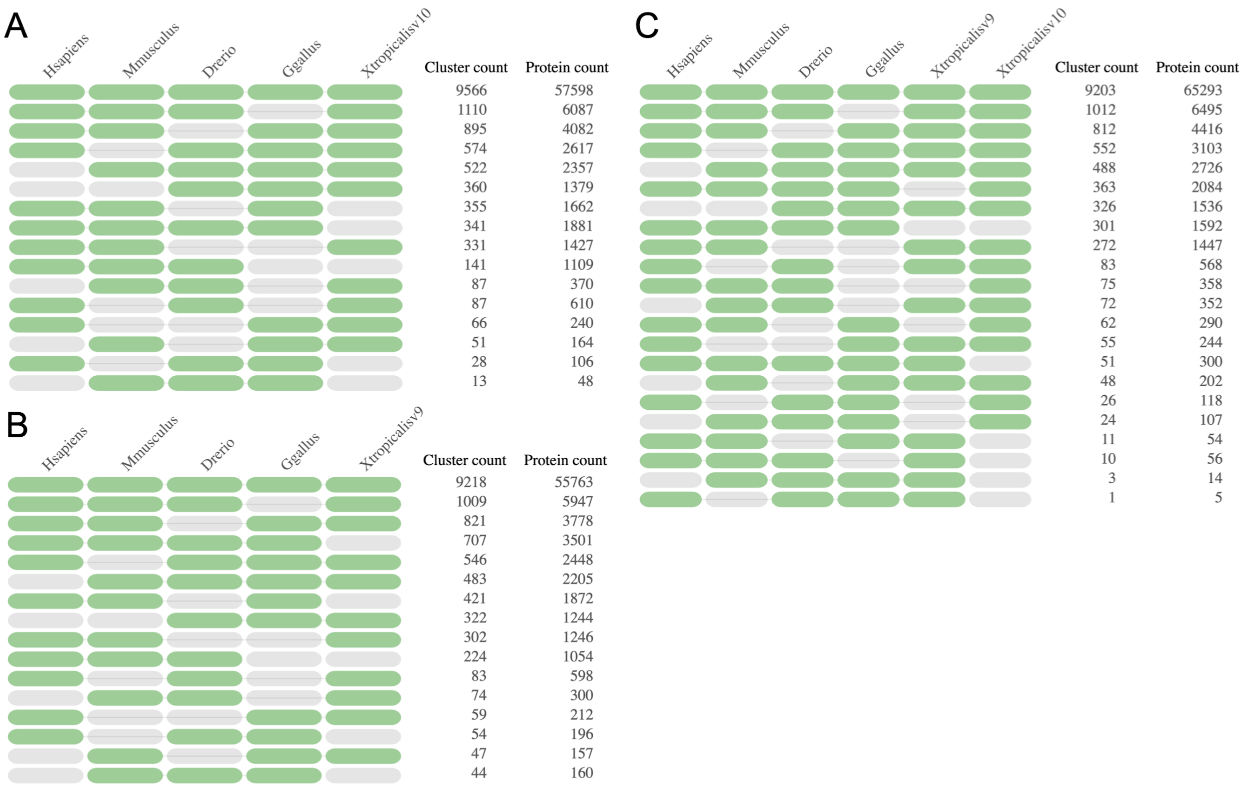

**Supplementary Fig. 4 Comparison of gene content in assemblies of model vertebrates.**
The occurrence tables of gene homology clusters (**A**) containing three or more species with the v10 annotation primary transcripts, (**B**) containing three or more species with the v9 annotation longest transcripts[1] (NCBI Annotation Release 103 of RefSeq assembly accession GCF_000004195.3), and (**C**) containing four or more species with both v9 and v10 against the longest transcripts of *Danio rerio* (NCBI Annotation Release 106 of RefSeq assembly accession GCF_000002035.6), *Gallus gallus* (NCBI Annotation Release 104 of RefSeq assembly accession GCF_000002315.6) *Homo sapiens* (NCBI Annotation Release 109 of RefSeq assembly accession GCF_000001405.39), and *Mus musculus* (NCBI Annotation Release 108 of RefSeq assembly accession GCF_000001635.26). All analyses were completed with OrthoVenn2 (ref.[2]), and the longest transcript amino acid sequences were extracted using gff3ToGenePred and genePredToProt from the UCSC Genomics Institute[3] (binaries downloaded March 5, 2019) as well as custom script largestgenePred.py (v1.0; https://github.com/abmudd/Assembly). In the occurrence tables, the green and grey represent the presence or absence, respectively, of that species in the OrthoVenn2 clustering.

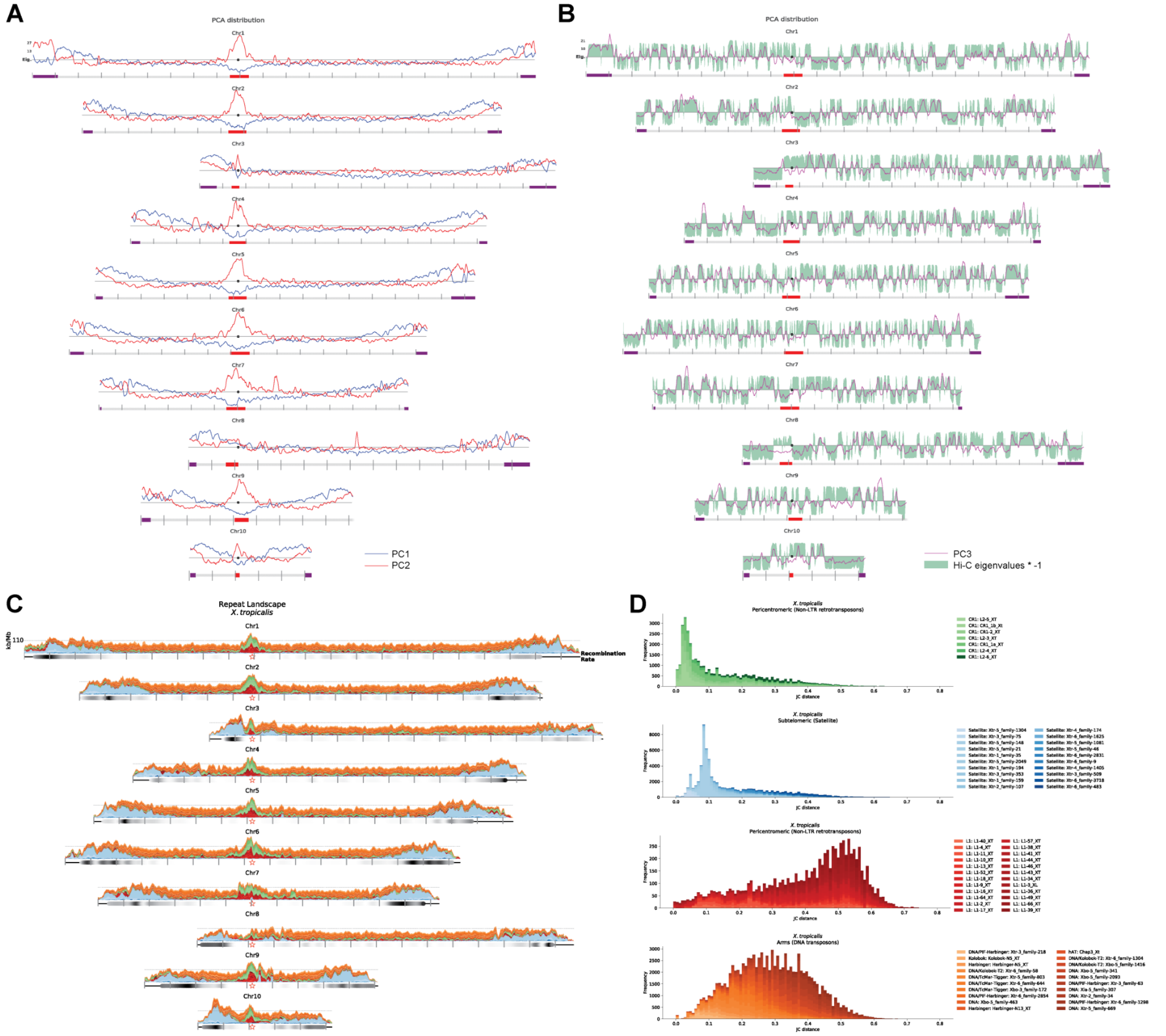

**Supplementary Fig. 5 PCA eigenvectors projected on genomic coordinates.**
(**A**) The first two principal components (PCs) describe subtelomeric and pericentromeric regions of the chromosomes. The first component (PC1, blue lines) strongly correlates with GC content (R = +0.796). The grey horizontal line is the value for which the smoothed PC2 (red lines) was thresholded to discriminate between distal-subtelomeric (purple boxes), pericentromeric (red boxes), and arm regions (grey) of each chromosome. (**B**) The third principal component (PC3, purple lines) is plotted with the eigenvector obtained from the HiC contact matrix that defines A/B compartment structure (green). The PC3 and HiC eigenvectors show a moderately negative correlation (Pearson R = −0.64). The third principal component (PC3) of the repeat density matrix positively correlates with the HiC correlation matrix eigenvectors (R = +0.41). Harbinger-N9_XT is positively associated with PC3 (R = +0.50), whereas DNA/hAT-Ac appears to strongly correlate with PC3 (R = −0.80). Chromosomes are centered by the median position of the Xtr-centromeric associated tandem repeats (black dot). Tick lines correspond to distances of 10 Mb. (**C**) Repeat landscape of L1 (red) and CR1 (green) Non-LTR retrotransposons, Satellite repeats (blue), and DNA transposons (orange). (**D**) Jukes-Cantor (JC) distance from the consensus sequence for the repeats shown on panel **A**. Only the most abundant L1 repeat families are shown on panel **B**.

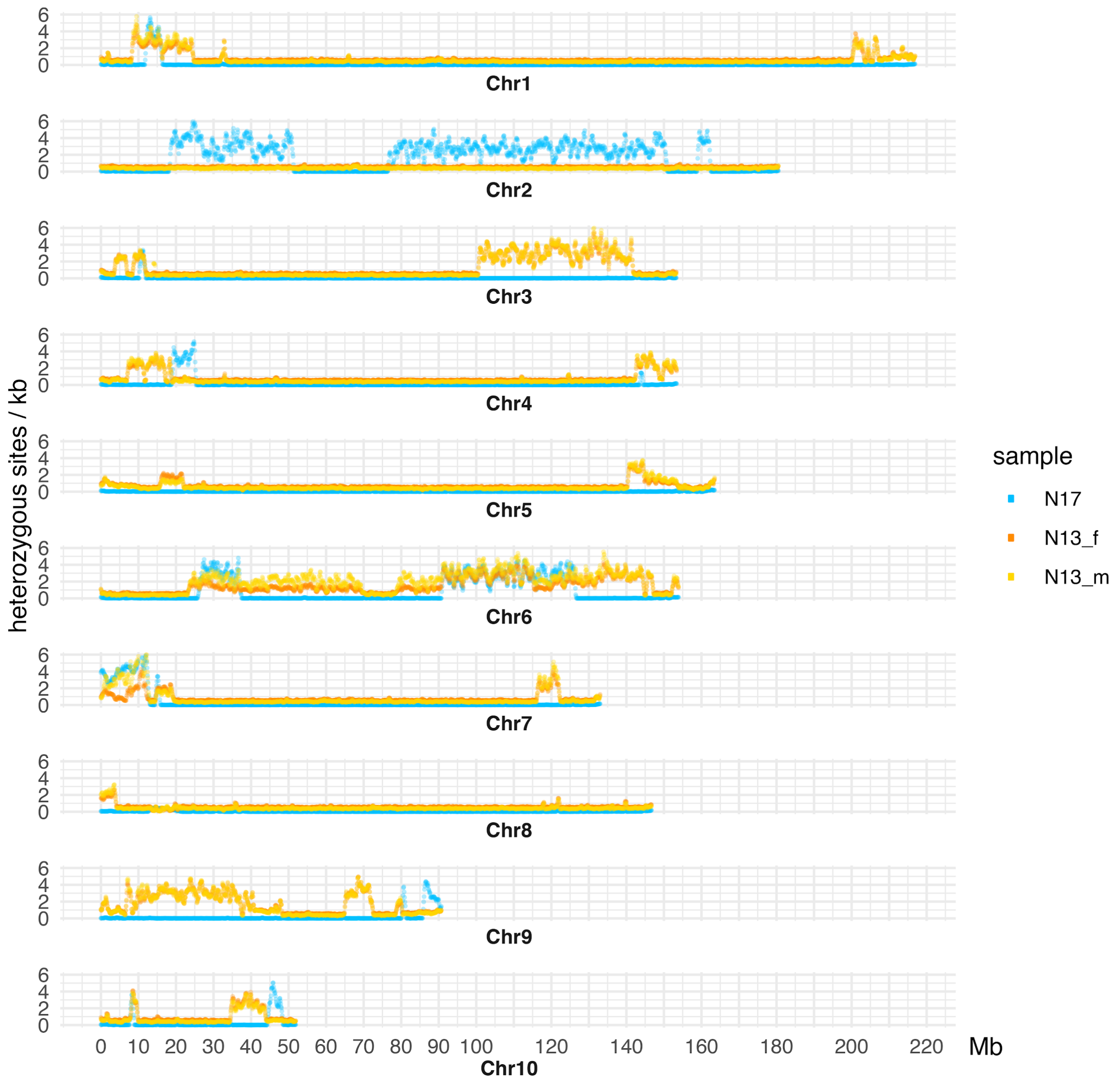

**Supplementary Fig. 6 *Xenopus tropicalis* Nigerian strain residual heterozygosity.**
Heterozygosity rate for the genome frog, N17, and a pool of 8 Nigerian females (N13_f) and 12 Nigerian males (N13_m) from the F13 generation of Nigerian frogs.

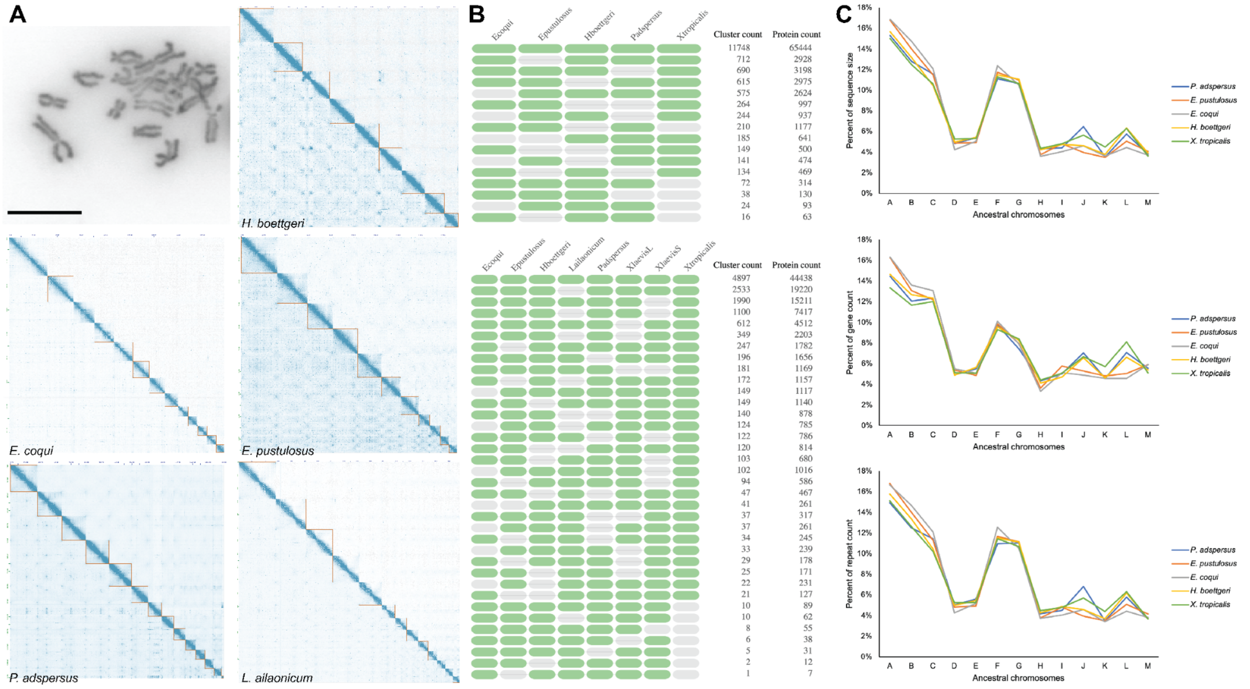

**Supplementary Fig. 7 Assembly and annotation of other frog species.**

(**A**) Nine chromosome pairs are counted for *Hymenochirus boettgeri* metaphase chromosome spread samples, which exhibited both a mode and mean of 2*n* = 18 (75 analyzed images) (top left, scale bar = 10 μm). Whole genome visualization of HiC contact maps from *H. boettgeri*, *Eleutherodactylus coqui, Engystomops pustulosus*, *Pyxicephalus adspersus,* and *Leptobrachium* (*Vibrissaphora*) *ailaonicum* chromosomes. (**B**) The occurrence table of gene homology clusters containing three or more species between the five frog annotations completed in this analysis. (**C**) Percent of total chromosome sequence size for each ancestral unit, gene density in genes per 100 kb for each ancestral unit (middle), and repeat density in repeats per 1 kb for each ancestral unit (bottom). Boundaries of the ancestral units were extracted from runs of collinearity containing at least 1 kb of sequence aligned against *L. ailaonicum*.

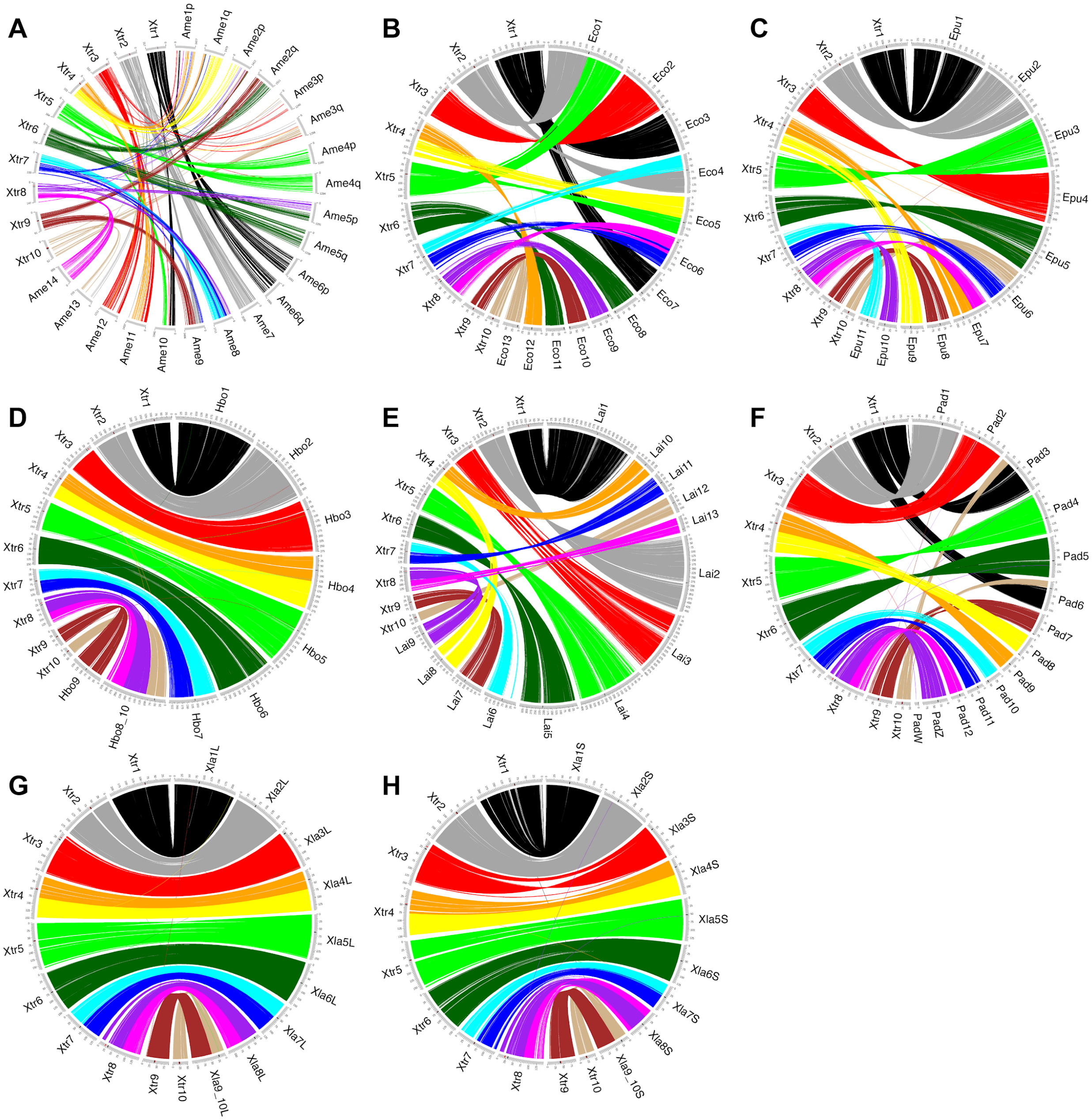

**Supplementary Fig. 8 Pairwise gene colinearity of frog genomes.**
Circos plots with runs of collinearity containing at least 5 kb of aligned sequence between *Xenopus. tropicalis* (left, *Xtr)* and (**A**) *Ambystoma mexicanum* (right, *Ame*), (**B**) *Eleutherodactylus coqui* (right, *Eco*), (**C**) *Engystomops pustulosus* (right, *Epu*), (**D**) *Hymenochirus boettgeri* (right, *Hbo*), (**E**) *Leptobrachium* (*Vibrissaphora*) *ailaonicum* (right, *Lai)*, (**F**) *Pyxicephalus adspersus* (right, *Pad*), (**G**) *X. laevis* L subgenome (right, *Xla*), and (**H**) *X. laevis* S subgenome (right, *Xla*). In panel **A**, *X. tropicalis* and *A. mexicanum* chromosomes are scaled evenly. Runs of collinearity are colored with respect to the 13 ancestral chromosomes (A to M).

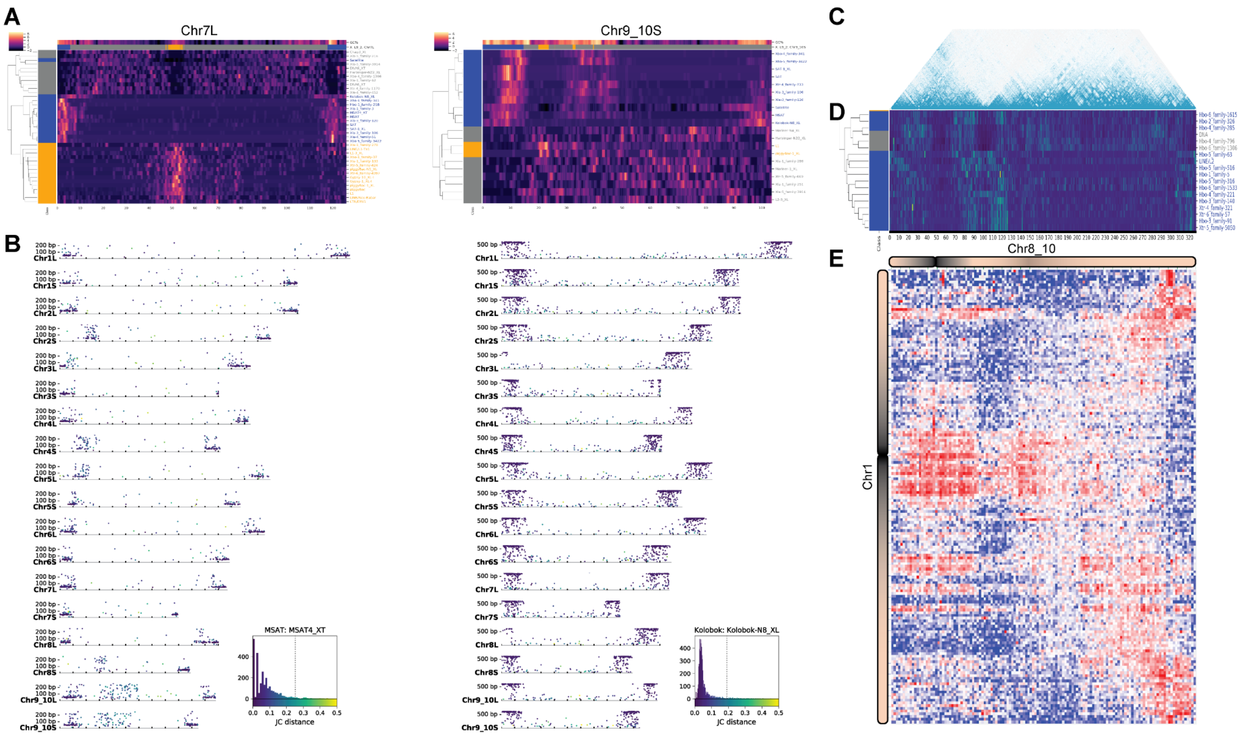

**Supplementary Fig. 9 Chromosome fusions in *Xenopus laevis* and *Hymenochirus boettgeri*.**
(**A**) Heatmap of repeat density in *X. laevis* showing repeats enriched in subtelomeric (blue), pericentromeric (orange), and arm regions (grey) for Chr7L and the fused chromosome Chr9_10S. Three bands of subtelomeric signal can be detected on Chr9_10S, the signal between 30 and 50 Mb corresponds to the fusion of the subtelomeres from ancestral pipid chromosomes Chr9 and Chr10. (**B**) Scatterplots of two subtelomeric repeats showing, for each *X. laevis* chromosome, the length (y-axis) and Jukes-Cantor (JC) distance of the repeats colored as indicated by the histogram on the bottom right. The dotted vertical line on the histogram indicates the 95th percentile. The median JC distances from subtelomeres from all chromosomes (JC = 0.054) is lower than the median JC distances from the region of the Chr9_10 fusion (JC = 0.157), and for the relatively recent p-arm inversion on Chr8S (JC = 0.099) and Chr2S (JC = 0.076), suggesting that the p-arm inversions from Chr8S and Chr2S[4] occurred after the divergence from the pipid ancestor but after the fusion of Chr9_10S. Kolobok-N8_XL, a more recent repeat that expanded post-chromosome fusion and inversion of the p-arm of Chr9S and Chr2S. (**C**) HiC contacts from Chr8_10 chromosome of *H. boettgeri* shows conserved intra-chromosomal contact boundaries compared to ancestral chromosomes Chr8 and Chr10. (**D**) Subtelomeric repeats at the fusion of the ancestral chromosomes between 85 and 120 Mb. (**E**) Enriched centromere-centromere contacts between *H. boettgeri* Chr1 (Y-axis) and Chr8_10 (X-axis), with the strongest centromeric interactions at X, Y = 55 Mb, 170 Mb; suggesting that the active centromere of Chr8_10 was inherited from ancestral chromosome 10. Contacts enriched in the upper-right and lower-right corners represent subtelomere-subtelomere contacts between the two chromosomes. Matrix of observed counts divided by expected counts, at 2.5 Mb matrix resolution (MapQ ≥ 30, Knight-Ruiz balanced).

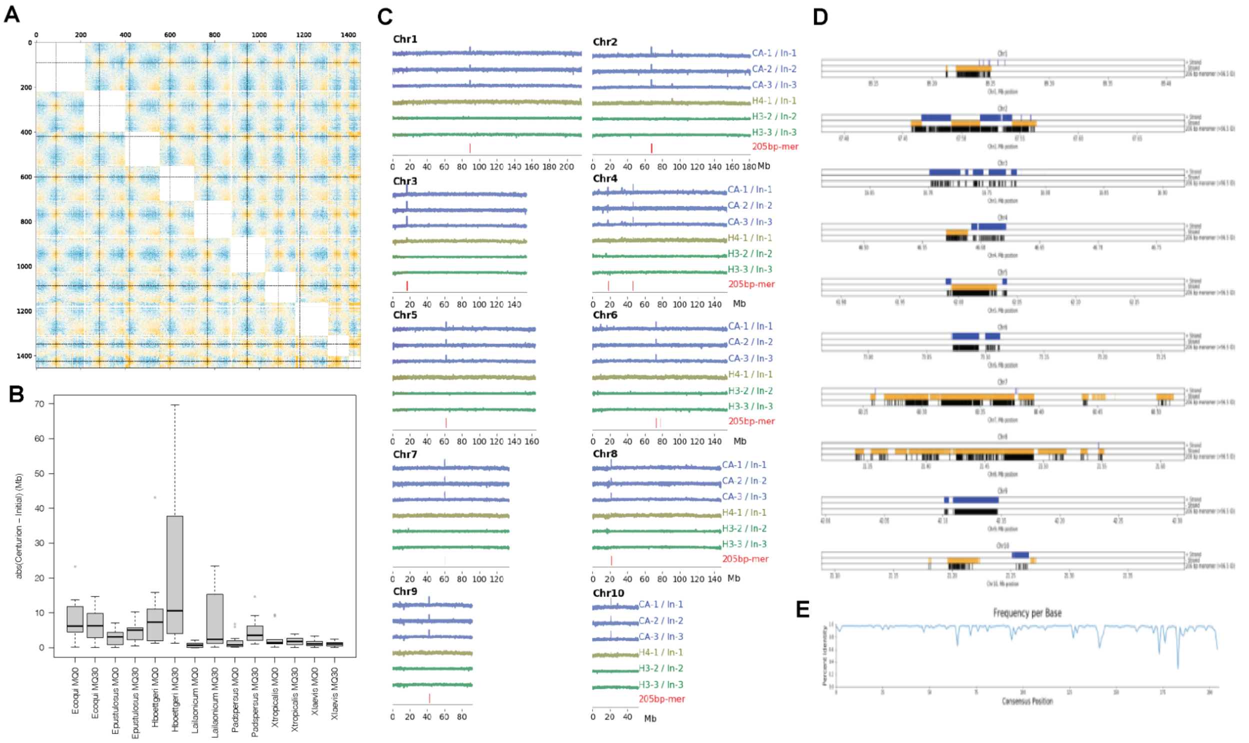

**Supplementary Fig. 10 Estimating the positions of *Xenopus tropicalis* centromeres.**
(**A**) Genome-wide HiC contact map of *X. tropicalis*. Intersecting horizontal and vertical dashed black lines mark the estimated centers of centromeres. Yellow puncta reflect centromere-centromere contact enrichment between chromosomes, suggesting centromere clustering on the inner nuclear envelope. Yellow, white, and blue colors indicate the normalized relative enrichment, parity, and depletion (respectively) between a pair of 1 Mb genomic loci. Intra-chromosomal contacts have been masked (solid white boxes). (**B**) Box plots comparing the variation in differences between manual and Centurion centromere estimates at both MapQ ≥ 0 and MapQ ≥ 30, where each teal point represents an estimate for a single chromosome. Box edges denote the first and third quartiles, while whiskers extend up to 1.5 times the interquartile range. (**C**) ChIP-seq signal for Cenp-a, Histones H4 and H3. Samples are normalized by the read depth of the DNA associated with free-mononucleosomes (input). The presence of 205 bp tandem monomer sequences associated with the Centromeric Tandem Repeats are indicated in red. (**D**) Zoom into the organization of the tandem centromeric repeat sequences (monomer = 205 bp). A 300 kb window surrounding the presumptive centromeric repeats. Monomers found in the forward and reverse strands are shown in blue and yellow, respectively. Monomers sharing above 95% identity to the repeat consensus are shown in black. (**E**) The percentage of the major base per position of the multiple alignment of 3,693 genomic blastn hits from the initial 205 bp monomer sequence. To generate the 205 bp consensus we considered the most represented base per site (positions where alignments were over 50% gaps were omitted).

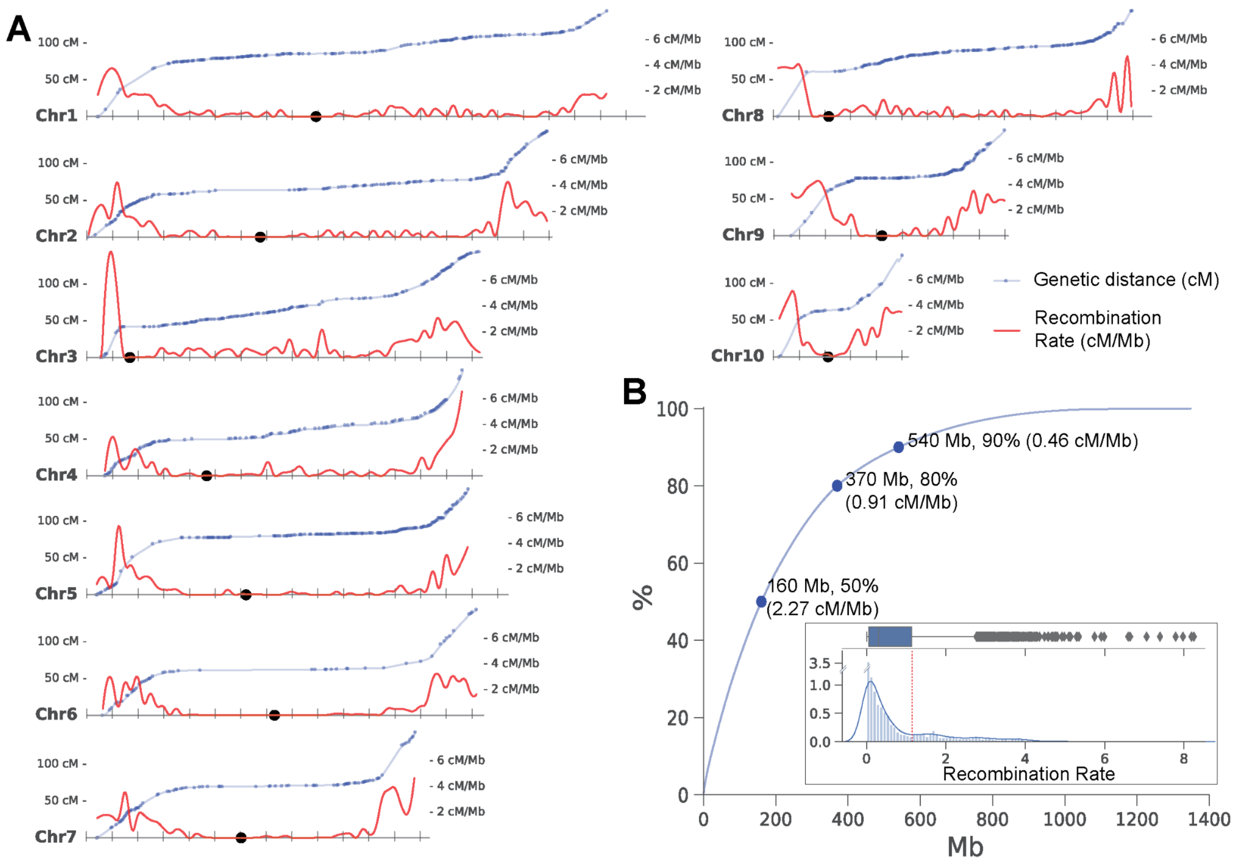

**Supplementary Fig. 11 *Xenopus tropicalis* recombination landscape.**
(**A**) Scatterplot of physical (Mb) vs genetic (cM) position of the genetic map based on the *X. tropicalis* v10 genome assembly. Genetic map distances (blue dots) smoothened by the cubic spline linear interpolation (blue line). Recombination rates (cM/Mb, red line) obtained from the smoothened genetic distances. (**B**) Cumulative distribution of the recombination rate across the whole genome. The distribution of recombination rate (lower histogram) used to determine genetic positions with high and low recombination (75th-percentile, red dotted line).

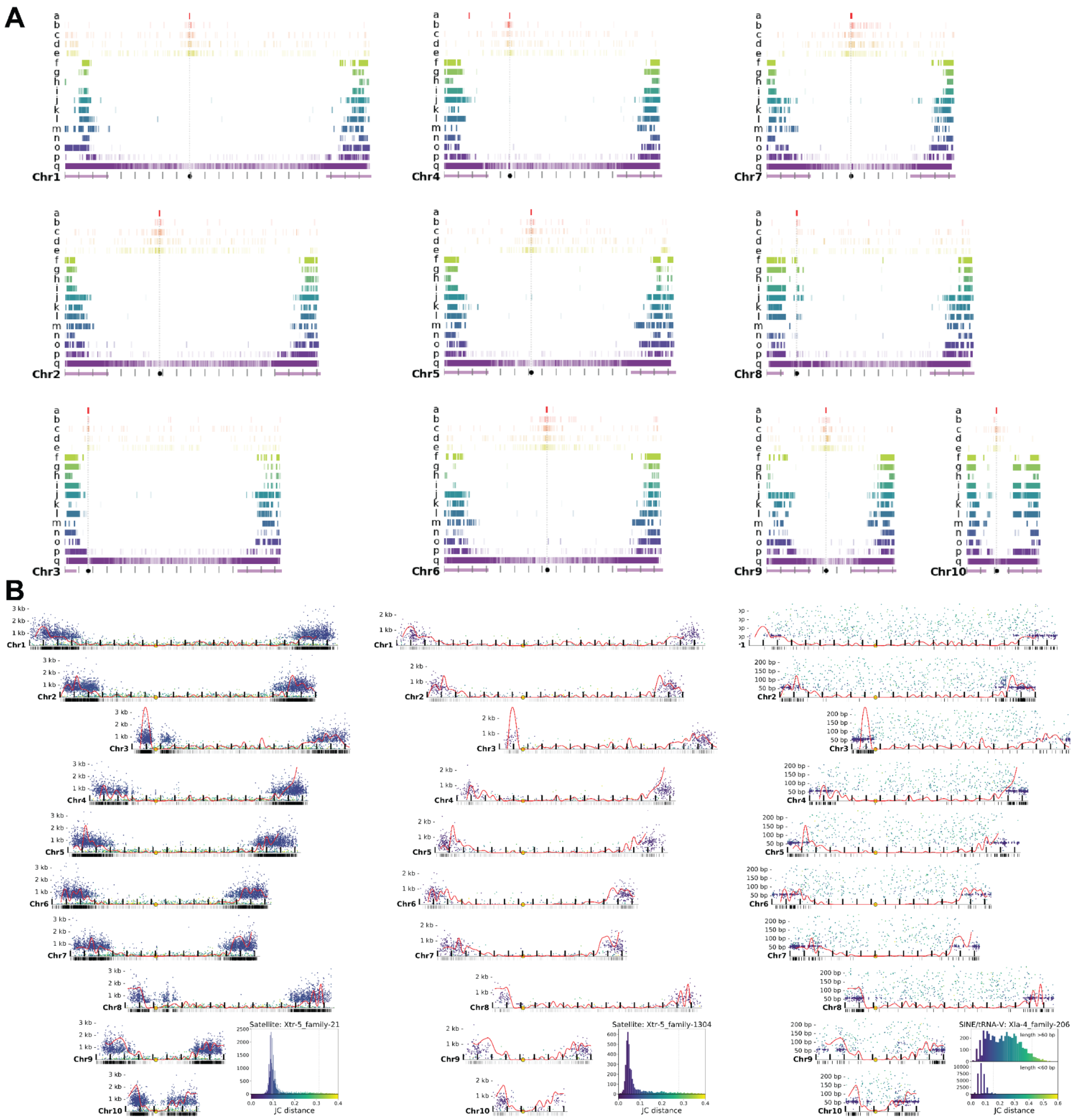

**Supplementary Fig. 12 Distribution of Satellite repeats in *Xenopus tropicalis*.**
(**A**) Distribution of tandem repeats enriched in pericentromeric and subtelomeric regions. Tandem repeat monomers are overrepresented in pericentromeric (tracks *a–e*; red and oranges) and subtelomeric (tracks *f–q*; yellow–purple) regions. Each tick on the graph represents the alignment of a consensus monomer sequence sharing >90% of sequence identity. Sequence track *'a'* (red), 205-bp monomer, is the only monomer exclusive to the pericentromeric regions and referred to as the Centromeric Tandem Repeat (CTR). The median position of the CTRs per chromosome is indicated by the vertical dotted line and the black circle. The dashed vertical line indicates the estimated centromeric position from HiC using stringent mapping parameters. The purple horizontal lines correspond to 30 Mb spanning the subtelomeric domains. Monomer sequences best hit aligned against the repeat database can be found in **Supplementary Table 16**. (**B**) Scatter plot representing repeat length (y-axis) and sequence divergence (Jukes-Cantor distance, color scheme) from satellite families: Xtr-5_family-21 (track q in panel **A**), Xtr-5_family-1304, and a SINE-V/tRNA: Xla-4_family-206 (track p in panel **A**). Long satellite repeats with low sequence divergence localize at subtelomeric portions overlapping areas of high recombination rates (red line). The bottom ticks indicate the presence of the monomer unit of the satellite repeats. The complete sequence of SINE-V/tRNA is relatively uniformly distributed in chromosome arms, except near the pericentromeres and subtelomeres. A minisatellite originated from a portion of the SINE-V/tRNA. (**C**) Histogram of JC distances of SINE/tRNA-V subdivided by sequence length.

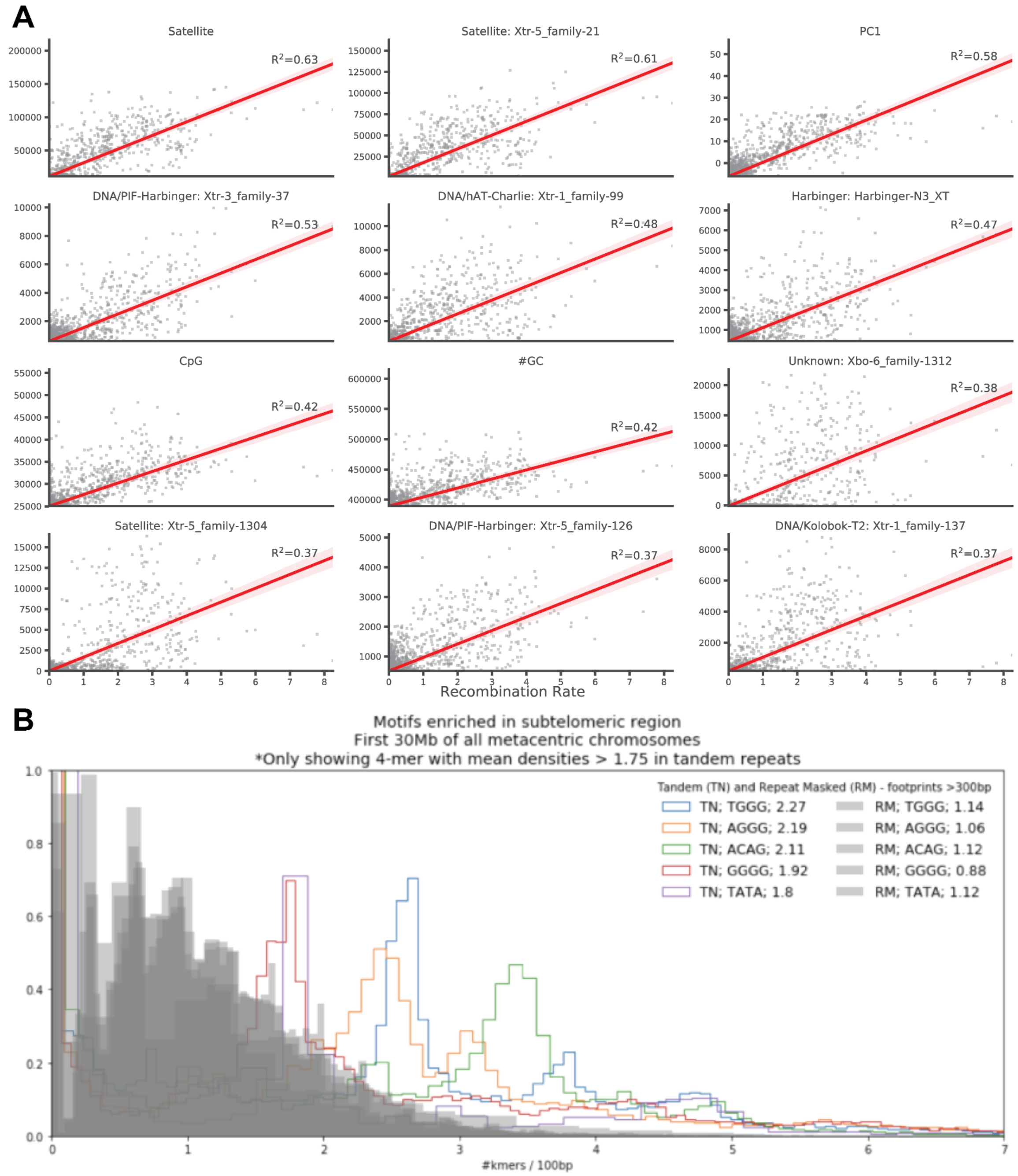

**Supplementary Fig. 13 Correlates of recombination rate.**
(**A**) Scatterplots comparing the genome-wide recombination rates against the top 12 highest correlated genomic features. PC1 corresponds to the eigenvectors of the first principal component obtained from repeat densities. Y-axis is the number of bases per 1-Mb size bins for regions with available genetic markers. A table of all genomic features correlated with recombination rate can be found in **Supplementary Table 14**. (**B**) Enriched tetramers in telomeric sequences. The distribution of tetramers in genomic regions (> 100 bp) overlapping tandem repeat and other non-tandem repeats (grey) in the distal subtelomeric portions of metacentric chromosomes. The two most enriched monomers (AGGG/CCCT and TGGG/CCCA) are similar to the 7-nucleotide oligomer 'CCTCCCT' and 'CCCACCCC' that have been associated with recombination hotspots in human[5] and in mouse[6].

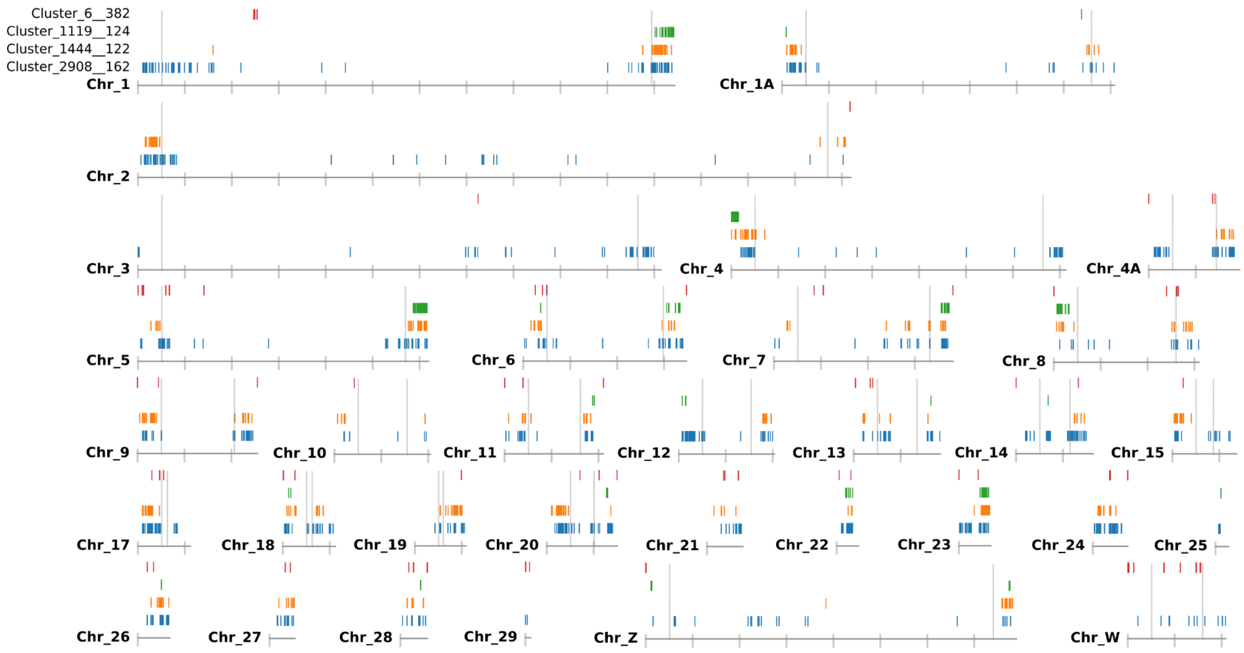

**Supplementary Fig. 14 Zebra finch subtelomeric tandem repeats.**
Genomic distribution of tandem repeats that are enriched in subtelomeric portions of chromosomes (larger than 20 Mb in size). Grey marks denote the first and last 5 Mb where recombination is highest[7]. The tandem repeat sequences appear in at least four of the larger chromosomes and in most microchromosomes.

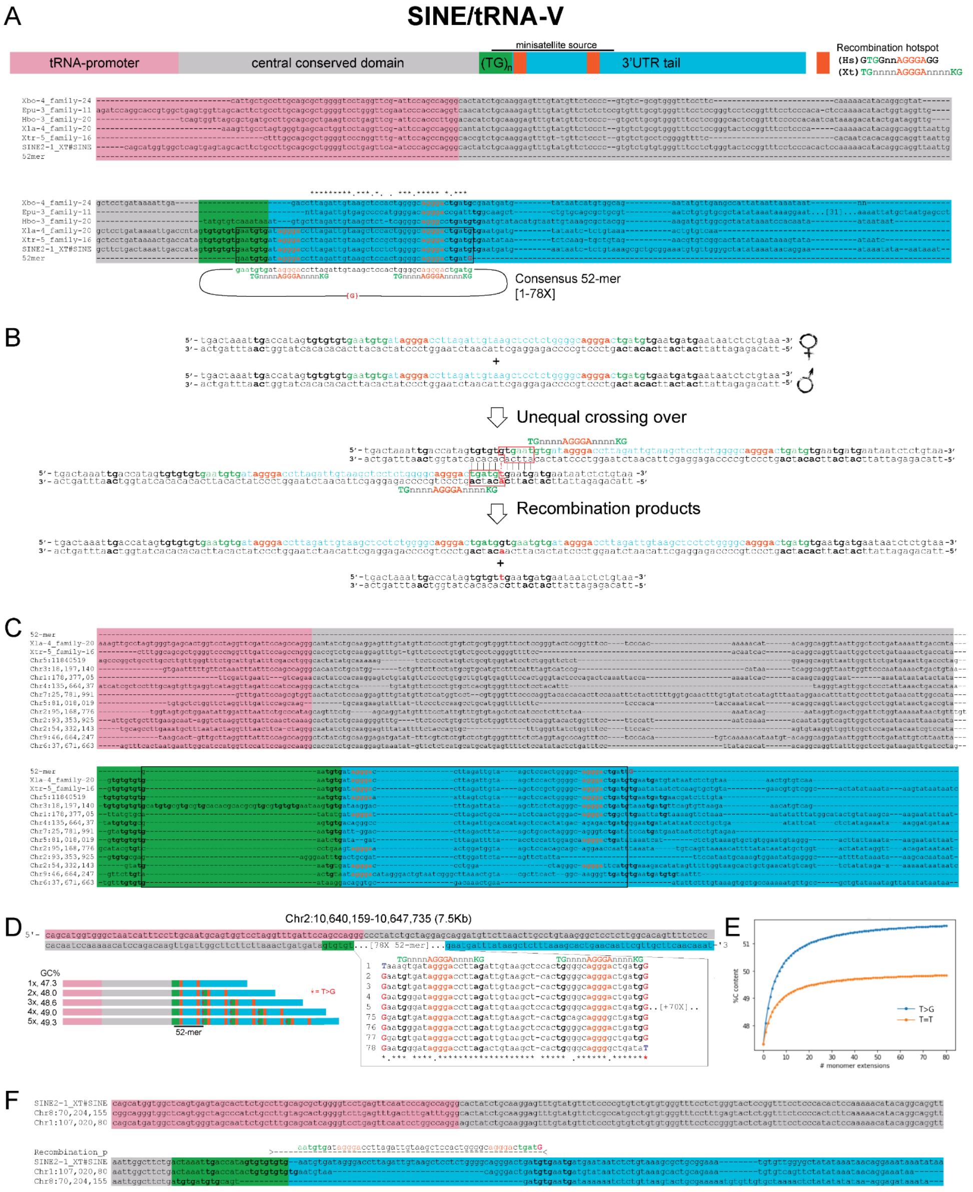

**Supplementary Fig. 15 Microsatellite origin SINE/tRNA evolved into a microsatellite sequence.** (**A**) The structure of *Xenopus* SINE/tRNA (top), as described by Ogiwara et al.[8]. The nucleotide sequence alignment of consensus SINE/tRNA obtained for different frog species (bottom) and the 52-mer sequence where the minisatellite sequence is derived. The multiple sequence alignment of the consensus SINE/tRNA identified for the frogs in this study (except *E. coqui*) shows high levels of sequence similarity. The alignment includes the 52-bp monomer sequence that conforms to the minisatellite. The 52-mer consensus sequence shows perfect alignment with *Xenopus tropicalis* and *X. laevis*. The last base of the monomer is a 'G' substitution of a 'T'. Notice that *E. pustulosus, P. adspersus* and *H. boettgeri* lack the first 'AGGA' box from the consensus sequence. (**B**) Representation of unequal crossing over event between a pair of unequally aligned 3'UTR SINE/tRNA sequences. The unequal crossing over produces a duplication of the 51-mer + 'G' (top) and a deletion of the sequence (bottom). (**C**) An example of the alignment of an intact SINE/tRNA in *X. tropicalis* genome. (**D**) Example of an ancestral SINE/tRNA (Chr2:10,640,159–10,647,735, 7.5Kb) that contains 78 copies of the 52-mer sequence and has extended over 7.5 kb in length. (**E**) The GC% of the consensus SINE/tRNA oscillates between 46 and 48%, while the sequence of the monomer is slightly higher at 51% (51.9% with the addition to the T>G substitution), thus it would be expected that the extension of the tandem would cause a gradual increase in local GC content (see graph on the right). (**F**) Example of the deletion of the 52-mer that possibly resulted from unequal crossing over.

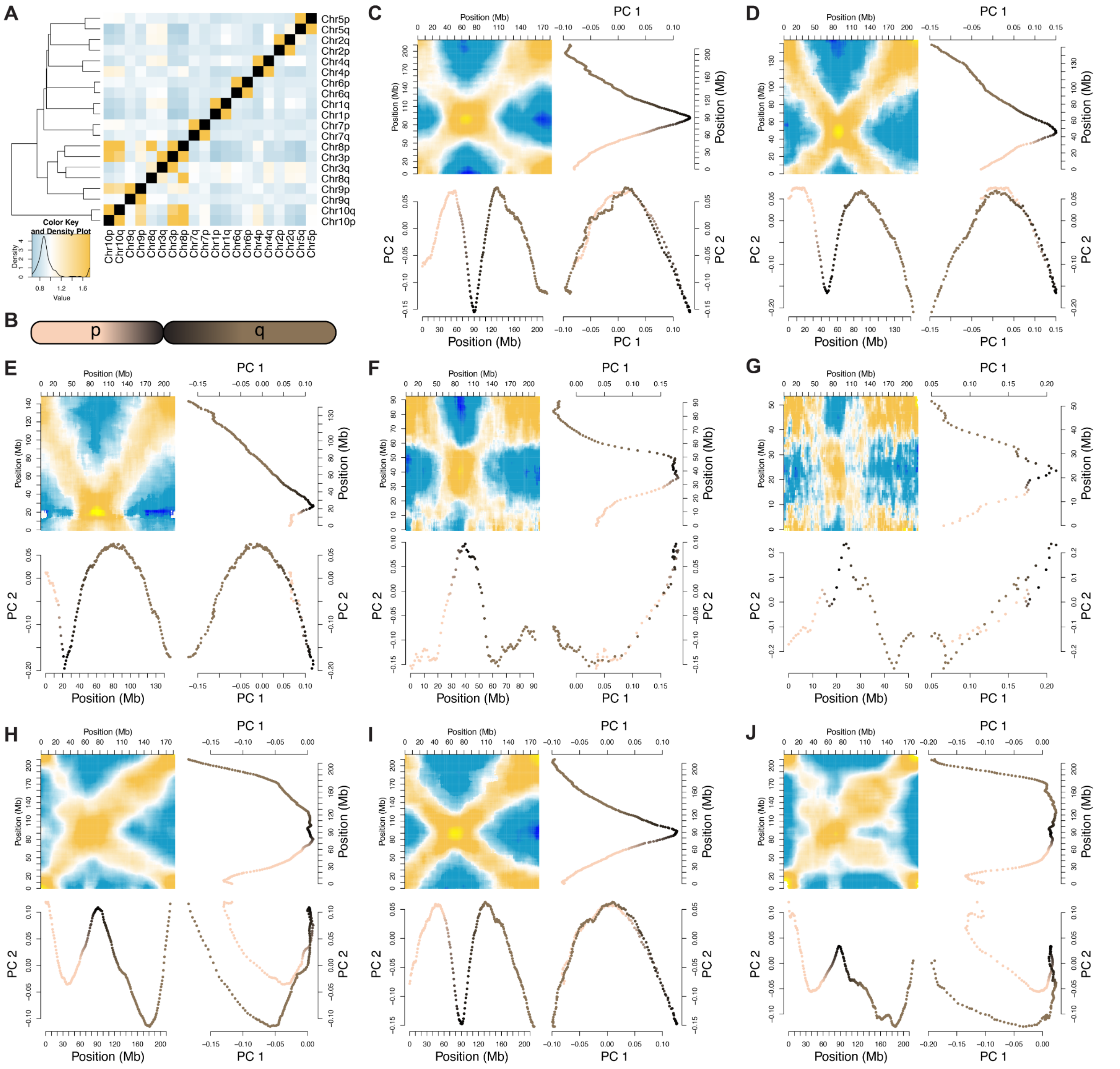

**Supplementary Fig. 16 *Xenopus tropicalis* 3D chromatin structure and nuclear organization.**
(**A**) Genome wide, a non-random association of chromosomal arm-arm contacts is observed (MapQ ≥ 30; $\chi^2$ (361, $n$ = 28,366,570) = 58,25,879.96, $p$-value < $2.2\times10^{-308}$; inter-arm relative range: 0.654–3.829). Chromosome arms are 2.60× enriched for intra-chromosomal contacts over inter-chromosomal contacts ($\chi^2$ (1, $n$ = 4,957,102) = 979,083.44, $p$-value < $2.2\times10^{-308}$). Between chromosomes, p-p and q-q arms contacts are, respectively, 1.077× and 1.0412× enriched over p-q arm contacts ($\chi^2$ (1, $n$ = 24,786,496) = 17,037, $p$-value < $2.2\times10^{-308}$). Note: $2.2\times10^{-308}$ is the lower numerical limit of a signed, double-precision floating-point value on a 64-bit computer. (**B**) In panels **C**–**G**, points are colored tan, black, and brown to differentiate eigenvectors of 1 Mb windows along the p-arm, near the centromere, and the q-arm, respectively; a schematic of this coloring is shown for clarity. Visualizing Rabl chromatin configurations for (**C**) chromosome 1, blood cell nuclei; (**D**) chromosome 4, blood cell nuclei; (**E**) chromosome 8, blood cell nuclei; (**F**) chromosome 9, blood cell nuclei; (**G**) chromosome 10, blood cell nuclei; (**H**) chromosome 1, sperm; (**I**) chromosome 1, NF stage 8; (**J**) chromosome 1, adult brain. For visualization, the contrast level for each HiC contact density map has been adjusted to accommodate each dataset. The target chromosome is situated on the Y-axis of each heatmap, with the comparator chromosome on the X-axis.

**Supplementary Table 1 Sequence Completeness.**

| Statistic | v9 | | v10 | |
|---|---|---|---|---|
| Total assembled scaffold sequence | 1,440,398,454 | bp | 1,451,261,802 | bp |
| Total assembled contig sequence | 1,369,865,365 | bp | 1,448,444,368 | bp |
| Total placed contig sequence | 1,227,064,696 | bp | 1,446,226,505 | bp |
| Total uplaced contig sequence | 142,800,669 | bp | 2,217,863 | bp |
| Contig N50 | 5,552 | | 32 | |
| Contig N50 length | 71,041 | bp | 14,634,335 | bp |
| Contig NG50 | 8,291 | | 41 | |
| Contig NG50 length | 51,188 | bp | 12,113,547 | bp |

Genome statistics comparisons between *X. tropicalis* v9 and v10 assemblies. NG50 calculations assume a genome size of 1.7 Gb[9].

**Supplementary Table 2 Transcript coverage of *X. tropicalis* assemblies v9 and v10.**

| Transcript minimum Coverage (%) | v9 FTS (%) | v10 FTS (%) | v10 SCO (%) |
|---|---|---|---|
| 90.0 | 97.32 | 99.67 | 99.71 |
| 75.0 | 98.12 | 99.80 | 99.83 |
| 50.0 | 98.76 | 99.92 | 99.94 |
| 25.0 | 99.17 | 99.96 | 99.97 |
| 10.0 | 99.38 | 99.96 | 99.97 |
| <10.0 | 0.62 | 0.04 | 0.03 |
| **BUSCO** | **v9 Genome (%)** | **v10 Genome (%)** | **v10 Protein (%)** |
| Complete | 88.5 | 91.7 | 97.0 |
| Single-copy | 87.5 | 90.7 | 95.6 |
| Multi-copy | 1.0 | 1.0 | 1.4 |
| Fragmented | 6.0 | 4.4 | 1.5 |
| Missing | 5.5 | 3.9 | 1.5 |

| Translated MGC minimum coverage (%) | v9 MGC (%) | v10 Translated MGC minimum identity (%) | v10 MGC (%) |
|---|---|---|---|
| 95.0 | 98.18 | 100.0 | 90.99 |
| 80.0 | 99.38 | 95.0 | 99.60 |
| 65.0 | 99.65 | 90.0 | 99.89 |
| 50.0 | 99.86 | 85.0 | 99.96 |
| 35.0 | 99.95 | 80.0 | 99.99 |
| 20.0 | 100.00 | 75.0 | 100.00 |

Minimal coverage and identity percentages of FTS and SCO transcript sets, translated MGC cDNA clones, and BUSCO proteins mapped to the current assembly. BUSCO scores are derived with the BUSCO[10–12] v3.0.2-11-g1554283 pipeline run using tetrapoda_odb9 ($n$ = 3,950). FTS, Full Transcript Set ($n$ = 52,323); SCO, single-copy orthologs ($n$ = 15,613), taken from Session et al.[4], defined using v9 assembly; MGC, Mammalian Gene Collection ($n$ = 8,558).

**Supplementary Table 3** ***Xenopus tropicalis*** **protein coding loci annotation summary statistics.**

| Statistic | Value |
|---|---|
| Annotation version | 10.7 |
| Taxonomy ID | 8364 |
| Primary transcripts (loci) | 25,016 |
| Alternate transcripts | 37,983 |
| Total transcripts | 62,999 |
| **Primary transcripts:** | |
| Average number of exons | 8.7 |
| Median exon length | 137 |
| Median intron length | 1,075 |
| Number of complete genes | 24,167 |
| Number of incomplete genes with start codon | 276 |
| Number of incomplete genes with stop codon | 360 |
| **Gene model support:** | |
| Number of genes with Pfam annotation | 18,895 |
| Number of genes with Panther annotation | 20,727 |
| Number of genes with KOG annotation | 12,109 |
| Number of genes with KEGG Orthology annotation | 10,252 |
| Number of genes with E.C. number annotation | 5,175 |

**Supplementary Table 4 *Xenopus tropicalis* repeat abundances.**

| Repeat type / family | % in Genome | Repeat type / family | % in Genome |
|---|---|---|---|
| **DNA Transposon** | **26.302%** | **LTR Retrotransposon** | **3.312%** |
| hAT | 6.380% | Ty3 | 1.879% |
| Kolobok | 6.048% | DIRS | 0.556% |
| Harbinger | 6.015% | LTR | 0.282% |
| Mariner | 4.057% | Ngaro | 0.211% |
| piggyBac | 1.882% | Endogenous Retrovirus | 0.159% |
| DNA | 1.295% | Copia | 0.105% |
| RC | 0.647% | BEL | 0.071% |
| DNA transposon | 0.522% | Pao | 0.069% |
| CMC | 0.120% | | |
| MULE | 0.062% | **Non-LTR Retrotransposon** | **7.197%** |
| Polinton | 0.023% | LINE | 6.847% |
| Novosib | 0.019% | SINE | 0.351% |
| Sola | 0.009% | SINE? | 0.004% |
| Zisupton | 0.002% | | |
| Merlin | 0.002% | **Other** | **0.228%** |
| Ginger | 0.001% | Transposable Element | 0.188% |
| Crypton | 0.001% | Interspersed repeat | 0.008% |
| Maverick | 0.001% | Inverted repeat | 0.001% |
| | | tRNA | 0.010% |
| **Tandem Repeats*** | **3.817%** | rRNA | 0.011% |
| Satellite | 2.889% | snRNA | 0.010% |
| Simple repeat | 0.932% | Low complexity | 0.093% |

**Supplementary Table 5 Summary of other frog genome assemblies.**

| Genomic feature | *E. coqui* | *E. pustulosus* | *H. boettgeri* |
|---|---|---|---|
| Total scaffold length (bp) | 2,789,403,129 | 2,592,984,374 | 3,214,299,233 |
| Number of scaffolds | 105,233 | 120,236 | 26,522 |
| Scaffold N50 length (bp) | 109,468,876 | 172,109,237 | 293,320,900 |
| Total contig length (bp) | 2,367,745,368 | 2,583,262,543 | 3,210,201,336 |
| Number of contigs | 480,045 | 137,711 | 42,879 |
| Contig N50 length (bp) | 10,801 | 295,193 | 783,846 |
| Contigs sequence in chromosomes (%) | 65.52 | 74.22 | 82.47 |
| Contig GC content (%) | 43.46 | 42.53 | 39.51 |
| Masked contig repeat sequence (%) | 56.10 | 47.92 | 48.60 |

Statistics were calculated using assembly-stats (commit 506a640; https://github.com/sanger-pathogens/assembly-stats).

#### Supplementary Table 6 Summary of annotations for other frog genomes.

| Annotation feature | *E. coqui* | *E. pustulosus* | *H. boettgeri* | *P. adspersus* |
|---|---|---|---|---|
| Number of genes / primary transcripts | 23,346 | 30,613 | 20,684 | 18,673 |
| Number of alternate transcripts | 11,220 | 34,629 | 7,374 | 2,574 |
| Total number of transcripts | 34,566 | 65,242 | 28,058 | 21,247 |
| Average number of exons per gene | 7.3 | 6.5 | 8.5 | 8.8 |
| Median exon length | 137 | 141 | 133 | 133 |
| Median intron length | 1,752 | 1,002 | 1,483 | 1,089 |
| No. of complete genes | 19,899 | 25,989 | 18,244 | 16,861 |
| No. of incomplete genes with start codon | 1,149 | 1,417 | 872 | 637 |
| No. of incomplete genes with stop codon | 1,662 | 2,125 | 1,164 | 941 |
| No. of genes with Pfam annotation | 17,101 | 20,393 | 16,677 | 15,711 |
| No. of genes with Panther annotation | 19,243 | 23,598 | 18,610 | 17,098 |
| No. of genes with KOG annotation | 9,633 | 10,365 | 10,162 | 10,347 |
| No. of genes with KEGG Orthology annotation | 9,799 | 10,528 | 9,973 | 9,779 |
| No. of genes with E.C. number annotation | 4,795 | 6,199 | 4,684 | 4,211 |

Statistics were calculated by the Integrated Gene Call (IGC) pipeline[13].

#### Supplementary Table 7 BUSCO genome scores of other frog genome assemblies.

| BUSCO | *H. boettgeri* | | *P. adspersus* | | *E. pustulosus* | | *E. coqui* | |
|---|---|---|---|---|---|---|---|---|
| | Genome | Protein | Genome | Protein | Genome | Protein | Genome | Protein |
| Complete | 80.0 | 88.4 | 87.5 | 88.6 | 75.7 | 80.2 | 76.4 | 82.5 |
| Single-copy | 77.7 | 85.4 | 86.3 | 86.6 | 72.4 | 76.0 | 75.4 | 80.7 |
| Multi-copy | 2.3 | 3.0 | 1.2 | 2.0 | 3.3 | 4.2 | 1.0 | 1.8 |
| Fragmented | 9.7 | 6.7 | 6.2 | 6.1 | 11.9 | 12.0 | 12.1 | 11.1 |
| Missing | 10.3 | 4.9 | 6.3 | 5.3 | 12.4 | 7.8 | 11.5 | 6.4 |

The following BUSCO scores are derived with the BUSCO[10–12] v3.0.2-11-g1554283 pipeline run using tetrapoda_odb9 ($n$ = 3,950). Percentages are shown for both genome and proteome BUSCO runs.

**Supplementary Table 8 Ancestral chromosome fusions.**

| Species and chromosome | Fusion region | Fused ancestral chromosomes |
|---|---|---|
| *E. coqui* Chr1 | 167,326,341–167,391,435 | B+F |
| *E. coqui* Chr4 | 64,368,134–64,506,892 | H+B |
| *E. coqui* Chr5 | 90,030,948–90,111,922 | E+F |
| *E. coqui* Chr6 | 51,920,009–52,085,381<br>62,174,693–62,190,413<br>74,807,529–74,967,042 | K+I<br>I+K<br>K+I |
| *E. pustulosus* Chr6 | 78,294,430–79,251,447 | M+I |
| *E. pustulosus* Chr7 | 67,080,716–67,193,535 | K+D |
| *H. boettgeri* Chr4 | 132,010,981–132,334,876 | D+E |
| *H. boettgeri* Chr7 | 112,455,236–114,827,918 | H+I |
| *H. boettgeri* Chr8_10 | 103,984,075–104,182,343 | M+J |
| *H. boettgeri* Chr8_10 | 227,189,815–227,531,594 | J+K |
| *P. adspersus* Chr3 | 27,789,267–27,840,023 | M+A |
| *P. adspersus* Chr6 | 25,252,335–25,589,174 | M+A |
| *X. tropicalis* Chr4 | 76,607,359–76,630,690 | D+E |
| *X. tropicalis* Chr7 | 63,487,360–63,537,129 | H+I |
| *X. tropicalis* Chr8 | 81,812,475–81,878,653 | J+K |

Locations of ancestral chromosome fusions in the examined species using runs of collinearity from *L. ailaonicum* containing at least 1 kb of aligned sequence.

**Supplementary Table 9 N50 lengths for collinear runs of orthologous genes between frogs.**

| Species | *L. ailaonicum* | *P. adspersus* | *E. pustulosus* | *E. coqui* | *H. boettgeri* | *X. tropicalis* | *X. laevis* L |
|---|---|---|---|---|---|---|---|
| *P. adspersus* | 90 | – | – | – | – | – | – |
| *E. pustulosus* | 89 | 221 | – | – | – | – | – |
| *E. coqui* | 91 | 119 | 150 | – | – | – | – |
| *H. boettgeri* | 105 | 162 | 168 | 101 | – | – | – |
| *X. tropicalis* | 113 | 175 | 169 | 125 | 323 | – | – |
| *X. laevis* L | 122 | 167 | 186 | 127 | 346 | 1064 | – |
| *X. laevis* S | 111 | 163 | 199 | 116 | 245 | 688 | 676 |

N50 lengths for chromosomal collinear runs of gene orthologs, requiring each run to contain five or more gene orthologs. Calculated with cluster-collinear-bedpe (v0.0.1; https://bitbucket.org/bredeson/artisanal).

#### Supplementary Table 10  Four-fold degeneracy nucleotide divergence.

| Species | *A. mexicanum* | *L. ailaonicum* | *P. adspersus* | *E. pustulosus* | *E. coqui* | *H. boettgeri* | *X. tropicalis* | *X. laevis* L |
|---|---|---|---|---|---|---|---|---|
| *L. ailaonicum* | 1.686 | – | – | – | – | – | – | – |
| *P. adspersus* | 1.704 | 1.039 | – | – | – | – | – | – |
| *E. pustulosus* | 1.730 | 1.065 | 0.753 | – | – | – | – | – |
| *E. coqui* | 1.705 | 1.040 | 0.728 | 0.356 | – | – | – | – |
| *H. boettgeri* | 1.622 | 1.067 | 1.085 | 1.111 | 1.086 | – | – | – |
| *X. tropicalis* | 1.516 | 0.962 | 0.980 | 1.005 | 0.981 | 0.562 | – | – |
| *X. laevis* L | 1.537 | 0.982 | 1.000 | 1.026 | 1.001 | 0.582 | 0.206 | – |
| *X. laevis* S | 1.550 | 0.995 | 1.013 | 1.039 | 1.014 | 0.595 | 0.218 | 0.166 |

Pairwise nucleotide divergence in substitutions per site based on fourfold degeneracy between the examined species extracted from the RAxML[14] (v8.2.11) phylogenetic tree using Newick utilities[15] (v1.6).

#### Supplementary Table 11  Estimation of divergence times.

| Node | TimeTree input minimum | TimeTree input maximum | MEGA7 output estimate |
|---|---|---|---|
| *E. coqui* – *E. pustulosus* | 71 | 108 | 74 |
| *E. coqui* – *P. adspersus* | 147 | 162 | 149 |
| *E. coqui* – *L. ailaonicum* | 167 | 205 | 202 |
| *X. laevis* L – *X. laevis* S | – | – | 38 |
| *X. laevis* L – *X. tropicalis* | 34 | 79 | 52 |
| *X. laevis* L – *H. boettgeri* | 104 | 158 | 126 |
| *X. laevis* L – *E. coqui* | 187 | 220 | 205 |

Divergence time input intervals from TimeTree[16] and estimated output times from MEGA7[17] (v7.0.26) in millions of years ago.

**Supplementary Table 12  Centromeric Associated Tandem Repeat monomer lengths and counts.**

| | *X. tropicalis* centromeric tandem repeat (205 bp monomer) | | | |
|---|---|---|---|---|
| Chromosome | Start | End | Length (kb) | No. monomers |
| Chr1 | 89,211,804 | 89,262,389 | 50.585 | 163 |
| Chr2 | 67,457,118 | 67,564,135 | 107.017 | 512 |
| Chr3 | 16,701,143 | 16,799,031 | 97.888 | 267 |
| Chr4 | 46,570,283 | 46,621,730 | 51.447 | 222 |
| Chr5 | 61,989,064 | 62,041,259 | 52.195 | 227 |
| Chr6 | 73,071,389 | 73,112,569 | 41.180 | 175 |
| Chr7 | 60,256,156 | 60,514,842 | 258.686 | 767 |
| Chr8 | 21,339,510 | 21,551,929 | 212.419 | 834 |
| Chr9 | 42,101,630 | 42,147,670 | 46.040 | 207 |
| Chr10 | 21,179,881 | 21,271,318 | 91.437 | 233 |

**Supplementary Table 13  Mapping statistics for ChIP-seq samples.**

| Sample | Total reads (Millions) | % mapped | % properly paired | % singletons | % mate mapped to diff. chr | Enrichment | | |
|---|---|---|---|---|---|---|---|---|
| | | | | | | CTR region | No-CTR region | CTR / Non-CTR |
| In-1 | 60.72 | 96.93% | 78.68% | 0.83% | 1.46% | 1065.6 | 410.2 | 2.6 |
| In-2 | 43.60 | 96.49% | 77.70% | 1.25% | 1.60% | 834.8 | 292.5 | 2.9 |
| In-3 | 80.31 | 96.53% | 78.21% | 1.24% | 1.14% | 1539.2 | 539.1 | 2.9 |
| Ig-2 | 4.15 | 96.64% | 80.58% | 1.27% | 1.21% | 83.7 | 27.9 | 3.0 |
| Ig-3 | 45.93 | 97.37% | 82.03% | 1.01% | 1.02% | 2246.9 | 742.8 | 3.0 |
| H3-2 | 66.02 | 96.68% | 79.64% | 1.23% | 1.19% | 1371.6 | 443.8 | 3.1 |
| H3-3 | 110.50 | 96.68% | 81.26% | 1.22% | 1.07% | 1145.1 | 311.1 | 3.7 |
| H4-1 | 52.94 | 97.39% | 85.16% | 0.65% | 1.07% | 1613.5 | 359.2 | 4.5 |
| CA-2 | 54.48 | 96.98% | 80.25% | 0.67% | 1.16% | 1600.3 | 92.7 | 17.3 |
| CA-3 | 13.84 | 97.11% | 81.47% | 1.11% | 1.10% | 35698.8 | 1563.8 | 22.8 |
| CA-1 | 233.54 | 97.39% | 82.45% | 1.02% | 1.01% | 9855.7 | 363 | 27.2 |

#### Supplementary Table 14 Correlates of recombination rate.

| Genetic map v1 | Recomb. Rate Correlation | p-value | Median Jungle | Median Desert | Delta | Ratio |
|---|---|---|---|---|---|---|
| Recombination Rate | 1.00 | 0 | 2.511 | 0.146 | 2.37 | 0.89 |
| PCA1 | 0.76 | 0 | 13.768 | −5.885 | 19.65 | 2.49 |
| Satellite: Xtr-5_family-21 | 0.68 | 0 | 49 | 5 | 44.00 | 0.81 |
| Unknown: Xbo-6_family-1312 | 0.67 | 0 | 7 | 0 | 7.00 | 1.00 |
| CpG | 0.66 | 0 | 32934 | 24550 | 8384.00 | 0.15 |
| GC% | 0.65 | 0 | 43.816 | 38.843 | 4.97 | 0.06 |
| Satellite | 0.62 | 0 | 131 | 57 | 74.00 | 0.39 |
| MSAT | 0.59 | 0 | 3 | 0 | 3.00 | 1.00 |
| SINE/tRNA-V: Xla-4_family-206 | 0.58 | 0 | 47 | 3 | 44.00 | 0.88 |
| SINE/tRNA-V | 0.58 | 0 | 51 | 6 | 45.00 | 0.79 |
| CTCTCCC | 0.56 | 0 | 338 | 132 | 206.00 | 0.44 |
| DNA/PIF-Harbinger: Xtr-3_family-37 | 0.56 | 0 | 13 | 4 | 9.00 | 0.53 |
| MSAT: MSAT4_XT | 0.55 | 0 | 2 | 0 | 2.00 | 1.00 |
| CTCF_Hashimoto | 0.54 | 0 | 28 | 11 | 17.00 | 0.44 |
| Unknown: Xtr-4_family-155 | 0.54 | 0 | 4 | 0 | 4.00 | 1.00 |
| CCCCCCC | 0.51 | 0 | 520 | 194 | 326.00 | 0.46 |
| Unknown: Xbo-3_family-280 | 0.47 | 0 | 5 | 0 | 5.00 | 1.00 |
| CpG_ctrl | 0.47 | 0 | 95898 | 88169 | 7729.00 | 0.04 |
| Rec_hotspot | 0.43 | 0 | 193 | 141 | 52.00 | 0.16 |
| Cohesin | 0.42 | 0 | 22 | 8 | 14.00 | 0.47 |
| PCA2 | 0.27 | 1.01E−246 | 4.039 | −4.094 | 8.13 | −147.87 |
| LINE/CR1 | 0.22 | 6.55E−152 | 68 | 57 | 11.00 | 0.09 |
| PCA3 | 0.17 | 3.09E−99 | 0.092 | −2.172 | 2.26 | −1.09 |
| N | 0.10 | 1.59E−31 | 0 | 0 | 0.00 | NaN |
| Ty3/Metaviridae | 0.00 | 0.628721 | 14 | 16 | −2.00 | −0.07 |
| DNA/TcMar-Tigger: Xbo-2_family-40 | −0.10 | 1.71E−32 | 6 | 7 | −1.00 | −0.08 |
| DNA/PiggyBac: Xtr-5_family-424 | −0.10 | 1.40E−35 | 0 | 2 | −2.00 | −1.00 |
| piggyBac: piggyBac-N2_XT | −0.13 | 3.35E−55 | 0 | 2 | −2.00 | −1.00 |
| Harbinger: Harbinger-1_XT | −0.14 | 1.31E−65 | 1 | 2 | −1.00 | −0.33 |
| piggyBac | −0.15 | 1.28E−75 | 11 | 15 | −4.00 | −0.15 |
| DNA_transposon: DNA1_Xt | −0.15 | 1.26E−76 | 13 | 16 | −3.00 | −0.10 |
| piggyBac: piggyBac-N1_XT | −0.18 | 2.32E−103 | 0 | 1 | −1.00 | −1.00 |
| DNA/PiggyBac | −0.19 | 2.67E−122 | 14 | 20 | −6.00 | −0.18 |
| DNA/TcMar-Tigger | −0.20 | 1.61E−127 | 89 | 109 | −20.00 | −0.10 |
| piggyBac: piggyBac-N2A_XT | −0.21 | 5.81E−148 | 0 | 1 | −1.00 | −1.00 |
| DNA_transposon: DNA10_XT | −0.25 | 2.39E−206 | 6 | 10 | −4.00 | −0.25 |
| L1 | −0.26 | 3.47E−228 | 5 | 12 | −7.00 | −0.41 |
| LINE/Penelope | −0.28 | 3.02E−258 | 37 | 48 | −11.00 | −0.13 |
| CR1: CR1_1a_XT | −0.30 | 8.50E−299 | 4 | 7 | −3.00 | −0.27 |
| Helitron: Helitron-N1A_XT | −0.34 | 0 | 0 | 3 | −3.00 | −1.00 |
| DNA_transposon | −0.34 | 0 | 27 | 41 | −14.00 | −0.21 |
| DNA | −0.35 | 0 | 83 | 119 | −36.00 | −0.18 |
| Kolobok: Kolobok-1N3_XT | −0.36 | 0 | 0 | 2 | −2.00 | −1.00 |
| DNA/Kolobok-T2: Xtr-1_family-281 | −0.38 | 0 | 0 | 1 | −1.00 | −1.00 |
| DNA/TcMar-Tigger: Xtr-2_family-11 | −0.38 | 0 | 8 | 15 | −7.00 | −0.30 |
| DNA_transposon: DNA6_XT | −0.40 | 0 | 2 | 6 | −4.00 | −0.50 |
| LINE/CR1: Xtr-4_family-283 | −0.41 | 0 | 2 | 7 | −5.00 | −0.56 |
| Unknown: Xtr-1_family-262 | −0.42 | 0 | 0 | 3 | −3.00 | −1.00 |
| DNA/Kolobok-T2: Xtr-1_family-33 | −0.51 | 0 | 0 | 4 | −4.00 | −1.00 |
| DNA/Kolobok-T2: Xtr-1_family-31 | −0.53 | 0 | 0 | 4 | −4.00 | −1.00 |
| Mariner/Tc1: DNA4_Xt | −0.59 | 0 | 7 | 23 | −16.00 | −0.53 |
| DNA/TcMar-Tigger: Xtr-4_family-702 | −0.61 | 0 | 4 | 21 | −17.00 | −0.68 |
| Pos_Rel_to_end | −0.72 | 0 | 16.167 | 17.784 | −1.62 | −0.05 |

**Supplementary Table 15  Subtelomeric enrichment for tandem repeats.**

| Chromosome | Subtelomere | Non-subtelomere | Ratio |
|---|---|---|---|
| Chr1 | 23.68% | 3.59% | 6.60 |
| Chr2 | 24.27% | 2.76% | 8.79 |
| Chr3 | 26.26% | 3.08% | 8.53 |
| Chr4 | 24.83% | 3.99% | 6.22 |
| Chr5 | 24.17% | 3.72% | 6.50 |
| Chr6 | 23.36% | 3.49% | 6.69 |
| Chr7 | 22.76% | 3.46% | 6.58 |
| Chr8 | 32.63% | 3.99% | 8.18 |
| Chr9 | 20.60% | 3.85% | 5.35 |
| Chr10 | 20.28% | 5.82% | 3.48 |

Sampled from 25.0 Mb from chromosome ends non-overlapping with pericentromeres.

**Supplementary Table 16  Correspondence of monomer sequence with annotated repeat elements.**

| ID | Size | Type | Class/Family | Subfamily | Monomer |
|---|---|---|---|---|---|
| A | 205 | Satellite | Satellite | Centromeric Tandem Repeat | Chr10:21,202,863–21,221,377:206;90.0;206 |
| B | 140 | DNA | PIF-Harbinger | Harbinger-1_XT | Chr4:44,687,900–44,688,471:140;4.1;140 |
| C | 58 | Non-LTR | LINE/CR1 | CR1-2_XT, CR1-1_XL | Chr1:5,339,979–5,340,451:58;8.2;58 |
| D | 134 | DNA | PIF-Harbinger | Xtr-1_family-625, Xtr-1_family-181 | Chr9:43,007,496–43,007,890:132;2.9;134 |
| E | 55 | DNA | PiggyBac | Xtr-4_family-150, Xtr-6_family-62 | Chr6:78,080,043–78,080,155:55;2.1;55 |
| F | 57 | N/A | N/A | N/A | Chr7:130,271,437–130,276,838:57;94.8;57 |
| G | 104 | N/A | N/A | N/A | Chr10:15,393,501–15,394,099:104;5.8;104 |
| H | 112 | Non-LTR | CR1 | Xtr-5_family-2393, Xla-1_family-417 | Chr2:177,493,031–177,494,141:112;9.9;112 |
| I | 82 | N/A | N/A | N/A | Chr7:3,459,827–3,460,032:84;2.5;82 |
| J | 135 | Satellite | MSAT | MSAT1_XT | Chr10:16,832,997–16,834,248:135;9.3;135 |
| K | 71 | N/A | N/A | N/A | Chr1:214,792,050–214,792,755:71;9.9;71 |
| L | 100 | Satellite | SAT | SAT-1_XL | Chr4:142,973,443–142,974,440:100;9.9;100 |
| M | 132 | N/A | N/A | N/A | Chr1:210,557,533–210,557,838:135;2.3;132 |
| O | 61 | N/A | N/A | N/A | Chr6:74,371,563–74,371,758:61;3.2;61 |
| P* | 52 | Non-LTR | SINE/tRNA-V | SINE2-1_XT | Chr3:13,767,515–13,767,949:52;8.5;52 |
| Q | 113 | Satellite | Satellite | DNA4Sat_Xt, Xtr-5_family-21 | Chr10:15,491,890–15,492,956:113;9.4;113 |

*See Supplementary Table 17.

**Supplementary Table 17 Copy counts of 52-mer minisatellite.**

| Species | Count |
|---|---|
| *X. tropicalis* | 41,014 |
| *X. laevis* | 17,375 |
| *N. parkeri* | 3,738 |
| *H. boettgeri* | 305 |
| *L. ailaonicum* | 128 |
| *E. pustulosus* | 46 |

Number of Blast hits of 52-mer monomer sequence derived from SINE2/tRNA in frog genome assemblies.

**Supplementary Table 18 Quantification of Rabl structure strength and significance.**

| Chr | Blood cell nuclei | Sperm† | Stage 8† | Stage 9† | Stage 10† | Stage 11† | Stage 12† |
|---|---|---|---|---|---|---|---|
| 1 | 0.213** | 0.864** | 0.078** | 0.125** | 0.136** | 0.157** | 0.155** |
| 2 | 0.031** | 0.875** | 0.022** | 0.010** | 0.009** | 0.004** | 0.003** |
| 3 | 0.233** | 0.265** | 0.402** | 0.524** | 0.586** | 0.364** | 0.273** |
| 4 | 0.101** | 0.596** | 0.057** | 0.194** | 0.159** | 0.103** | 0.100** |
| 5 | 0.017** | 0.675** | 0.013** | 0.013** | 0.008** | 0.008** | 0.013** |
| 6 | 0.185** | 0.553** | 0.064** | 0.059** | 0.060** | 0.062** | 0.067** |
| 7 | 0.076** | 0.546** | 0.042** | 0.026** | 0.043** | 0.038** | 0.044** |
| 8 | 0.027** | 0.362** | 0.141** | 0.135** | 0.143** | 0.066** | 0.053** |
| 9 | 0.163** | 0.914 | 0.499** | 0.411** | 0.491** | 0.674** | 0.676** |
| 10 | 0.421 | 0.635 | 0.061 | 0.076 | 0.076 | 0.038 | 0.062 |
| **Sum** | **1.465** | **6.285** | **1.376** | **1.573** | **1.711** | **1.514** | **1.447** |

| | Stage 13† | Stage 15† | Stage 17† | Stage 23† | Liver† | Brain† |
|---|---|---|---|---|---|---|
| 1 | 0.202** | 0.202** | 0.211** | 0.176** | 0.952** | 1.260** |
| 2 | 0.008** | 0.018** | 0.020** | 0.015** | 0.687** | 0.885** |
| 3 | 0.341** | 0.246** | 0.223** | 0.080** | 0.113** | 0.179 |
| 4 | 0.105** | 0.098** | 0.112** | 0.050** | 0.354** | 1.077** |
| 5 | 0.012** | 0.025** | 0.032** | 0.037** | 0.584** | 0.432 |
| 6 | 0.058** | 0.056** | 0.063** | 0.068** | 0.454** | 0.745** |
| 7 | 0.049** | 0.050** | 0.056** | 0.067** | 0.070** | 0.123 |
| 8 | 0.045** | 0.042** | 0.042* | 0.036* | 0.031** | 0.203* |
| 9 | 0.412** | 0.379** | 0.348** | 0.443** | 0.170** | 1.146 |
| 10 | 0.074 | 0.096 | 0.117 | 0.118 | 0.559 | 0.441 |
| **Sum** | **1.308** | **1.213** | **1.224** | **1.088** | **3.975** | **6.490** |

†Data from Niu et al.[18].
Numerical values reported below are the residual sum of squares (RSS) estimates for strength of Rabl structure. Double-asterisks ("**") label chromosomes with *p*-values ≤ $1\times10^{-3}$, single-asterisks ("*") as *p*-values ≤ $1\times10^{-2}$, and chromosomes with *p*-values > $1\times10^{-2}$ are left unlabeled.

#### Supplementary Table 19 Contact enrichment between chromosomes and chromosome arms.

| Enrichment between chromosome pairs | Degrees of Freedom (df) | Sample size (N) | Chi-squared statistic | *p*-value | Relative Enrichment Min. | Relative Enrichment Max. |
|---|---|---|---|---|---|---|
| *E. coqui* | 144 | 6,918,603 | 769,032.76 | < 2.23E−308 | 0.714 | 1.602 |
| *E. pustulosus* | 100 | 21,963,827 | 2,650,230.54 | < 2.23E−308 | 0.853 | 1.313 |
| *H. boettgeri* | 64 | 11,472,000 | 1,544,949.63 | < 2.23E−308 | 0.874 | 1.190 |
| *L. ailaonicum* | 144 | 57,076,729 | 7,466,579.43 | < 2.23E−308 | 0.680 | 1.723 |
| *P. adspersus* | 121 | 128,641,903 | 13,700,811.18 | < 2.23E−308 | 0.882 | 1.201 |
| *P. adspersus** | 169 | 144,706,271 | 14,272,071.75 | < 2.23E−308 | 0.626 | 18.763 |
| *X. laevis* | 289 | 23,901,183 | 1,519,869.44 | < 2.23E−308 | 0.924 | 1.281 |
| *X. tropicalis* | 81 | 24,987,749 | 3,049,787.53 | < 2.23E−308 | 0.828 | 1.168 |

| Enrichment between chromosome arms | Degrees of Freedom (df) | Sample size (N) | Chi-squared statistic | *p*-value | Relative Enrichment Min. | Relative Enrichment Max. |
|---|---|---|---|---|---|---|
| *E. coqui* | 625 | 8,199,375 | 6,186,850.95 | < 2.23E−308 | 0.256 | 10.999 |
| *E. pustulosus* | 441 | 26,317,421 | 11,057,282.37 | < 2.23E−308 | 0.610 | 5.842 |
| *H. boettgeri* | 289 | 14,196,106 | 7,422,295.55 | < 2.23E−308 | 0.510 | 6.145 |
| *L. ailaonicum* | 625 | 71,298,515 | 54,254,909.53 | < 2.23E−308 | 0.317 | 7.842 |
| *P. adspersus* | 529 | 143,487,710 | 27,537,888.69 | < 2.23E−308 | 0.618 | 4.401 |
| *P. adspersus** | 729 | 160,073,454 | 29,474,072.39 | < 2.23E−308 | 0.626 | 18.763 |
| *X. laevis* | 1225 | 26,203,393 | 7,391,101.40 | < 2.23E−308 | 0.579 | 5.781 |
| *X. tropicalis* | 361 | 28,366,571 | 5,825,879.96 | < 2.23E−308 | 0.654 | 3.830 |

| Contact enrichment within chromosomes over between chromosomes | Degrees of Freedom (df) | Sample size (N) | Chi-squared statistic | *p*-value | Relative enrichment |
|---|---|---|---|---|---|
| *E. coqui* | 1 | 1,634,268 | 691,927.92 | < 2.23E−308 | 4.725 |
| *E. pustulosus* | 1 | 5,644,909 | 2,131,647.86 | < 2.23E−308 | 4.188 |
| *H. boettgeri* | 1 | 3,486,504 | 1,215,445.75 | < 2.23E−308 | 3.883 |
| *L. ailaonicum* | 1 | 17,073,360 | 8,945,125.49 | < 2.23E−308 | 6.242 |
| *P. adspersus* | 1 | 21,829,747 | 4,806,499.22 | < 2.23E−308 | 2.768 |
| *P. adspersus** | 1 | 22,157,873 | 5,585,252.08 | < 2.23E−308 | 3.017 |
| *X. laevis* | 1 | 3,332,189 | 1,136,530.06 | < 2.23E−308 | 3.808 |
| *X. tropicalis* | 1 | 4,957,102 | 979,083.44 | < 2.23E−308 | 2.600 |

| p-p and q-q arm contact enrichment over p-q arm contacts | Degrees of Freedom (df) | Sample size (N) | Chi-squared statistic | *p*-value | Relative enrichment |
|---|---|---|---|---|---|
| *E. coqui* | 1 | 6,850,547 | 3,914.92 | < 2.23E−308 | 0.928 |
| *E. pustulosus* | 1 | 21,760,538 | 4,752.74 | < 2.23E−308 | 1.032 |
| *H. boettgeri* | 1 | 11,423,575 | 8,322.19 | < 2.23E−308 | 1.061 |
| *L. ailaonicum* | 1 | 56,582,771 | 8,462.43 | < 2.23E−308 | 1.026 |
| *P. adspersus* | 1 | 127,451,200 | 3,059.42 | < 2.23E−308 | 1.011 |
| *P. adspersus** | 1 | 143,432,205 | 160.96 | = 9.99E−37 | 1.002 |
| *X. laevis* | 1 | 23,564,270 | 59,815.60 | < 2.23E−308 | 1.119 |
| *X. tropicalis* | 1 | 24,786,496 | 17,037.87 | < 2.23E−308 | 1.059 |

Asterisked rows include sex chromosomes in calculations.

**Supplementary Table 20 Relative enrichment of HiC contacts between chromosomes.**

| ***X. tropicalis*** | **Chr1** | **Chr2** | **Chr3** | **Chr4** | **Chr5** | **Chr6** | **Chr7** | **Chr8** | **Chr9** |
|---|---|---|---|---|---|---|---|---|---|
| Chr2 | 1.090 | – | – | – | – | – | – | – | – |
| Chr3 | 1.069 | 1.018 | – | – | – | – | – | – | – |
| Chr4 | 1.058 | 1.017 | 0.999 | – | – | – | – | – | – |
| Chr5 | 1.052 | 1.069 | 0.997 | 1.001 | – | – | – | – | – |
| Chr6 | 1.044 | 1.016 | 0.984 | 1.000 | 1.010 | – | – | – | – |
| Chr7 | 1.010 | 0.992 | 0.971 | 0.981 | 0.990 | 0.999 | – | – | – |
| Chr8 | 1.041 | 0.999 | 1.052 | 0.984 | 0.978 | 0.968 | 0.954 | – | – |
| Chr9 | 0.952 | 0.952 | 0.916 | 0.952 | 0.958 | 0.981 | 0.983 | 0.929 | – |
| Chr10 | 0.828 | 0.876 | 0.878 | 0.943 | 0.890 | 0.962 | 1.000 | 0.971 | 1.168 |

### Supplementary Methods

#### Supplementary Note 1: High-throughput sequencing, *Xenopus tropicalis*

##### High molecular weight DNA isolation

DNA was isolated from blood cells of an *X. tropicalis* $F_{17}$ Nigerian strain female using a method similar to that described in Mitros et al. (2019). Briefly, blood was collected from a benzocaine-anesthetized frog into a 15 mL polypropylene tube filled with 0.85× SSC. The tube was capped and inverted to mix, and spun down at 900 *g* for 5 min. The supernatant was removed by decanting. Cells were resuspended in 10 mL 0.85× SSC, and spun down again at 500 *g* for 5 min. The supernatant was removed and cells were resuspended in 10 mL lysis buffer (1% SDS, 20 mM EDTA, 100 mM NaCl, 20 mM Tris pH 7.5–8.0; supplemented with 200 μg/ml Proteinase K) and incubated overnight at 55°C. Phenol extraction was performed with an equal volume of buffered phenol and mixed very gently end-over-end overnight, then spun at 900 *g* for 5 min. DNA was precipitated from the aqueous phase by addition of 1/10 vol 3 M ammonium acetate, thorough mixing, then the addition of 0.6 vol isopropanol, then more thorough mixing. DNA was spooled with an end-closed Pasteur pipet and scraped into a tube of 70% EtOH, then spooled again into a clean tube and all traces of EtOH removed. DNA was resuspended in 1 mL of TE pH 8.0. This DNA was used for PacBio, 10x, and Illumina WGS sequencing libraries.

##### PacBio SMRT sequencing

DNA from the $F_{17}$ Nigerian strain female was sent to HudsonAlpha Institute for Biotechnology for library preparation and sequencing. A total of 4.83 million SMRT sequencing reads were generated using a PacBio RSII with P6-C4 chemistry, resulting in 65.4 Tb and an estimated

38.5× depth of coverage. Reads were filtered for lengths of 3 kb or longer; half of the sequenced bases were captured in reads 18.7 kb or longer.

#### 10x Chromium linked reads

We generated 366.2 million pairs (65× sequencing depth) of Illumina 2×151 bp sequences on a HiSeq X Ten at HudsonAlpha from DNA from an $F_{17}$ Nigerian strain female frog. Average molecule length from 1.36 million Chromium GEMs was inferred to be 55.3 kb, with 84.8% of molecules estimated to be 20 kb or longer, and 20% of molecules 100 kb or longer.

#### In vivo, high-throughput chromatin conformation capture (HiC)

Three HiC libraries were constructed by Dovetail Genomics LLC. from blood and sequenced by the QB3 Vincent J. Coates Genomics Sequencing Laboratory (VGCSL) at the University of California, Berkeley for 381,335,524 total pairs (**Supplementary Data 1**).

#### Shotgun sequencing

Whole-genome shotgun mate-pair Sanger sequencing of the $F_7$ female Nigerian strain used in this study is described by Mitros *et al.*[1]. Additionally, a 670 bp-insert library (JBL052) was prepared by the Functional Genomics Laboratory at UC Berkeley from the female $F_{17}$ Nigerian strain genomic DNA using the KAPA HyperPrep Kit. We then generated 192.5 M pairs (58× depth of coverage) of 2×251 bp Illumina whole-genome shotgun sequencing reads on a HiSeq 2500 at the QB3 VCGSL.

### Supplementary Note 2: *Xenopus tropicalis* genome assembly and annotation

#### *De novo* and hybrid contig assemblies

Two independent PacBio long-read contig assemblies were constructed; the first contig dataset was assembled *de novo* using Canu[19], the second using the hybrid genome assembler, DBG2OLC[20].

Canu (v1.6-132-gf9284f8) assembled 1,561 Mb of total sequence, with half of the assembled bases represented in ($n$ = 91) contigs 3.5 Mb or longer. Canu was invoked with the following parameters: genomeSize=1.7g minReadLength=5000 minOverlapLength=2000.

DBG2OLC (commit 1f7e752) hybrid contigs were assembled by incorporating 10x Genomics Supernova-assembled contigs (described below) with the raw/uncorrected PacBio long-read sequences. Parameter sweeps to maximize assembly contiguity and total assembled sequence determined that k 17 KmerCovTh 2 MinOverlap 20 AdaptiveTh 0.005 RemoveChimera 1 MinLen 3000 yielded the optimal assembly for the data. DBG2OLC parameters were varied in all combinations over the following ranges: KmerCovTh, 2–10; MinOverlap, 10–150; and AdaptiveTh, 0.0001–0.02. The optimal contig assembly was polished twice with PBDAGCON[21] (v0.3) using reads aligned by BLASR[22] (v5.3) as input (PBDAGCON params: -t 0 -c 0; BLASR params: --bestn 1 --nproc 8 --minAlnLength 2500 --minPctSimilarity 70). These contigs represent 1,429 Mb of total assembled sequence, with half of the assembled bases in ($n$ = 174) contigs 2.4 Mb or longer.

The Supernova[23] (v1.1.5) assembly used in the hybrid assembly above was generated by HudsonAlpha using default parameters and output in “pseudohaploid1” mode. Of the 380 million read pairs input to Supernova, 35% of the reads passed its internal QC procedures and were used in assembly. This resulted in an effective depth of 30.4× in sequence coverage. Supernova scaffolds were broken into contigs at scaffolding gaps prior to inputting to DBG2OLC. These Supernova-assembled scaffolds represent 1,365 Mb of total sequence, 1,204 Mb of which is captured in contig sequence (N50 length = 18.3 kb, N50 count = 17,057).

The 10x Genomics linked-reads were later reassembled with Supernova (v2.0.1) using default parameters and output in “pseudohaploid1” mode, resulting in a more complete assembly: 1,507 Mb total scaffold sequence and 1,327 Mb total contig sequence, with an N50 length of 20.5 kb (N50 count = 16,433).

#### Contig meta-assembly

Both *de novo*- and hybrid-assembled contig datasets were used as quickmerge[24] (commit e4ea490) “donor” and “acceptor” inputs in a two-way hierarchical contig merging strategy. This process was motivated by observing complementary overlaps between the contig sets when compared using nucmer[25] (MUMmer v3.23). The merging strategy is illustrated for clarity in **Supplementary Fig. 1A** and described in detail below, where *“D*” and *“H*” represent *de novo* (Canu) and hybrid (DBG2OLC) assemblies, respectively.

Prior to merging, contig extension errors in the initial *D* and *H* contig sets were first broken. These mis-assemblies were detected by performing a preliminary round of scaffolding on each contig set with Juicer (v1.5.6-37-gd3ee11b) and 3D-DNA[26,27] (commit 2796c3b), identified via visualization in Juicebox[28] (v1.9.0), and manually broken with Juicebox Assembly Tools[29] (JBAT). Merging was then performed in two rounds:

*Round 1*: Corrected *D* and *H* contig sets (*D*' and *H*') were merged (-hco 16 -c 5 -l 100000 -ml 10000) using alignments generated by nucmer (default parameters) and filtered with delta-filter[25] (-q -r -i 95). However, the outcome of merging is asymmetric and depends on contig set input order (i.e., their “acceptor” or “donor” status). To exhaust potential merges and maximize the amount of metassembled sequence, reciprocal metassemblies were performed. This resulted in $M_{DH}$ and $M_{HD}$ metassembled contig sets, where subscripts list which contig sets were used as acceptor and donor in each merge, respectively. These initial metassembled contig sets were then corrected for merge errors with JBAT, as was performed above.

*Round 2*: Correcting the reciprocal metassemblies above created two contig sets, called $M'_{DH}$ and $M'_{HD}$, each accomplishing a subset of the total potential merges (a result of the asymmetric nature of the quickmerge algorithm). However, the unrealized potential merges in Round 1 could be exploited by a second round of merging to produce a single, optimal metassembly, $M_{HD,DH}$. Finally, this metassembly was corrected of merge errors, as was performed above, generating a final metassembly ($M'_{HD,DH}$) with a contig N50 length of 7.7 Mb (N50 = 48) and capturing 1,453.4 Mb of total sequence.

#### Removing redundant contig sequences

Upon aligning the short-read Illumina sequences to the metassembled contigs, we examined the median depth profiles for each contig (**Supplementary Fig. 1B**) and observed an abundance of contigs with approximately half the depth expected from sequencing, as well as many contigs that have near-zero depth. The latter contigs were discarded as assembly artifacts, while the former were aligned all-vs.-all to the entire metassembled contig set using nucmer[25] (MUMmer v3.23; default parameters). After removing self-mapped alignments, the shorter of two contigs that aligned over at least 90% of their length at 90% identity (as these were yet-unpolished sequences) or greater were discarded (redund-contigs; https://bitbucket.org/bredeson/artisanal). This process was repeated until half-depth sequences could no longer be purged from the contig set. If a half-depth sequence could be identified as redundant to another, larger contig in JuiceBox[28], it was also removed. The final non-redundant assembly converged to 1,453 Mb, with 127 Mb of duplicate/repetitive sequences removed.

#### Recovering genic contigs

Because the process for removing redundant sequences was aggressive, it had the potential to remove legitimate contigs that partially overlapped (i.e., were partially redundant with) another, larger contig yet still contained uniquely assembled sequence. To assess the degree with which this happened and recover lost sequences, we aligned all 52,323 *X. tropicalis* transcript CDS sequences available at XenBase (http://ftp.xenbase.org/pub/Genomics/JGI/Xentr9.1) to the filtered metassembled contigs using GMAP[30] (version 2019-03-15; --npaths=0 --min-identity=90 --format=gff3_match_cdna) and minimap2[31] (v2.5-284-g1739a26; -c -x splice --secondary=no). Of those sequences, 131 were found to be mitochondrial or other contaminant and were excluded from further analysis. Among the 52,192 remaining CDS sequences, only 356 (0.68%) could not be found with minimum thresholds of 90% identity and 50% sequence coverage. These unmapped sequences were then used to probe (in the following order) the Canu, DBG2OLC, Supernova (both v1.1.5, then v2.0.1), v9, and v4 assemblies to recover gene-containing contigs that were excluded previously or that were not assembled at all. In total, 315 CDS sequences were recovered and 41 (0.079%) could not be (see the “Evaluating assembly completeness and correctness” section).

#### Mate-pair, HiC, and synteny-based scaffolding

Merged and error-corrected contigs from the metassembly procedure were scaffolded using SSPACE[32] v3.0 with 3 kb, 8 kb, and 40 kb mate-pair libraries, and 140 kb BAC-end libraries[1]. Reads were downloaded from the NCBI Trace Archive (SPECIES_CODE = "XENOPUS TROPICALIS" and CENTER_NAME = "JGI" and SOURCE_TYPE = "GENOMIC") and their 5' ends trimmed to remove low-quality (baseQ < 20) bases. Reads were then truncated to a max length of 750 bases and aligned to the metassembly with BWA-MEM[33] (v0.7.17-r1188) independently as single-end reads. Read alignments were paired and filtered for proper pair orientation and insert distance (3 kb, 8 kb, 40 kb libraries, within 6 SDs from the mean; 140 kb BAC-ends, within 4 SDs from the mean) using custom scripts (repair and scaffold-read-filter.py). Pairs with mapQ ≥ 30 were converted to TAB format and used for scaffolding with SSPACE v3 (-Z 6 -k 2 -n 1e12).

Mate-pair scaffolded contigs were ordered and oriented into chromosomal scaffolds with 3D-DNA (commit 2796c3b) using 108.9M HiC contacts (mapQ ≥ 1) extracted by Juicer[26,27] (v1.5.6-37-gd3ee11b). The HiC scaffolded assembly was manually curated in the Juicebox[28] (v1.9.0) visualizer with Juicebox Assembly Tools[29] (JBAT) to optimize the adjacency of high-density inter-contig contacts along the main diagonal. This was performed by identifying and correcting the HiC contact patterns created by scaffolding errors, such as those described by Dudchenko et al.[29]. Additionally, contigs not placed into scaffolds by 3D-DNA were scaffolded manually if sufficient signal was observed.

The synteny with a preliminary PacBio- and HiC-based *X. laevis* assembly (in preparation) was used to refine ambiguities in contig order and orientation, particularly within subtelomeric regions of the *X. tropicalis* assembly, where contigs tended to be small and the HiC contact densities may have been too sparse or signal-to-noise too low. Contigs were aligned using nucmer (MUMmer v3.23) (default parameters), filtered using delta-filter (-q -r -o 10), then visualized using mummerplot. Contigs were ordered and oriented along the chromosomes using layout files (via -Q and -R options). Of the 20 chromosome ends, four telomeric sequences were captured in the assembly, and are capped by (TTAGGG)n repeats (**Supplementary Fig. 3A)**. The final chromosome-scale scaffolding is presented in **Supplementary Fig. 2**.

#### Gap closing

Gaps introduced by HiC scaffolding are of unknown size and were assigned a fixed length of 1,000 bp. A fixed-length gap was re-sized if it could be spanned by two or more Sanger mate pairs (aligned as described in the previous section), with both ends in a pair with mapping quality 30 or greater. Gaps were then re-sized using a custom script (mpGapLen, -q1 -k3 -S -Z6, https://bitbucket.org/bredeson/artisanal). Given the orders and orientations of contigs within chromosomal scaffolds, we were able to resolve with greater specificity overlaps between segments of chromosome created by premature contig extension termination. These overlaps constitute gaps of negative length in the assembly. For a potential negative gap between two contigs to be closeable, contigs were required to overlap by at least 1,000 bp and align at 95% sequence identity; the shorter of the two overlapping sequences was then trimmed and the two contigs concatenated together. This procedure closed 80 of 1,248 gaps in the chromosomes. To close positive gaps, we performed two rounds of PBJelly[34] (PBSuite v15.8.24) gap filling, re-sizing gaps after each round as described above, and closed 478 gaps. An additional six gaps were closed via polishing with Pilon (see next section).

#### Assembly Polishing

Canu and DBG2OLC assemblies were evaluated for short-range structural correctness with custom assembly error detection software called WOMBAT (https://gitlab.com/Bredeson/wombat; parameters: -A3 -a1 -m10000 -l10000 -W2000 -w100 -S500 -s50) using PacBio read alignments. Common errors identified included collapsed multi-copy sequences and artifactual sequences introduced by the DBG2OLC assembly process (**Supplementary Fig. 1C**). Sequences identified to be artifactual were hard masked with Ns and the Canu contig sequences aligned to them with nucmer[25] (MUMmer v3.23). Canu contigs completely spanning artifacts with flanking alignments 10 kb or longer were used to fill hard-masked regions using custom Python scripts (assembly-patch-finder and assembly-patch-patcher, https://bitbucket.org/bredeson/artisanal).

Long-read, signal-based polishing was performed twice on the final scaffolds with the Arrow[21] polishing tool from the SMRTlink (v5.0.0.6792) software suite. BLASR[22] (v5.3) was used to perform the read alignments (--algorithm arrow --fancyChunking --skipUnrecognizedContigs --noEvidenceConsensusCall lowercasereference --refineDinucleotideRepeats --reportEffectiveCoverage).

Illumina-based polishing was performed using the JBL052 2×251 bp WGS sequences and Pilon[35] (v1.23; --fix all --minqual 20 --mindepth 10 --minmq 20 --diploid). Manual inspection of read alignments to the Pilon-polished scaffolds, however, revealed errors persisted (24.4/Mb) in the assembly. This prompted us to devise and implement the following custom polishing procedure: Reads were aligned using BWA-MEM[33] (v0.7.17-r1188) and filtered for proper-pairing using SAMtools[36] (v1.6). Variants were then called using FreeBayes[37] (v1.1.0-54-g49413aa; --use-mapping-quality --genotype-qualities --report-genotype-likelihood-max --min-alternate-count 2 --min-alternate-fraction 0.05 --min-base-quality 20 --min-mapping-quality 20 --min-repeat-entropy 1 --max-complex-gap -1 --strict-vcf) and homozygous variants (SNVs and indels) within the depth range 10–80× were selected. Furthermore, because the predominant error type inherent to long-read sequencing data are indels, and because the Arrow consensus algorithm does not fully support diploid datasets, heterozygous multi-nucleotide indel variants in the depth range 10–80× were also selected to allow incorporating heterozygous loci that may have been excluded by the Arrow consensus procedure. These selected variants were patched into the contigs using a custom Python script called ILEC (v0.1.3; https://bitbucket.org/rokhsar-lab/map4cns). This procedure was iterated (6 times) until the number of frameshift errors observed in the coding regions converged. The number of variants plateaued.

#### Evaluating assembly completeness and correctness

The fraction of Illumina reads ($n$ = 385,527,636) mapped 99.74% total aligned, 96.57% properly paired (**Supplementary Table 1**). The sequence length placed in chromosomes (1,449,319,640 bp) is 99.86% of the assembly, while the remaining unplaced sequences (1,963,959 bp) are distributed in 156 scaffolds. The assembly comparisons between the v9 sequence[1] and this new *X. tropicalis* v10 are shown in **Supplementary Tables 1** and **2**.

The comparison of the mapping of the 52,323 full length *X. tropicalis* transcript predictions show that the current version recovered the sequences of genes that were missing from the previous assembly (**Supplementary Fig. 1E–G**, **Supplementary Table 2**). To determine the genic completeness of this assembly we utilized two sets of *X. tropicalis* transcriptomes obtained from Sanger data and independent from other genome annotations. We aligned the 8,558 full-insert cDNA sequences from the Mammalian Gene Collection (which, despite its name, includes non-mammalian species) using exonerate v2.4.0 (model: coding2genome) (**Supplementary Table**

**2**). A total of 119 frameshifts were identified in 60 cDNA alignments that spanned 11,423,049 bp. Further inspection of the sequences with indels revealed that the significant majority of these were caused by Sanger sequencing errors, rather than errors in the assembly. The second estimate of gene set completeness was assessed by BUSCO[10–12]. Although the BUSCO completeness estimate for the genome sequence is low (91.7%), we find 99.4% of known *X. tropicalis* genes, from the Mammalian Gene Collection[38], are present in the assembly and 97.0% of BUSCOs among its predicted proteins, attesting to its substantial completeness (**Supplementary Table 2**).

Two examples of fixed gene annotation predictions with respect to the previous assembly are *dnai1* and *atp4a* (**Supplementary Fig. 1F–G**). In these cases, some of the exons of the genes lacked Sanger read support and were surrounded by tandem repeats. One case of broken gene prediction corresponds to *myod1,* a locus also surrounded by large stretches of tandem repeats and low-quality sequence (**Supplementary Fig. 1H).** These artifacts were not able to be resolved, possibly due to the limitation of library inset size and the high complexity of the region caused by tandem arrays. In the current assembly most of the gaps are in the subtelomeres (**Supplementary Fig. 3**), as these are packed with larger tandem arrays and have a high GC-content compared to the inner portions of the chromosomes (**Supplementary Fig. 1E**).

#### Estimating the genome size and residual heterozygosity of *Xenopus tropicalis*

Prior to estimating the genome size of *X. tropicalis* using a *k*-mer counting approach, positively identified contaminants were removed from the JBL052 Illumina shotgun sequencing reads. To reduce the amount of sequence to search databases for contaminants, Illumina reads were first aligned to the v10 genome sequence. Read pairs that did not map properly or did not map at all against the nuclear genome were aligned against 1) the complete *X. tropicalis* mitochondrial genome and 2) the NCBI nt database (downloaded 2019-01-22) using BLASTN[39] (v2.9.0; -num_threads 4 -evalue 1e-3 -num_alignments 1). Pairs with at least one read with a best hit to a non-vertebrate species or to the mitochondrial genome were discarded (129,575 pairs). Likewise, any pair with at least one read aligned for 90% of its length at 99% identity to a non-*Xenopus* vertebrate sequence was also discarded (627 pairs). Counted *k*-mers reached a maximum frequency of one million.

Non-contaminant reads were then *k*-mer counted canonically using jellyfish[40] (v2.2.10); where *k* = 51. The resulting 51-mer histogram was analyzed to estimate the genome size with GenomeScope[41] (v1.0.0-6-gd2aefdd). The genome size of *X. tropicalis* was estimated to be 1.452 Gb, with 0.336 Gb (23.1%) of that determined to be captured in repetitive elements (**Supplementary Fig. 1D**).

The reference individual used for sequencing was an $F_{17}$ female derived from the same Nigerian line used to produce the previous v4 and v9 assemblies[1,42]. To assess the degree of residual heterozygosity potentially latent in the inbred reference strain, we used BWA-MEM[33] v0.7.17-r1188 to align the JBL052 2×251 bp paired-end whole-genome shotgun sequencing reads to the v10 assembly, then filtered for proper read-pairing (pair orientation and distance) and variants called with FreeBayes[37] (v1.1.0-54-g49413aa; parameters: --genotype-qualities --use-mapping-quality --report-genotype-likelihood-max --min-base-quality 20 --min-mapping-quality 30 --haplotype-length 1 --use-best-n-alleles 4 --strict-vcf --report-monomorphic). Multi-allelic variants and indels were discarded, and bi-alleleic SNVs were filtered for depth of coverage between $\mu \pm 1.78\sigma$ and allele balance between 0.3 and 0.7, inclusive, to mitigate false positive variant calls caused by low mappability and/or genomic repeats. Rates of heterozygous loci and loci homozygous for non-reference alleles were estimated using 500-kb sliding windows every

50 kb[43] along each chromosome. This approach reveals that this $F_{17}$ Nigerian strain female retains blocks of heterozygosity in 14.1% of its genome (**Supplementary Fig. 6**). The average heterozygosity within these blocks is estimated as $2.96 \times 10^{-3}$, consistent with previous estimates within this species[44].

#### Repeat annotation and estimation of sequence divergence

RepeatModeler[45] v1.0.11 was run on an intermediate, unpolished version of the assembled *X. tropicalis* contigs. Identified repeats were manually curated to exclude false positives and recover false negatives, leading to a total of 973 repeats. Repeats from seven other anuran genome assemblies (*A. truei* [*n* = 1,769], *E. coqui* [*n* = 1,441], *E. pustulosus* [*n* = 1,146], *H. boettgeri* [*n* = 1,160], *P. adspersus* [*n* = 908], *X. borealis* [*n* = 1,026], and *X. laevis* [*n* = 913]) were also identified and included into a pan-anuran repeat library. We combined the frog and ancestral RepBase[46] v23.12 dataset (*n* = 934) with the curated repeats above to create the final repeat library (*n* = 10,270). The curated repeat library was subsequently utilized as input for RepeatMasker[47] v4.0.7 to annotate repeats on *Xenopus tropicalis* v10. A corrected measurement of sequence divergence was obtained by applying the Jukes-Cantor model to the substitution rate of the sequence calculated by RepeatMasker with respect to the consensus repeat family obtained by RepeatModeler and the RepBase database. The repeat annotation using RepeatMasker found that 40.12% of the *X. tropicalis* genome is covered by repeats (**Supplementary Table 4**). Tandem Repeats Finder[48] v4.09 was used to identify all tandem repeats in the chromosomes (**Methods**). A total of 154.75 Mb in the genome are covered by tandem arrays consisting of > 5 consecutive monomers that are greater than 10 bp long (**Supplementary Fig. 3**).

#### Selection of ESTs and valid transcript models for genome annotation

We obtained 1,271,375 *Xenopus tropicalis* EST sequences from NCBI. ESTs sequenced from the forward and reverse orientations were assembled using PEAR[49] v0.9.8. Successfully assembled ESTs and single ESTs with a minimum sequence length of 250 bp were mapped against the v10 genome assembly using STARlong[50] v2.7.0e. Stranded RNA-seq data from adult tissues[51] and non-stranded RNA-seq data from different developmental stages[52] were obtained from the Sequence Read Archive (SRA). We only considered samples that were poly-A selected. We pooled samples from equivalent stage or tissue type to increase sequencing read depth (9 stages: cleavage, blastula, gastrula_early, gastrula_late, neurula_early, neurula_mid, neurula_late, tadpole_early, tadpole_mid, tadpole_late; 6 adult tissues: heart, testis, brain, liver, ovary, kidney. **Supplementary Data 1**). The RNA-seq reads were aligned against the unmasked version of the *Xenopus tropicalis* genome assembly v10 using STAR[50] v2.7.0e. Alignments became the inputs for the Trinity transcriptome assembler[53,54] v2.5.1. We obtained a total of 2.4 billion transcript models, which were subsequently evaluated after being mapped to the genome using STARlong. The EST and transcript model predictions mapped by STARlong were used to evaluate the quality of the transcript models. To prevent fused transcript models, we required that the read coverage spanning splice junctions remained consistent along the transcript. Transcripts and ESTs were also discarded if they presented abnormal mapping features (e.g., low mapping quality, short intron lengths, short first or last exon lengths).

#### Protein coding gene annotation

*X. tropicalis* clones (*n* = 8,980) from the Mammalian Gene Collection[38] (MGC) were used as the set of confident and full-length mRNA evidence. MGC clones from *X. laevis* (*n* = 11,515) were utilized as sister transcripts. Human, mouse, chicken, and zebrafish proteins were used for peptide homology evidence. Several rounds of genome annotation and evaluation were implemented to assess the completeness of the gene predictions. Filtered EST and transcript models were used as mRNA evidence by the JGI Integrated Gene Call[13] (IGC) pipeline for genome annotation (**Supplementary Table 3**).

### Supplementary Note 3: Additional chromosome-scale frog assemblies

#### Genome and transcriptome sequencing

All sequencing data have been deposited in the NCBI SRA and are summarized in **Supplementary Data 1**.

#### DNA extraction and sequencing of *E. pustulosus*

High molecular weight DNA was extracted, as previously described[4], from whole blood from two sisters (237g6f4 and 237g6f5) maintained at the University of the Pacific. Using DNA from one sister (237g6f4), a 10x Genomics Chromium Genome library[23] was prepared and sequenced on the Illumina HiSeq X by the HudsonAlpha Institute for Biotechnology. Using DNA from the other sister (237g6f5), Pacific Biosciences SMRT libraries were prepared and sequenced on the Pacific Biosciences Sequel by the DNA Technologies and Expression Analysis Cores at the

University of California Davis Genome Center. Using liver dissected from a niece of the sisters (291g2f_3603 also coded 291g2f3), also maintained at the University of the Pacific, a HiC library was prepared using the Dovetail Genomics HiC library preparation kit and sequenced on the Illumina HiSeq 4000 by the VCGSL. Two additional HiC libraries were prepared from the dissected liver and sequenced on the Illumina NextSeq by Dovetail Genomics.

#### DNA extraction and sequencing of *H. boettgeri*

High molecular weight DNA was extracted, as previously described[4], from whole blood from one female ($F_2$) purchased at the Albany Aquarium. A 10x Genomics Chromium Genome library[23] was prepared and sequenced on the Illumina HiSeq X by the HudsonAlpha Institute for Biotechnology. Pacific Biosciences SMRT libraries were prepared and sequenced on the Pacific Biosciences Sequel by the HudsonAlpha Institute for Biotechnology. Using liver dissected from a second, unrelated female ($F_3$) purchased at the Albany Aquarium, a HiC library was prepared using the Dovetail Genomics HiC library preparation kit and sequenced on the Illumina HiSeq 4000 by the VCGSL.

#### Additional sequencing of *H. boettgeri*

DNA was extracted, as previously described[4], from whole blood from a female ($F_1$) purchased at the Albany Aquarium. A short insert library was prepared using the Takara PrepX DNA Library Kit by the Functional Genomics Laboratory at the University of California Berkeley and sequenced on the Illumina HiSeq 2500 by the VCGSL.

#### DNA extraction and sequencing of *E. coqui*

Kidney and liver tissue were dissected from one male collected in Hawaii (HN-11 male), and DNA was extracted from these tissues using the Zymo Research Quick gDNA MiniPrep Kit (cat# D3007). Two short insert libraries were prepared using the Takara PrepX DNA Library Kit with the kidney DNA by the Functional Genomics Laboratory at the University of California Berkeley and sequenced on the Illumina HiSeq 2500 and 4000 by the VCGSL. Two mate pair libraries were prepared using liver DNA and sequenced on the Illumina HiSeq 2500 by the HudsonAlpha Institute for Biotechnology. Using the liver tissue sample, a HiC library was prepared and sequenced on the Illumina NextSeq by Dovetail Genomics. High molecular weight DNA was extracted, as previously described[4], from whole blood from a second, unrelated male maintained at Harvard University (C4M). Using DNA from this second male, a 10x Genomics Chromium Genome library[23] was prepared and sequenced on the Illumina HiSeq X by the HudsonAlpha Institute for Biotechnology.

#### Additional sequencing of *E. coqui*

Liver tissue was dissected from one female collected in Hawaii (HN-13 female), and DNA was extracted from the tissue using the Zymo Research Quick gDNA MiniPrep kit (cat# D3007). Two short insert libraries were prepared using the Takara PrepX DNA Library Kit by the Functional Genomics Laboratory at the University of California Berkeley and sequenced on the Illumina HiSeq 2500 and 4000 by the VCGSL.

#### RNA extraction and sequencing of *E. pustulosus*

In addition to the two whole tadpoles (excluding gut) at approximated stages 45 and 56, the following tissues were dissected from adult frogs maintained at the University of the Pacific:

brain (*n* = 3), dorsal skin (*n* = 2), eggs (*n* = 2), eye (*n* = 2), heart (*n* = 2), intestine (*n* = 2), larynx (*n* = 3), liver (*n* = 2), lung (*n* = 2), and ventral skin (*n* = 2). All samples were washed twice with PBS, homogenized in TRIzol Reagent, and centrifuged, followed by flash freezing of the supernatant. RNA was isolated following the *TRIzol Reagent User Guide* (Pub. No. MAN0001271 Rev. A.0) protocol. Illumina mRNA libraries were prepared using the Illumina TruSeq Stranded mRNA Library Prep Kit and sequenced on the Illumina HiSeq 4000 by the VCGSL.

##### RNA extraction and sequencing of *H. boettgeri*

Eggs were homogenized in TRIzol Reagent and processed according to manufacturer's instructions. RNA was then isolated using the QIAGEN RNeasy Mini Kit (cat# 74104). An Illumina mRNA library was prepared using the Takara PrepX RNA-Seq for Illumina Library Kit by the Functional Genomics Laboratory at the University of California Berkeley and sequenced on the Illumina HiSeq 4000 by the VCGSL.

#### Genome assembly

The assembly of *E. coqui, E. pustulosus*, and *H. boettgeri*, detailed below, followed a hierarchical strategy, starting with short-read data and progressing to longer reads and linkages. Hi-C sequence libraries were prepared and sequenced as described in Online Methods.

##### Shotgun assembly of *E. pustulosus*

10x Genomics linked reads were assembled with Supernova[23] (v2.0.1). As previously described[55], putative archaeal, bacterial, viral, and vector contaminants were identified and removed by querying the assembly using BLAST+[39] (v2.6.0) against the respective RefSeq and UniVec databases, using general_decon.sh[55] (v1.0). Putative mitochondrial sequence was also identified and removed by querying the assembly using BLAST+ (v2.6.0) against the closest available mitochondrial assembly[56] (NCBI JX564888.1), using mt_decon.sh[55] (v1.0). Finally, putative nonvertebrate contamination was identified and removed through two rounds of filtering, using custom script nt_decon.sh (v1.0; https://github.com/abmudd/Assembly): (1) the assembly was queried using BLAST+ (v2.6.0) against the NCBI NT database, flagging sequences with an E-value less than $1\times10^{-10}$ best hit to a nonvertebrate sequence, as identified by the corresponding taxonomic information; (2) flagged sequences were queried using BLAST+ (v2.6.0) against previously published frog genomes (*Hyla arborea*[57], *Nanorana parkeri*[58,59], *P. adspersus*[60] (v29Jun2017), *Rana catesbeiana*[61] (v3-20170621), *R. temporaria*[62,63], *X. laevis*[4] (GCA_001663975.1), and *X. tropicalis*[1] (GCA_000004195.3)) as well as frog sequences from NCBI EST, GSS, and nucleotide databases, removing sequences without any hits based on a cutoff of 75% identity and an E-value less than $1\times10^{-10}$. The decontamination removed 8,581 scaffolds totaling 2.11 Mb from the Supernova assembly.

#### Initial PacBio scaffolding of *E. pustulosus*

To improve the contiguity of the *E. pustulosus* assembly, decontaminated Supernova contigs were scaffolded with PacBio long-read data. This was achieved by performing a hybrid assembly of the filtered Supernova contigs and PacBio long reads using DBG2OLC[20] (commit 1f7e752). PacBio reads were then mapped to the DBG2OLC output assembly with BLASR[22] (commit 4323a52) and polished with PBDAGCON[21] (commit 1a2f1e7) two times, using the map4cns pipeline (commit dd89f52; https://bitbucket.org/rokhsar-lab/map4cns). Decontaminated Supernova contigs were mapped back to the polished DBG2OLC assembly using MUMmer[25] (v3.23) with a cutoff of 90% identity and then ordered and oriented into scaffolds using tsvtk and maptk from the GBS analysis pipeline[64] (commit 80613d5).

#### Initial chromosome assembly of *E. pustulosus*

After PacBio-based long-read scaffolding, the assembly was organized into chromosomes with HiC data using the Dovetail Genomics HiRise pipeline[65]. HiC reads were then aligned to the assembly with Juicer[26] (commit d3ee11b), and the assembly was manually corrected in Juicebox[28] (v1.9.0) with Juicebox Assembly Tools[29]. PacBio reads were aligned to the assembly with BWA[33] (v0.7.17-r1188), and gaps were resized using scripts pbGapLen and expand-gaps.py (https://bitbucket.org/bredeson/artisanal). Gaps in the assembly were then filled with PacBio data using PBJelly[34] (PBSuite v15.8.24).

#### Revised PacBio and 10x Genomics scaffolding of *E. pustulosus*

Given the limited resolution of the *E. pustulosus* HiC data for determining the proper order and orientation of scaffolds as well as the large number of gaps listed as overfilled by PBJelly—suggesting incorrect scaffolding, rearrangements, or other assembly errors—the gap-filled assembly was broken into contigs and then scaffolded with PacBio and 10x Genomics data. First, the contigs were scaffolded against the error corrected DBG2OLC assembly using MUMmer[25] (v3.23) as well as tsvtk and maptk in the GBS analysis pipeline[64] (commit 80613d5). Next, the assembly was scaffolded with the 10x Genomics linked reads using Scaff10X (v2.1; https://sourceforge.net/projects/phusion2/files/scaff10x). Gaps were resized with PacBio data, as previously described, and filled with PBJelly[34] (PBSuite v15.8.24).

#### Revised chromosome assembly of *E. pustulosus*

The resulting assembly was organized back into chromosomes based on alignment against the initial chromosome assembly using MUMmer[25] (v3.23) as well as tsvtk and maptk scripts in the GBS analysis pipeline[64] (commit 80613d5). Gaps were again resized with PacBio data using pbGapLen and expand-gaps.py (https://bitbucket.org/bredeson/artisanal), and gaps were filled with PBJelly[34] (PBSuite v15.8.24).

#### Final assembly correction of *E. pustulosus*

The assembly was polished with two rounds of Illumina error correction. In this, 10x Genomics data were adapter trimmed using trim_10X.py[55] (v1.0) and aligned to the assembly with BWA[33] (v0.7.17-r1188). Variants called by FreeBayes[37] (commit 49413aa) with a read depth within two standard deviations of the Gaussian fit (mean of 26.4 and standard deviation of 10.4) were corrected using the script ILEC in the map4cns pipeline (commit dd89f52; https://bitbucket.org/rokhsar-lab/map4cns).

After error correction, the HiC data were realigned to the assembly with Juicer[26] (commit d3ee11b). Misjoins were identified and broken in Juicebox[28] (v1.9.0) with Juicebox Assembly Tools[29]. Remaining gaps were resized with the PacBio data using BWA[33] (v0.7.17-r1188), pbGapLen, and expand-gaps.py, and closure was attempted with the adapter-trimmed 10x Genomics data using Platanus[66] (v1.2.1).

#### Final assembly release of *E. pustulosus*

Scaffolds smaller than one kb were removed from the final assembly with seqtk (v1.3-r106; https://github.com/lh3/seqtk), and chromosomes and scaffolds were numbered in order of size using SeqKit[67] (v0.7.2-dev). Chromosomes were oriented arbitrarily.

#### Assembly of *H. boettgeri*

The assembly process followed the same procedure as outlined for *E. pustulosus* with four differences: the closest available mitochondrial assembly[68] used in decontamination was NCBI NC_015615.1; the decontamination removed 108 scaffolds totaling 39.9 kb from the Supernova assembly; variants with a read depth within only one standard deviation of the Gaussian fit (mean of 18.4 and standard deviation of 12.3) were corrected; and chromosomes were numbered and oriented based on alignment with MUMmer[25] (v3.23) to the *X. tropicalis* chromosomes.

#### Shotgun assembly of *E. coqui*

The short insert libraries were adapter trimmed with ea-utils fastq-mcf[69] (commit bd148d4). The mate pair libraries were adapter trimmed and split with NxTrim[70] (commit 53c2193). Using custom script nxtrim_pipeline.sh (v1.0; https://github.com/abmudd/Assembly), the output from NxTrim was divided into two files: (1) reads flagged as mate pair or unknown were merged into a final mate pair file; (2) reads flagged as short insert paired-end or single-end were merged into a final short insert library, with the single-end reads given a corresponding blank second end. All trimmed data was then assembled with Meraculous[71,72] (v2.2.4). Mitochondrial sequence was assembled from adapter-trimmed short insert data using custom script organelle_pipeline.py (v1.0; https://github.com/abmudd/Assembly) and NOVOPlasty[73] (v2.6.3), with other Hyloidea mitochondrial assemblies available on NCBI as input seeds.

Mirroring the *E. pustulosus* shotgun assembly process and as previously described[55], putative archaeal, bacterial, viral, and vector contamination was identified and removed by querying the assembly with BLAST+[39] (v2.3.0) against the respective RefSeq and UniVec databases, using general_decon.sh[55] (v1.0). Putative mitochondrial sequence was also identified and removed by querying the assembly with BLAST+ (v2.3.0) against the assembled mitochondrial sequence, using mt_decon.sh[55] (v1.0). In addition, the assembly was queried with BLAST+ (v2.3.0) against the NT database, previously published frog genomes (*H. arborea*[57], *N. parkeri*[58,59], *P. adspersus*[60], (v29Jun2017), *R. catesbeiana*[61] (v3-20170621), *R. temporaria*[62,63], *X. laevis*[4] (GCA_001663975.1), and *X. tropicalis*[1] (GCA_000004195.3)), and frog sequences from NCBI EST, GSS, and nucleotide databases to identify and remove non-vertebrate sequences, using custom script nt_decon.sh (v1.0; https://github.com/abmudd/Assembly). The decontamination removed 7,272 scaffolds totaling 2.06 Mb from the meraculous assembly.

Residual redundancy due to split haplotypes was identified and removed using custom script align_pipeline.sh (v1.0; https://github.com/abmudd/Assembly). To summarize, the adapter-trimmed libraries were aligned to the assembly with BWA[33] (v0.7.15-r1140). Read depth was extracted from the alignments and used as a cutoff to separate half-depth and full-depth scaffolds. Half-depth scaffolds were then queried against each other with BLAST+ (v2.3.0), and the smaller of each best-hit scaffold pair was extracted. Half-depth scaffolds were also *k*-mer counted with Jellyfish[40] (v2.1.4), and the smaller of each scaffold pair with a unique, shared 31-mer was extracted. Scaffolds identified in both BLAST+ and Jellyfish analyses were removed from the assembly. The redundancy pipeline removed 192,996 scaffolds totaling 31.1 Mb from the decontaminated assembly. The assembly was then scaffolded with SSPACE[32] (v3.0).

#### Chromosome assembly of *E. coqui*

The assembly was next scaffolded with 10x Genomics linked reads using Scaff10X (v2.1; https://sourceforge.net/projects/phusion2/files/scaff10x). Attempts to further scaffold the assembly into chromosomes with the HiC data using the Dovetail Genomics HiRise pipeline[65] and 3D-DNA[27] (commit 745779b) were unsuccessful. Therefore, the Scaff10X output was mapped to the *E. pustulosus* assembly using MUMmer[25] (v3.23) and then scaffolded based on synteny using tsvtk and maptk from the GBS analysis pipeline[64] (commit 80613d5). HiC reads were aligned to the assembly with Juicer[26] (commit d3ee11b), and the synteny-based scaffolding was manually corrected in Juicebox[28] (v1.9.0) with Juicebox Assembly Tools[29]. Closure of the remaining gaps was attempted with the adapter-trimmed short insert data using Platanus[66] (v1.2.1).

#### Final assembly release of *E. coqui*

Scaffolds smaller than one kb were removed from the final assembly with seqtk (v1.3-r106; https://github.com/lh3/seqtk), and chromosomes and scaffolds were numbered in order of size using SeqKit[67] (v0.7.2-dev). Chromosomes were oriented arbitrarily.

### Correction of published genomes

#### Reassembly of *L. ailaonicum*

As previously described for *E. pustulosus*, putative archaeal, bacterial, viral, and vector contamination was checked by querying the published assembly[74,75] using BLAST+[39] (v2.9.0) against the respective RefSeq and UniVec databases. Putative mitochondrial sequence was also checked by querying the assembly using BLAST+ (v2.9.0) against the closest available mitochondrial assembly[76] (NCBI NC_024427.1). No contaminant scaffolds were identified or removed from the assembly.

HiC reads[74] (BioProject PRJNA523649) were aligned to the assembly with Juicer[26] (commit d3ee11b), and existing scaffolding was manually error corrected in Juicebox[28] (v1.11.08) with Juicebox Assembly Tools[29]. All gaps were then resized to 100 bp. Chromosomes and scaffolds were numbered in order of size using SeqKit[67] (v0.7.2-dev). Chromosomes were oriented arbitrarily.

#### Reassembly of *P. adspersus*

As previously described for *E. pustulosus*, putative archaeal, bacterial, viral, and vector contamination was identified and removed by querying a precursor (v29Jun2017) of the released assembly[60] (GCA_004786255.1) using BLAST+ (v2.6.0) against the respective RefSeq and UniVec databases. Putative mitochondrial sequence was also identified and removed by querying the assembly using BLAST+ (v2.6.0) against the closest available mitochondrial assembly[56] (NCBI JX564898.1). The decontamination removed one scaffold totaling 1.45 kb from the assembly.

Chicago and HiC reads[60] (BioProject PRJNA439445) were aligned to the assembly with Juicer[26] (commit d3ee11b), and existing scaffolding was manually error corrected in Juicebox[28] (v1.9.0) with Juicebox Assembly Tools[29]. PacBio reads[60] (BioProject PRJNA439445) were aligned to the assembly with BWA[33] (v0.7.17-r1188), and gaps were resized using scripts pbGapLen and expand-gaps.py (https://bitbucket.org/bredeson/artisanal). Closure of the remaining gaps was attempted with Platanus[66] (v1.2.1) using TruSeq data[60] (BioProject PRJNA439445) adapter-trimmed with ea-utils fastq-mcf[69] (commit bd148d4).

Scaffolds smaller than one kb were removed from the final assembly with seqtk (v1.3-r106; https://github.com/lh3/seqtk), and chromosomes and scaffolds were numbered in order of size using SeqKit[67] (v0.7.2-dev). Chromosomes were later renamed and reoriented based on alignment with MUMmer[25] (v3.23) to the released assembly[60] (GCA_004786255.1).

#### Protein-coding gene annotation

Using the previously described final repeat library (*n* = 10,270; **Supplementary Note 2**), which combined the pan-anuran repeat library and ancestral RepBase dataset, the chromosome-scale assemblies of *E. coqui*, *E. pustulosus*, *H. boettgeri*, *L. ailaonicum*, and *P. adspersus* were soft masked with RepeatMasker[45,47] (v4.0.7 and v4.0.9). In addition to the RNA sequencing above, additional RNA data for *H. boettgeri*[4] (BioProject PRJNA306175) and *P. adspersus*[60] (BioProject PRJNA439445) were downloaded from NCBI SRA. Unpublished *E. coqui* RNA sequencing of stages 7, 10, and 13 hindlimb and stage 9–10 tail fin skin was obtained from Harvard University and the French National Center for Scientific Research, respectively. All RNA sequencing data was adapter trimmed with ea-utils fastq-mcf[69] (commit bd148d4) and aligned to the respective assemblies with STAR[50] (v2.5.3a and v2.7.0f), using the custom script STARalign.sh (v1.0; https://github.com/abmudd/Assembly).

Genome-guided transcriptomes were assembled with Trinity[53,54] (v2.5.1) for each individual RNA library: *E. coqui* (*n* = 7), *E. pustulosus* (*n* = 24), *H. boettgeri* (*n* = 9), and *P. adspersus* (*n* = 2). The assembled transcriptomes for these four species were aligned to the respective assemblies with STARlong (v2.7.1a) and then split into single-exon and multi-exon transcripts based on their alignments, using custom script filter_trinity.py (v1.0; https://github.com/abmudd/Assembly). Multi-exon transcripts were discarded if the first exon and/or last exon was less than 60 bp in length, if an intron was less than 60 bp or greater than 300,000 bp in length, or if the total transcript alignment length was less than 250 bp. Single-exon transcripts larger than 80 amino acids and containing start and stop codons were extracted with TransDecoder[54] (v3.0.1).

Filtered single-exon and multi-exon transcripts were merged and used as mRNA evidence in the IGC pipeline[13] for genome annotation of the four species. Peptides from the *X. tropicalis* annotation (**Supplementary Note 2**) as well as SwissProt eukaryotes[77] (downloaded November 2018) were used as protein homology evidence in this pipeline.

### Supplementary Note 4: Comparative analysis

#### Comparative analysis

##### Gene homology and orthology

From the resulting annotations, gene homology between *E. coqui*, *E. pustulosus*, *H. boettgeri*, *L. ailaonicum*[74,75], *P. adspersus*, *X. laevis*[4] (v9), and *X. tropicalis* was analyzed with OrthoVenn2 (ref.[2]) using an E-value of $1\times10^{-5}$ and an inflation value of 1.5. One-to-one gene orthologs between *E. coqui*, *E. pustulosus*, *H. boettgeri*, *P. adspersus*, and *X. tropicalis* were extracted from the OrthoVenn2 output, after requiring the ortholog sets to be either present in a single copy or absent in *L. ailaonicum*[74,75] and the L and S subgenomes of *X. laevis*[4] (v9). Regarding the exceptions for *L. ailaonicum* and *X. laevis*, an initial analysis of the OrthoVenn2 output found that the *L. ailaonicum* annotation[74,75] was missing 37% (3,670) of the one-to-one gene orthologs found in the five main frog species, whereas the *X. laevis* annotation[4] (v9) has confounding factors associated with the allopolyploidization and resulting gene evolution. In constructing this set, we also excluded genes on the *P. adspersus* W chromosome.

#### Whole-genome multiple alignment

The assemblies for *Ambystoma mexicanum*[78,79] (GCA_002915635.2), *E. coqui*, *E. pustulosus*, *H. boettgeri*, *L. ailaonicum*, and *P. adspersus* were each aligned pairwise against *X. tropicalis* with cactus[80] (commit e4d0859). *X. laevis*[4] (v9) was broken into subgenomes, and the chromosomes of each subgenome were aligned against *X. tropicalis* with cactus (commit e4d0859). As previously described[55], all pairwise output HAL alignment files were filtered and converted into MAF, using cactus_filter.py[55] (v1.0), and runs of collinearity were extracted from each pairwise MAF file. The pairwise MAF files were also merged with ROAST/MULTIZ[81] (v012109), using the phylogenetic topology from TimeTree[16], and sorted with last[82] (v979).

#### Phylogeny

Using the 9,624 identified one-to-one orthologous genes and the ROAST-merged MAF file, as previously described[55], fourfold degenerate bases were extracted with script 4Dextract.py[55] (v1.0) and converted into PHYLIP format with BeforePhylo (commit 0885849; https://github.com/qiyunzhu/BeforePhylo). The maximum likelihood tree was estimated with RAxML[14] (v8.2.11) using the GTR+Gamma model of substitution with outgroup *A. mexicanum*.

#### Estimated divergence times

We estimated divergence times from the fourfold synonymous site alignment with MEGA7[17] (v7.0.26), as previously described[83]. The MEGA7 time tree was constructed using the Reltime method[84] with the GTR+Gamma model of substitution. The confidence intervals provided by TimeTree (retrieved on October 31, 2019) for all nodes except the *X. laevis* L – *X. laevis* S node were used as input to MEGA7. These input ranges and output times are noted in **Supplementary Table 11**. The time calculated for the split between L and S subgenomes in *X. laevis* was substantiated in the literature[4,85].

#### Chromosome evolution

As previously described[55], pairwise alignments were extracted from the ROAST-merged MAF file using custom script extract2speciesmaf.py[55] (v1.0) and converted into runs of collinearity following the process used in cactus_filter.py[55] (v1.0). The runs of collinearity were visualized (**Supplementary Fig. 8**) with Circos[86] (v0.69-6) and, following file conversion with custom scripts mcscan_convert_links.py[55] (v1.0) and mcscan_invert_chr.py[55] (v1.0), with jcvi.graphics.karyotype (v0.8.12; https://github.com/tanghaibao/jcvi). The previously extracted one-to-one gene orthologs were similarly visualized with Circos[86] (v0.69-6) and jcvi.graphics.karyotype (v0.8.12; https://github.com/tanghaibao/jcvi). Based on these visualizations and the analyzed phylogeny, with the assumption of the parsimony principle, we extracted chromosome changes using the following logic: changes that were shared in the same order and orientation between two sister species were present in the common ancestor. Any changes that did not meet this criterion were classified as lineage-specific changes.

### Supplementary Note 5: Genome analysis

#### Recombination rate and genomic landscapes

The genetic map was re-made on the final version of the genome (**Methods**). Of the total of 1,277 genetic markers, we considered 1,168 (91.5%) markers whose genetic distance was equal or lower than the distance of the marker immediately adjacent to the right. Recombination rates were calculated using the first derivative of the interpolated genetic distances (**Supplementary Fig. 11**). We observed that the recombination rate correlated the highest with satellite repeats, GC content, and the first principal component obtained from the repeat density matrix (**Supplementary Table 14**).

#### Classification of peri-centromeric and sub-telomeric regions based on repeat content

To determine the sources of variation in repeat content across the genome we used the repeat density matrix as input for the Principal Component Analysis (PCA) (**Methods**). The first 3 principal components described 7.53% of the total variance and the three of them distinguish landmarks of abundant repetitive regions. The first two principal components describe the higher-order structure of the chromosomes (**Supplementary Fig. 5A**). The first principal component (PC1) is correlated with GC% content (Pearson R = +0.796), and recombination rate (Pearson correlation R = +0.77, **Supplementary Fig. 5A**). The second principal component describes the repeats present in the pericentromeric and distal subtelomeric regions. The peaks observed from the second principal component (PC2) at the inner portions of the chromosomes are in close proximity with the HiC estimates of the centromeric positions and surrounding the centromeric repeats. To estimate the centromeric positions based on repeat content, we smoothed the PC2-eigenvectors of each chromosome using the natural cubic spline method (knots = 40). The position of the peak summit, of the highest and widest PC2-smoothed eigenvalues, was considered as the centromeric-estimate of each chromosome (**Supplementary Fig. 5A**). The third principal component captures the variance of repeats associated with A/B compartments (**Supplementary Fig. 5B**). The shifts in the signs of the eigenvectors of the PCA from the HiC coincide with the shift in signs of the PC3 on the repeat density. A moderate correlation with the eigenvalues of the HiC used to define A/B compartments (Pearson R value = –0.44). Harbinger-N9_XT is positively associated with PC3 (R = +0.50), whereas DNA/hAT-Ac appears to strongly correlate with PC3 (R = –0.80).

#### Identification of Tandem Repeats enriched in Centromeric and Subtelomeric regions

Centromeric tandem repeats were identified as described in **Methods**. Pericentromeric tandem repeats shown in **Supplementary Fig. 12A** are enriched 4-fold in the pericentromeric region compared to the rest of the chromosome and have a footprint > 20 Kb inside the pericentromeric region. Subtelomeric enriched repeats were enriched at least 8-fold in subtelomeric regions and a footprint > 50 Kb in the subtelomeres of all metacentric chromosomes (**Supplementary Fig. 12A**, **Supplementary Table 15**). Monomer "a" represented in **Supplementary Fig. 12A** corresponds to the 205-bp consensus monomer (below) that forms part of the centromeric tandem repeat in all chromosomes in *Xenopus tropicalis* (**Supplementary Fig. 10D**). The multiple sequence alignment of the monomeric units span blocks between 41 and 258 kb per chromosome (**Supplementary Table 12**) and exhibit an average sequence identity over 95% at the per-base level (**Supplementary Fig. 10E**).

Some of the tandem repeats (**Supplementary Fig. 12A)** have sequence similarity to consensus repetitive elements (**Supplementary Table 16**). The distribution of the repeats, their length, density, and the JC distance from the consensus sequence indicate that repeats near the subtelomeres are targets of tandem repeat expansion (**Supplementary Fig. 5**).

```
>Xt_CR Xenopus tropicalis centromeric tandem repeat
TTGAATGCTACATGGCATTTGAAAGACAGTGCAGAAAATAAGCTCCTGCTAACGTTTCATGCTTGCAAACCAATCAGAGCACTTG
CAGTAACATGGGCTAAAACGCTTTACAGAGCAAACCGGCAAACTGAAGCAAATCACGCGAATGCCTCAGAAAAGCTCATTTAACA
CACAAAATTGACTATCTATAAGACAAAGTAAATAG
```

#### Centromere inference with HiC

Examining the genome-wide HiC contact matrix revealed "water lily"-shaped inter-chromosomal contact patterns and puncta derived from centromere clustering (**Fig. 1** and **Supplementary Fig. 2**), similar to the Rabl-like patterns described for budding yeast by Duan *et al*.[87]. This motivated estimating the centromeric positions using HiC data (**Supplementary Fig. 10A**). Centurion[88] v0.1.0-3-g985439c was invoked using MapQ ≥ 0 contact maps (for which automated estimates were more stable; **Supplementary Fig. 10B**) at 1 Mb matrix resolution, and with the *coef* parameter set to 10,  to refine initial centromere positions made by eye in Juicebox. All ten estimates localized between the centromere-flanking genes described by Uno *et al*.[89] .

The above procedure was then similarly conducted for *E. coqui*, *E. pustulosus*, *P. adspersus*, *H. boettgeri*, and *X. laevis*. Some of these species, however, have acrocentric or submetacentric chromosomes and Centurion could not reliably estimate the centromere positions for those chromosomes. The centromere positions for these chromosomes, and those with Centurion-estimated ($C_c$) positions differing from their initial estimates ($C_i$) by more than 10% [calculated as ($C_c - C_i$) / max($C_i$, $L - C_i$); where $L$ is the length of the chromosome], were reverted to their initial by-eye estimate; these include *E. coqui* chromosomes 2, 3, 6, 7, and 9–13; *E. pustulosus* chromosomes 8–11; *H. boettgeri* chromosomes 4, 6, 7, and 9; and *P. adspersus* chromosomes 9 and W.

#### Cenp-a binding to CTR-A tandem repeats

ChIP-seq targeting Cenp-a and Histones H3 and H4 was performed as described in **Methods**. Over 97% of all Cenp-a ChIP-seq reads were mapped to the genome (**Supplementary Table 13**). With H3 and H4 ChIP-seq there is an apparent enrichment of reads at the pericentromeric regions, suggesting a ~2–5× assembly collapse in the length of the centromeric repeats. The collapse is likely caused by the lack of variants that would enable the assembly of our PacBio reads across the long centromeric sequence. The ratios between Cenp-a/H4 and Cenp-a/S2 are shown in **Supplementary Fig. 10C** and **Supplementary Table 13**.

#### A/B compartment structure inference using HiC

HiC reads from *X. tropicalis* blood cells were aligned to the v10 assembly with Juicer[26] (v1.5.4-71-gd3ee11b). Normalized (observed over expected, Knight-Ruiz balanced) intra-chromosomal HiC contact matrices were then extracted at 250 kb matrix resolution with Juicer Tools (v1.5.4-71-gd3ee11b) from read pairs with mapping quality of at least 30 (MapQ ≥ 30). An R[90] (v3.5.0) script implementing a sliding-window-based principal component analysis (PCA) algorithm was used to call compartment structure along each chromosome (call-compartments.R, https://bitbucket.org/bredeson/artisanal). Localizing the PCA along the diagonal of the Pearson correlation matrix with sliding windows (here, of width 80 and with step of 40) mitigates confounding signal introduced by intra-chromosomal p-q arm contacts, thereby amplifying compartment signal. **Fig. 5A–B** presents the correlation matrix describing the compartment structure for chromosome 1 and the corresponding eigenvectors obtained from the localized PCA algorithm.

Gene densities (gene count/Mb) were calculated for each compartment bin with bedtools intersect[91] (v2.28.0). The A and B compartment assignments were determined for each chromosome by obtaining the correlation between gene density (genes count/Mb) and the eigenvector of the HiC correlation matrix. If a chromosome exhibited a negative correlation between gene density and the HiC eigenvector, then the sign of the eigenvector for that chromosome was inverted. Compartment A was assigned to bins associated with high gene density. Repeat densities were subsequently calculated for A and B compartments. These above methods were performed for all species included in this study.

#### Identifying extended subtelomere boundaries with HiC

Inter-chromosomal MapQ ≥ 0 and MapQ ≥ 30 HiC contact matrices were extracted at 1 Mb resolution for each chromosome 1-chromosome N pair with Juicer Tools (v1.5.4-71-gd3ee11b). MapQ ≥ 30 observed counts were subtracted from MapQ ≥ 0 observed counts to isolate repeat-specific signals observed in the presumptive subtelomeres (**Fig. 4** and **Supplementary Fig. 2**). We used two methods to extract subtelomere region boundaries from the subtracted matrices: 1) a *k*-means clustering method implementing nine tessellated rectangular Voronoi cells (**Supplementary Fig. 3**) categorized into three classes (corners, center, and edges) constrained to change only their relative dimensions during the optimization procedure, and 2) a principal component analysis (PCA) of the same subtracted matrix (**Supplementary Fig. 3C**).

Computations were performed in R v3.5.0 with the ‘prcomp’ function. See https://bitbucket.org/rokhsar-lab/xentr10/src/master/hic for more details.

#### Dissecting the Rabl chromosome structure

Because A/B compartmentalization (i.e., the checkerboard pattern) and high-density contacts between adjacent genomic loci (i.e., the strong diagonal band) were dominant sources of variance within chromosomes, dense inter-chromosomal contact matrices were used as a proxy instead for visualizing the Rabl structure of chromosomes within the nucleus. We extracted the observed counts of MapQ ≥ 30 contacts at 1 Mb resolution. For each chromosome 1-chromosome N pair, raw matrix counts were smoothed using a radius 5% of each chromosome’s length and taking the median of the sampled values (the median provided more stable smoothing in the presence of extreme-valued outliers, and so was preferred over the arithmetic mean).

To visualize Rabl chromatin structure, we performed PCA with the ‘prcomp’ function in R[90] (v3.5.0) on the smoothed matrix and plotted PC1 vs. PC2 (**Supplementary Fig. 16**). We measured the strength of each chromosome’s Rabl structure as the sum of squared distances (SSD) between chromosome arms (**Supplementary Table 18**) with the expectation that perfectly constrained arms will result in an SSD approaching zero, and unconstrained (i.e., freely-moving) arms taking large values. SSD was computed as the squared Euclidean distances, in PC1-PC2 dimensions, between points $c - i$ and $c + i$ for $i = 1 \rightarrow n$, where $c$ is the index of the 1-Mb window containing the centromere and $n$ is the length of the shortest chromosome arm. The significance $p$-value for each chromosome’s observed Rabl structure was calculated by permutation testing, with variance as the test statistic (alpha = 0.01, one-tailed, 1,000 iterations). Each permutation started from a smoothed matrix, itself derived from a raw-count matrix reordered by 5,000 random row-/column-swap operations, the PCA computed, and the variance of radian angle change (i.e., second derivative) calculated between two vectors created by triplets of consecutively-ordered points along a chromosome. Mathematically, this is calculated as:

$$\mathrm{var}_{i\,=\,1 \rightarrow n-2}\ \theta(v_{i+1}) - \theta(v_i),$$

where

$$\theta = \arccos\,[\ v \cdot u\ /\ |v| \cdot |u|\ ]$$

$$v_{i+1} = (\mathrm{PC1}_{i+2} - \mathrm{PC1}_{i+1},\ \mathrm{PC2}_{i+2} - \mathrm{PC2}_{i+1})$$

$$v_{i-1} = (\mathrm{PC1}_{i+1} - \mathrm{PC1}_{i},\ \mathrm{PC2}_{i+1} - \mathrm{PC2}_{i})$$

$$u = (\mathrm{sign}\ [\mathrm{PC1}_{i+1} - \mathrm{PC1}_{i}],\ 0)$$

Two vectors connecting three consecutively ordered points along a single path introduce relatively little angular change (i.e., little variance), whereas unpatterned (randomly ordered) points tend to induce random directional changes, and greater angular changes (i.e., greater variance), between connected vectors. The R script used to perform the calculations can be found at https://bitbucket.org/rokhsar-lab/xentr10/src/master/hic.

61. Hammond, S. A. *et al.* The North American bullfrog draft genome provides insight into

hormonal regulation of long noncoding RNA. *Nat. Commun.* **8**, 1433 (2017).
62. Palomar, G. *et al.* Comparative high-density linkage mapping reveals conserved genome structure but variation in levels of heterochiasmy and location of recombination cold spots in the common frog. *G3* **7**, 637–645 (2017).
63. Palomar, G. *et al.* Draft genome *Rana temporaria*. (2016) doi:10.6084/m9.figshare.4483439.v1.
64. International Cassava Genetic Map Consortium (ICGMC). High-resolution linkage map and chromosome-scale genome assembly for cassava (*Manihot esculenta* Crantz) from 10 populations. *G3: Genes|Genomes|Genetics* vol. 5 133–144 (2015).
65. Putnam, N. H. *et al.* Chromosome-scale shotgun assembly using an in vitro method for long-range linkage. *Genome Res.* **26**, 342–350 (2016).
66. Kajitani, R. *et al.* Efficient *de novo* assembly of highly heterozygous genomes from whole-genome shotgun short reads. *Genome Res.* **24**, 1384–1395 (2014).
67. Shen, W., Le, S., Li, Y. & Hu, F. SeqKit: A cross-platform and ultrafast toolkit for FASTA/Q file manipulation. *PLoS One* **11**, e0163962 (2016).
68. Irisarri, I., Vences, M., Mauro, D. S., Glaw, F. & Zardoya, R. Reversal to air-driven sound production revealed by a molecular phylogeny of tongueless frogs, family Pipidae. *BMC Evolutionary Biology* vol. 11 (2011).
69. Aronesty, E. Comparison of sequencing utility programs. *TOBIOIJ* **7**, 1–8 (2013).
70. O'Connell, J. *et al.* NxTrim: Optimized trimming of Illumina mate pair reads. *Bioinformatics* **31**, 2035–2037 (2015).
71. Chapman, J. A. *et al.* Meraculous: *De novo* genome assembly with short paired-end reads. *PLoS One* **6**, e23501 (2011).
72. Goltsman, E., Ho, I. & Rokhsar, D. Meraculous-2D: Haplotype-sensitive assembly of highly heterozygous genomes. *arXiv [q-bio.GN]* (2017).
73. Dierckxsens, N., Mardulyn, P. & Smits, G. NOVOPlasty: *de novo* assembly of organelle

genomes from whole genome data. *Nucleic Acids Res.* **45**, e18 (2017).

85. Evans, B. J., Kelley, D. B., Tinsley, R. C., Melnick, D. J. & Cannatella, D. C. A mitochondrial

DNA phylogeny of African clawed frogs: phylogeography and implications for polyploid evolution. *Mol. Phylogenet. Evol.* **33**, 197–213 (2004).

86. Krzywinski, M. *et al.* Circos: an information aesthetic for comparative genomics. *Genome Res.* **19**, 1639–1645 (2009).
87. Duan, Z. *et al.* A three-dimensional model of the yeast genome. *Nature* vol. 465 363–367 (2010).
88. Varoquaux, N. *et al.* Accurate identification of centromere locations in yeast genomes using Hi-C. *Nucleic Acids Res.* **43**, 5331–5339 (2015).
89. Uno, Y., Nishida, C., Takagi, C., Ueno, N. & Matsuda, Y. Homoeologous chromosomes of *Xenopus laevis* are highly conserved after whole-genome duplication. *Heredity* vol. 111 430–436 (2013).
90. R Core Team. R Core Team. R: A language and environment for statistical computing. *Foundation for Statistical Computing* (2013).
91. Quinlan, A. R. BEDTools: The Swiss-army tool for genome feature analysis. *Curr. Protoc. Bioinformatics* **47**, 11.12.1–34 (2014).
